# Identifying zoonotic origin of SARS-CoV-2 by modeling the binding affinity between Spike receptor-binding domain and host ACE2

**DOI:** 10.1101/2020.09.11.293449

**Authors:** Xiaoqiang Huang, Chengxin Zhang, Robin Pearce, Gilbert S. Omenn, Yang Zhang

## Abstract

Despite considerable research progress on SARS-CoV-2, the direct zoonotic origin (intermediate host) of the virus remains ambiguous. The most definitive approach to identify the intermediate host would be the detection of SARS-CoV-2-like coronaviruses in wild animals. However, due to the high number of animal species, it is not feasible to screen all the species in the laboratory. Given that the recognition of the binding ACE2 proteins is the first step for the coronaviruses to invade host cells, we proposed a computational pipeline to identify potential intermediate hosts of SARS-CoV-2 by modeling the binding affinity between the Spike receptor-binding domain (RBD) and host ACE2. Using this pipeline, we systematically examined 285 ACE2 variants from mammals, birds, fish, reptiles, and amphibians, and found that the binding energies calculated on the modeled Spike-RBD/ACE2 complex structures correlate closely with the effectiveness of animal infections as determined by multiple experimental datasets. Built on the optimized binding affinity cutoff, we suggested a set of 96 mammals, including 48 experimentally investigated ones, which are permissive to SARS-CoV-2, with candidates from primates, rodents, and carnivores at the highest risk of infection. Overall, this work not only suggested a limited range of potential intermediate SARS-CoV-2 hosts for further experimental investigation; but more importantly, it proposed a new structure-based approach to general zoonotic origin and susceptibility analyses that are critical for human infectious disease control and wildlife protection.

## Introduction

Identification of the direct zoonotic origin (intermediate host) of severe acute respiratory syndrome coronavirus 2 (SARS-CoV-2) is important for combating the coronavirus disease 2019 (COVID-19) pandemic [1, 2]. It has become well accepted that SARS-CoV-2 was likely to originate naturally from bats soon after its outbreak, built on the fact that SARS-CoV-2 shares a 96.2% nucleotide sequence identity with the bat coronavirus (CoV) RaTG13 isolated from *Rhinolophus affinis* [3] and that natural insertions were identified at the S1/S2 cleavage site of the Spike (S) protein of RmYN02-CoV isolated from *Rhinolophus malayanus* [4]. However, it remains unknown how the related CoV was transmitted from bats to humans.

*In vitro* experiments suggest that RaTG13 also binds to human ACE2 (hACE2) and can use hACE2 as an entry receptor [5]; thus, it could be possible that a progenitor of SARS-CoV-2, e.g. RaTG13 or RaTG13-like CoV, infected humans and evolved during human-to-human transmission [6]. However, recent experiments show that the binding efficiency of RaTG13 to hACE2 is quite low [7], probably due to the lack of critical hACE2-binding residues. Besides, no evidence has shown that RaTG13 can directly infect humans in nature.

It is widely believed that the novel CoV was transmitted from its natural host to humans via some intermediate host, during which a progenitor of SARS-CoV-2 acquired the critical ACE2 binding residues and/or furin cleavage site [6]. This point of view is supported in part by the fact that pangolin-CoV isolated from *Manis javanica* shares almost identical key ACE2-binding residues with SARS-CoV-2 [8-11]. However, it is controversial whether pangolins are the intermediate host [9, 10] or natural host [8, 11], or whether they are a host [12, 13]. Phylogenetic analyses show that some pangolin-CoVs are genetically related to SARS-CoV-2 but do not sufficiently support SARS-CoV-2 emerging directly from these pangolin-CoVs [14]. Obtaining related viral sequences from animal sources would be the most definitive approach to identify the zoonotic origin of a virus [6]. For instance, the full-length genome sequences of viruses isolated from palm civets and camels are 99.8% and 99.9% identical to human SARS-CoV and MERS-CoV [15, 16], respectively, thus consolidating that civets are the intermediate host for SARS-CoV and camels for MERS-CoV. In contrast, RaTG13 shares a genome identity of 96.2% with SARS-CoV-2 [3], and pangolin-CoVs only 85-93% [8-11, 13], which is not high enough to justify that bats or pangolins are a direct zoonotic host of SARS-CoV-2.

Early studies assumed that the outbreak of SARS-CoV-2 was associated with the Huanan Seafood and Wildlife Market, where one or more animals sold there may be the direct zoonotic source [1, 3, 8]. However, this point of view was challenged by the report that the first case of infection was suggested not to be related to the market [17, 18]. Therefore, strategies to trace back the origin of SARS-CoV-2 should not be limited to the animals sold in the market, but should also include a wide range of wild animals outside the market. Theoretically, all kinds of animals that may have close contact with humans should be investigated, but this would be extremely laborious as well as time- and money-consuming.

ACE2 recognition by SARS-CoV-2 is an important determinant of viral infectivity and host range [5, 19]. It has been reported that many animals can be infected by SARS-CoV-2 [20-28]. In this work, we computationally examined the ACE2 usage of SARS-CoV-2 for 285 vertebrates by modeling the binding energy between the SARS-CoV-2 Spike receptor-binding domain (S-RBD) and host ACE2. The binding data correlate well with the reported experimental studies, perfectly distinguishing the effective ACE2 receptors from the less effective ones. Our results reveal that many mammals could serve as intermediate hosts of SARS-CoV-2. This work presents a fast and reliable computational approach to screen potential animal hosts for further experimental analyses.

## Results

### A computational pipeline for ACE2 usage analysis

Since SARS-CoV-2 utilizes ACE2 to invade host cells, ACE2 usage is considered to be an important determinant of infectivity and host range [5, 29]. To examine the ACE2 usage by SARS-CoV-2, we developed a pipeline to model the binding energy between S-RBD and host ACE2 (Fig. 1). We hypothesized that an effective ACE2 receptor should exhibit a low binding energy (or equivalently, a high affinity) while a poor receptor should have a high binding energy. A total of 321 ACE2 orthologs were collected from NCBI, and 285 of them were analyzed in detail after discarding 36 defective sequences (see Methods and Supporting Information Tables S1, S2, and S3). Homologous structure models were built by Modeller [30] using the crystal structure of the hACE2/S-RBD complex (PDB ID: 6M0J) [31] as a template. Each initial complex model was then optimized using FASPR [32] and EvoEF2 [33] to generate structure ensembles for binding energy calculation (see Methods). The ACE2 that achieved a binding energy below a given cutoff was suggested to be an effective receptor for SARS-CoV-2. During structure modeling and binding energy calculation, the N-glycosylation on ACE2 and S-RBD was ignored because current methods are not well adapted for modeling glycosylated amino acids.

**Fig. 1.**
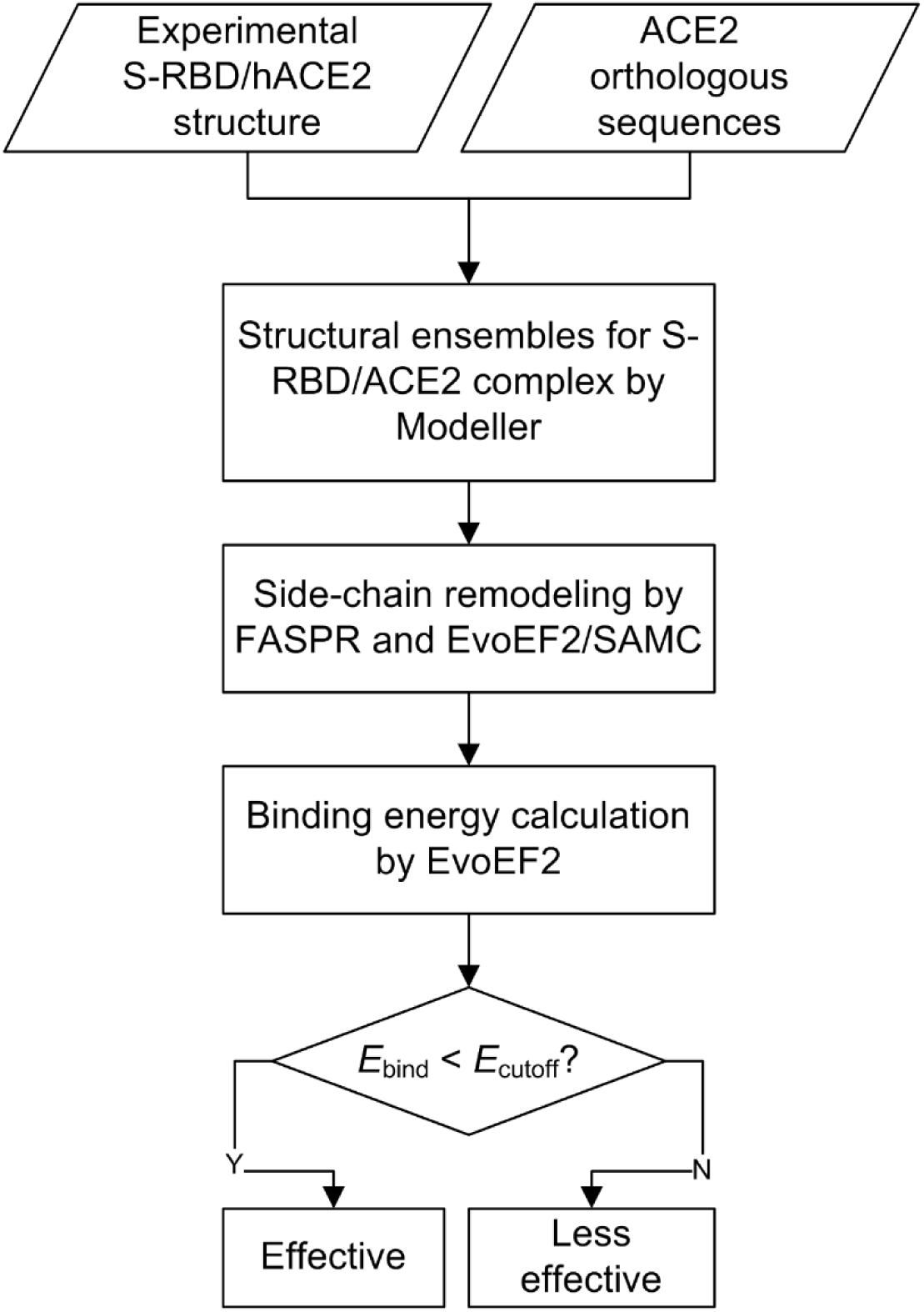
A computational pipeline for ACE2 usage analysis. 321 ACE2 orthologs are downloaded from NCBI. The crystal structure of the hACE2/S-RBD complex (PDB ID: 6M0J) was used as a template for homology modeling. For each ACE2/S-RBD pair, 100 initial Modeller complex models were constructed and repacked by FASPR, and then five models were generated by EvoEF2/SAMC remodeling for each FASPR model. The binding energy cutoff (*E*_cutoff_) was set to be -47 EvoEF2 energy units.

### Binding energy assessment and correlation with experimentally determined ACE2 usage information

The length of 285 ACE2 protein sequences range from 431 to 872 amino acids (Supporting Information Table S3), where most ACE2 sequences are composed of about 800 amino acids (Supporting Information Fig. S1a). Five ACE2 sequences are partial (*Bison bison bison, Thamnophis sirtalis, Haliaeetus albicilla, Fulmarus glacialis*, and *Panthera tigris altaica*), but there are no missing interface residues according to sequence analysis. The ACE2 orthologs share a sequence identity of ≥55% with hACE2 (Supporting Information Fig. S1b), indicating that ACE2 proteins are conserved. Therefore, reliable structure models can be built by homology modeling. Based on the experimental structure of the hACE2/S-RBD complex, 20 residues (Q24, T27, F28, D30, K31, H34, E35, E37, D38, Y41, Q42, L79, M82, Y83, N330, K353, G354, D355, R357, and R393) are present at the interface of hACE2 within 4 Å of S-RBD [31]. Among the ACE2 orthologs, the sequence identities of the 20 interface residues to hACE2 range from 30% to 100% (Supporting Information Fig. S1c), while the sequence identities for the five key interface residues (K31, E35, D38, M82, and K353), which were regarded as important elements to determine host range [29], vary from 0 to 100% (Supporting Information Fig. S1d).

The binding energy for 285 ACE2 proteins ranged from -56.21 to -33.30 EvoEF2 energy units (EEU) (Supporting Information Table S4), where a lower energy represents a stronger binding affinity, which may correspond to a higher susceptibility to SARS-CoV-2. However, one the one hand, it was unknown how trustable the energy values were. On the other hand, given a binding energy, it was not easy to understand whether or not an ACE2 was suggested to be an effective receptor. To address such issues, we compared the calculated binding energy with the experimental ACE2 usage data. Table 1 summarizes the reported infection cases in nature, and the infection studies *in vivo* and *in vitro*. Infection in nature represents that an infected case takes place naturally and has been confirmed by experiments such as quantitative real-time PCR [34]. *In vivo* infection means that the caged experimental animals can be infected by SARS-CoV-2 [20, 24], while *in vitro* infection signifies that ACE2-expressing cells (e.g. HeLa cells transiently expressing ACE2) are permissive to SARS-CoV-2 infection [3]. The discrepancy may exist between *in vivo, in vitro*, and natural infections due to different experimental settings. For instance, it was reported that SARS-CoV-2 replicates poorly in dogs and pigs *in vivo* [20], but it was shown that ACE2 of dogs and pigs could be effectively used for viral entry *in vitro* [3, 35]. Moreover, pet dogs were reported to be infected naturally by their owners with COVID-19 [23]. In this situation, an animal’s ACE2 protein was regarded as an effective receptor to SARS-CoV-2 if any kind of experimental evidence holds.

**Table 1.** The 59 animals whose ACE2 proteins are shown to be effective or less effective for SARS-CoV-2 entry by natural infection and/or experimental studies. The table is organized by ranking the binding energy from low to high.

| Index | Animal name | Binding energy (EEU)<sup>a</sup> | ACE2 usage<sup>b</sup> | Experimental evidence |
| --- | --- | --- | --- | --- |
| 1 | Sumatran orangutan | -56.21 | Y | *In vitro* [35] |
| 2 | Western gorilla | -55.84 | Y | *In vitro* [35] |
| 3 | Olive baboon | -55.77 | Y | *In vitro* [35] |
| 4 | Silvery gibbon | -55.73 | Y | *In vitro* [35] |
| 5 | Crab-eating macaque | -55.38 | Y | *In vitro* [35] |
| 6 | Gelada | -55.29 | Y | *In vitro* [35] |
| 7 | Rhesus macaque | -55.24 | Y | *In vitro* [35], *in vivo* [27, 28] |
| 8 | Human | -55.16 | Y | Natural [2, 58] |
| 9 | Golden snub-nosed monkey | -55.09 | Y | *In vitro* [35] |
| 10 | Chimpanzee | -54.97 | Y | *In vitro* [35] |
| 11 | Ugandan red colobus | -54.79 | Y | *In vitro* [35] |
| 12 | Golden hamster | -53.84 | Y | *In vitro* [35], *in vivo* [25] |
| 13 | Chinese hamster | -53.77 | Y | *In vitro* [35] |
| 14 | Steller sea lion | -53.47 | Y | *In vitro* [35] |
| 15 | Horse | -52.95 | Y | *In vitro* [7, 35] |
| 16 | Amur tiger | -52.93 | Y | Natural [59] |
| 17 | Goat | -52.86 | Y | *In vitro* [7, 35] |
| 18 | Rabbit | -52.84 | Y | *In vitro* [7, 35] |
| 19 | Wild yak | -52.83 | Y | *In vitro* [35] |
| 20 | Puma | -52.79 | Y | *In vitro* [35] |
| 21 | Leopard | -52.74 | Y | *In vitro* [35] |
| 22 | Cattle | -52.71 | Y | *In vitro* [7, 35] |
| 23 | Hawaiian monk seal | -52.56 | Y | *In vitro* [35] |
| 24 | Ferret | -52.55 | Y | *In vitro* [35], *in vivo* [20, 24]; |
| 25 | California sea lion | -52.53 | Y | *In vitro* [35] |
| 26 | Water buffalo | -52.45 | Y | *In vitro* [35] |
| 27 | Lesser egyptian jerboa | -52.44 | Y | *In vitro* [35] |
| 28 | Cat | -52.33 | Y | *In vitro* [7, 35], *in vivo* [20], natural [21] |
| 29 | Canada lynx | -52.21 | Y | *In vitro* [35] |
| 30 | Giant panda | -52.21 | Y | *In vitro* [35] |
| 31 | White-footed mouse | -52.15 | Y | *In vitro* [35] |
| 32 | Sheep | -52.09 | Y | *In vitro* [7, 35] |
| 33 | Beluga whale | -51.98 | Y | *In vitro* [35] |
| 34 | Sperm whale | -51.94 | Y | *In vitro* [35] |
| 35 | Polar bear | -51.93 | Y | *In vitro* [35] |
| 36 | Yangtze finless porpoise | -51.85 | Y | *In vitro* [35] |
| 37 | Malayan pangolin | -51.73 | Y | *In vitro* [7, 35] |
| 38 | Red fox | -51.58 | Y | *In vitro* [35] |
| 39 | Dog | -51.38 | Y | *In vitro* [35], *in vivo* [20], natural [23] |
| 40 | Southern white rhinoceros | -51.08 | Y | *In vitro* [35] |
| 41 | Pig | -50.74 | Y | *In vitro* [3, 7] |
| 42 | Arctic ground squirrel | -50.62 | Y | *In vitro* [35] |
| 43 | Chinese rufous horseshoe bat | -49.91 | Y | *In vitro* [35] |
| 44 | Bactrian camel | -49.88 | Y | *In vitro* [7, 35] |
| 45 | Killer whale | -49.47 | Y | *In vitro* [35] |
| 46 | Long-finned pilot whale | -49.19 | Y | *In vitro* [35] |
| 47 | Atlantic bottle-nosed dolphin | -49.05 | Y | *In vitro* [35] |
| 48 | Yangtze river dolphin | -49.05 | Y | *In vitro* [35] |
| 49 | Masked palm civet | -47.97 | Y | *In vitro* [3, 7] |
| 50 | Malayan tiger | n.d. | Y | Natural [59] |
| 51 | African lion | n.d. | Y | Natural [59] |
| 52 | Mink | n.d. | Y | Natural [26] |
| 53 | Marmoset | -46.81 | N | *In vitro* [35], *in vivo* [28] |
| 54 | Black-capped squirrel monkey | -46.71 | N | *In vitro* [35] |
| 55 | Tufted capuchin | -46.13 | N | *In vitro* [35] |
| 56 | Brown rat | -43.14 | N | *In vitro* [7, 35] |
| 57 | House mouse | -42.62 | N | *In vitro* [7, 35] |
| 58 | Duck | -42.54 | N | *In vivo* [20] |
| 59 | Chicken | -42.07 | N | *In vitro* [7], *in vivo* [20] |

<sup>a</sup>: The binding energy was not calculated for Malayan tiger, lion, and mink because their ACE2 proteins were not included in the list of 321 ACE2 orthologs. EEU stands for EvoEF2 energy unit.
<sup>b</sup>: Y, effective ACE2 receptors; N, less effective ACE2 receptors. An ACE2 protein is classified as effective if at least one of the three kinds of experimental evidence holds.

The calculated binding energy correlated well with the experimentally determined ACE2 usage data; the ACE2 proteins that can be more effectively used by SARS-CoV-2 achieved a relatively lower binding energy (Table 1). A binding energy cutoff of -47 EEU was able to discriminate the more efficient ACE2 receptors from the less efficient ones (Table 1), with the maximum Matthews correlation coefficient (MCC) of one (Supporting Information Fig. S2). Among all the experimental species, apes (Sumatran orangutan, western gorilla, silvery gibbon, and chimpanzee) and Old-World monkeys (olive baboon, crab-eating macaque, gelada, rhesus macaque, golden snub-nosed monkey, and Ugandan red colobus) and humans achieved the lowest binding energy ranging from -56.21 to -54.79 EEU (Table 1). Besides, a few rodents (golden hamster, Chinese hamster, jerboa, white-footed mouse, and Arctic ground squirrel) and carnivores (sea lion, tiger, puma, leopard, seal, ferret, dog, cat, lynx, and bear) also achieved a relatively low binding energy varying from -53.84 to -50.62 EEU (Table 1). Three New-World monkeys (marmoset, black-capped squirrel monkey, and tufted capuchin), rats, mice, ducks, and chickens achieved a higher binding energy score (>-47 EEU), consistent with the reports that these animals were less susceptible to SARS-CoV-2 [3, 20, 35, 36].

### Binding energy-based intermediate host range prediction

Based on the calculated binding energy and experimental data, we mapped the ACE2 usage effectiveness for all the 285 species (Fig. 2). Fish (including *Actinopterygii, Chondrichthyes*, and *Sarcopterygii*), amphibians, reptiles, and birds were predicted to have a relatively high binding energy (>-47 EEU), suggesting ACE2 of these species may be less permissive to SARS-CoV-2 binding. Mammals showed the broadest binding energy distribution, from -56.21 to -38.67 EEU (Fig. 2 and Supporting Information Table S4). 97 non-human mammals achieved a binding energy below -47 EEU; that is, besides the experimentally-validated species, another 49 species were also predicted to have an effective ACE2 receptor for SARS-CoV-2 (Fig. 2 and Supporting Information Table S4). These results suggest that mammals rather than other species are likely to be the main source of SARS-CoV-2 and hence they should be the major focus. This finding is also consistent with previous studies [7, 20, 35, 37-41], but a more quantitative measurement was given here. Our findings also refute isolated reports claiming that non-mammal vertebrates such as reptiles could be the intermediate host [42, 43].

**Fig. 2.**
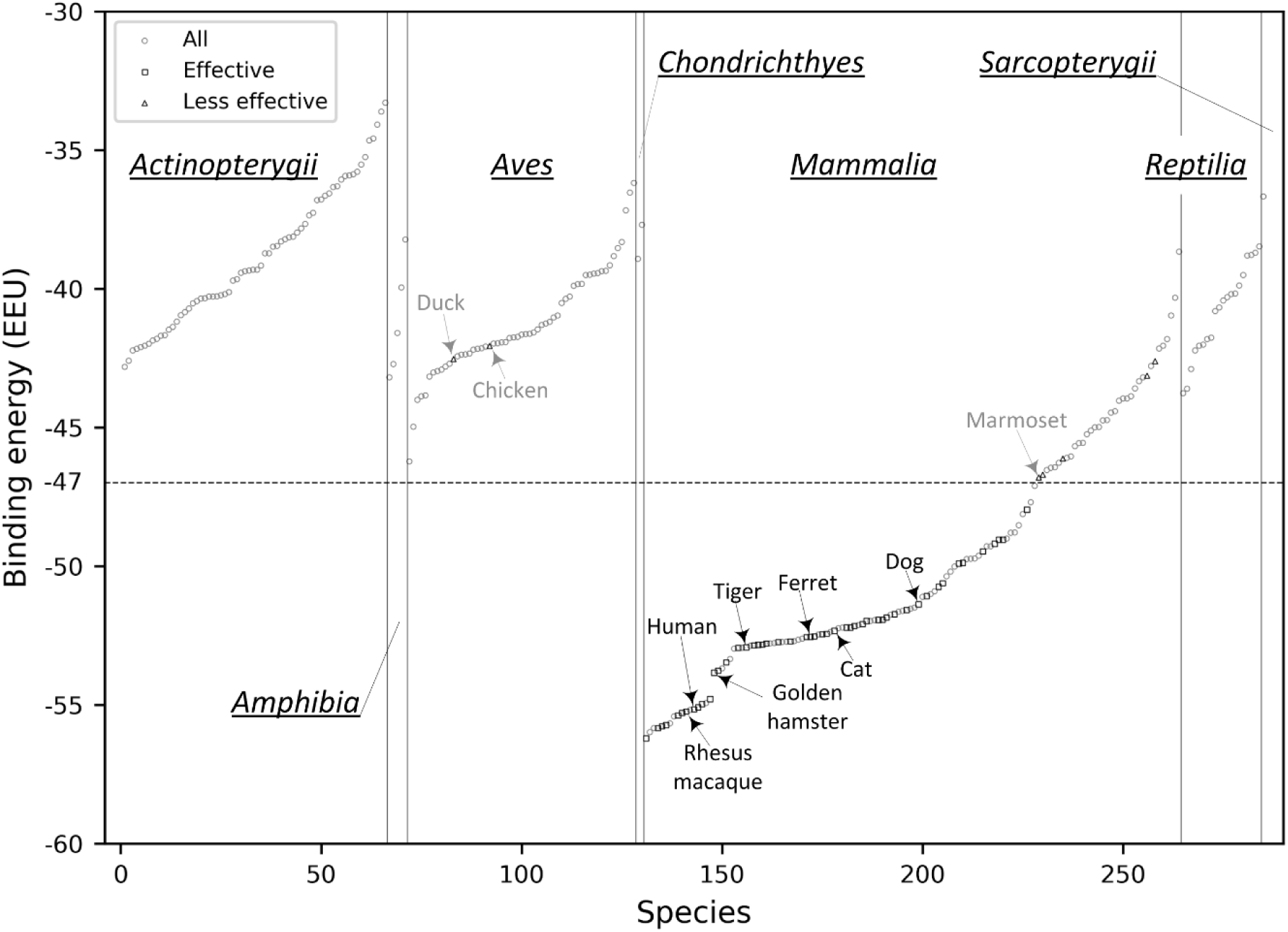
Mapping the calculated binding energy to 285 vertebrates. The ACE2 proteins are categorized by their animal Class (*Actinopterygii, Amphibia, Aves, Chondrichthyes, Mammalia, Reptilia*, and *Sarcopterygii*) and ranked by the binding energy from low to high in each Class. The ACE2 proteins that are experimentally shown to be effective or less effective to SARS-CoV-2 are shown in squares and triangles, respectively, while the others are shown in circles. Susceptible and insusceptible animals are highlighted in black and gray, respectively.

The binding energy calculation did not consider the impact of possible N-glycosylation of ACE2 and Spike. Although no N-glycosylation site is present at the interface of the hACE2/S-RBD complex [31], some ACE2 variants may have N-glycosylation sites at the interface region, which may prevent their binding to S-RBD due to steric hindrance. Thus, the analysis of interface N-glycosylation may help refine the list of effective ACE2 receptors classified by binding energy. N-glycosylation of asparagines occurs predominantly at the NX(T/S) motif, where X is any amino acid except proline. However, not all N-X-(T/S) sequons are glycosylated, so the motif alone may not be sufficient to discriminate between glycosylated and non-glycosylated asparagines. We tried three predictors, NGlycPred [44], N-GlyDE [45], and NetNGlyc (http://www.cbs.dtu.dk/services/NetNGlyc/), to predict N-glycosylation on hACE2. None of them could accurately predict all the experimentally identified N-glycosylation sites (Supporting Information Table S5). All seven NX(T/S) motifs are glycosylated in the experimentally determined structure (PDB ID: 6M17) [46], indicating that ACE2 is highly N-glycosylated. To avoid the omission of potential glycosylation sites, we systematically examined all of the NX(S/T) motifs for the 285 ACE2 proteins and manually checked if any N-glycosylation sites were present at the interface.

64 out of the 285 ACE2 proteins were found to have one or more interface glycosylation sites, including 22 fish, one amphibian, 27 birds, seven mammals, and seven reptiles (Supporting Information Table S6). Since many mammals are likely susceptible to SARS-CoV-2 (Fig. 2 and Supporting Information Table S4), we examined the seven mammals and mapped the putative interface N-glycosylation sites into their structure models. Interestingly, none of the effective ACE2 receptors in Table 1 has an interface N-glycosylation site. The seven mammals were the Eurasian common shrew (*Sorex araneus*), small Madagascar hedgehog (*Echinops telfairi*), western European hedgehog (*Erinaceus europaeus*), aardvark (*Orycteropus afer*), big brown bat (*Eptesicus fuscus*), star-nosed mole (*Condylura cristata*), and greater horseshoe bat (*Rhinolophus ferrumequinum*), where their binding energies were -40.97, -44.99, -38.67, -48.79, -46.46, -45.56, and -44.47 EEU, respectively. Following the binding energy criterion, aardvark’s ACE2 was predicted to an effective receptor, but it may be ineffective due to glycosylation. The shrew had two interface glycosylation sites, N23 and N41, which form hydrogen bonds with N487 and Y449, respectively (Fig. 3a); the aardvark had only one interface glycosylation site at N38, forming two hydrogen bonds with Y449 and Q498 (Fig. 3b). Since these asparagine residues could form direct contact with S-RBD, their glycosylation may hinder the binding of the two proteins (i.e. ACE2 and Spike).

**Fig. 3.**
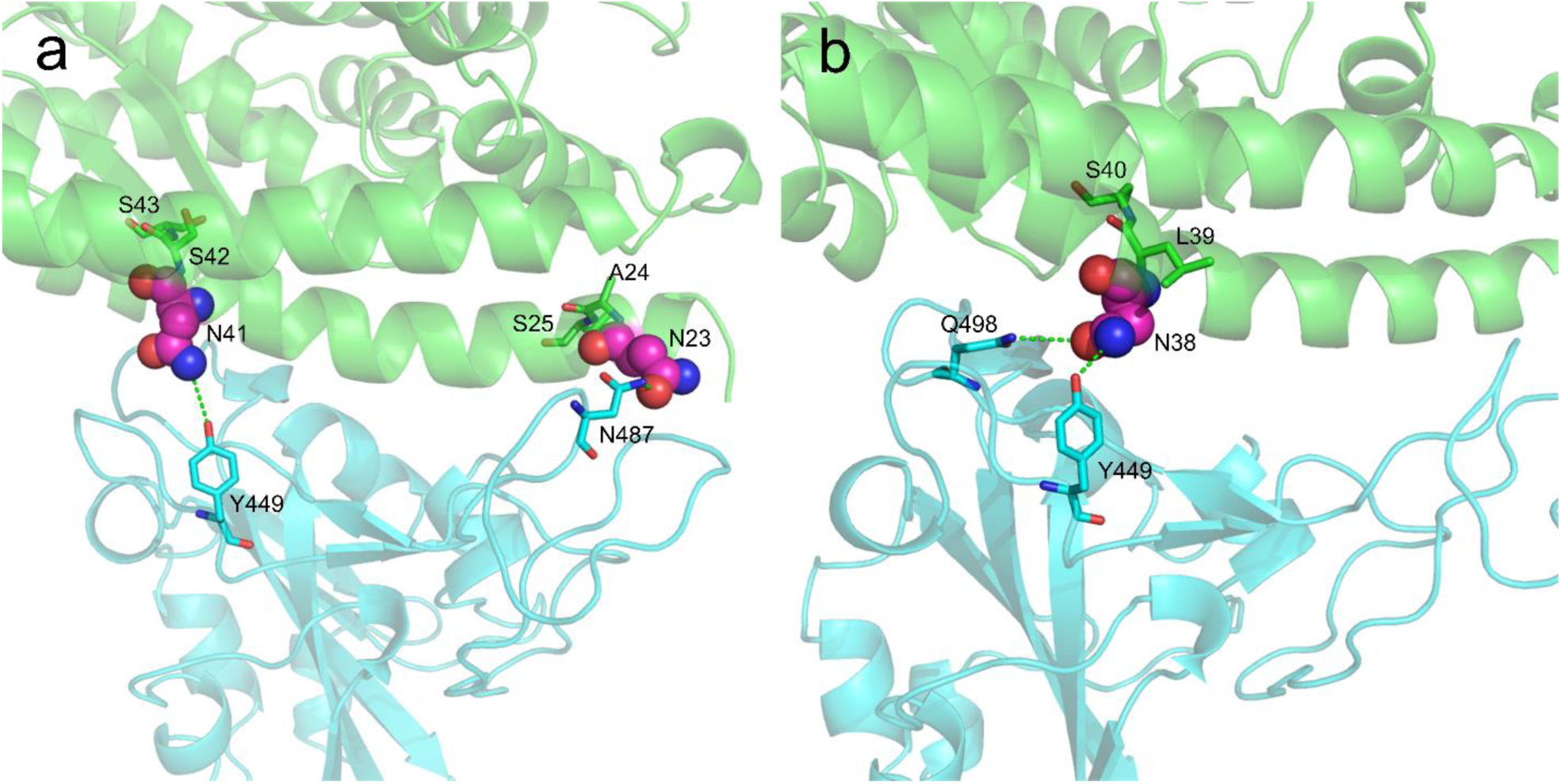
Putative N-glycosylation sites at the interface of two example ACE2/S-RBD complex structures. (a) Eurasian common shrew (*Sorex araneus*); and (b) Aardvark (*Orycteropus afer*). ACE2 and S-RBD are shown in green and cyan cartoons, respectively. The potential interface N-glycosylation motifs are shown with the asparagine residues highlighted in spheres.

Following the binding energy calculation and interface N-glycosylation site analysis, 96 non-human ACE2 proteins were suggested to be effectively utilized by SARS-CoV-2; half of them have been confirmed by experiments (Table 1) and the other half is summarized in Table 2. Therefore, compared with the original list of 285 animals, our method considerably narrowed the host range. The predicted potential zoonotic animals are distributed widely, including pets, domestic, agricultural, and zoological animals that may have close contact with humans (Tables 1 and 2).

**Table 2.** 48 other animals were predicted to have an effective ACE2 receptor capable of S-RBD binding. The table is organized by ranking the binding energy from low to high. The animals in Table 1 were not included in this table. The binding energy cutoff (−47 EEU) was chosen by maximally discriminating the experimentally determined effective ACE2 receptors from the less effective ones.

| Index | Species | Animal name | Binding energy (EEU) |
| --- | --- | --- | --- |
| 1 | *Pan paniscus* | Bonobo | -55.99 |
| 2 | *Nomascus leucogenys* | Northern white-cheeked gibbon | -55.84 |
| 3 | *Chlorocebus sabaues* | Green monkey | -55.67 |
| 4 | *Macaca nemestrina* | Pig-tailed macaque | -55.42 |
| 5 | *Cercocebus atys* | Sooty mangabey | -55.19 |
| 6 | *Mandrillus leucophaeus* | Drill | -54.94 |
| 7 | *Nannospalax galili* | Northern israeli blind subterranean mole rat | -53.69 |
| 8 | *Propithecus coquereli* | Coquerel's sifaka | -53.35 |
| 9 | *Callorhinus ursinus* | Northern fur seal | -52.98 |
| 10 | *Equus przewalskii* | Mongolian wild horse | -52.95 |
| 11 | *Acinonyx jubatus* | Cheetah | -52.88 |
| 12 | *Heterocephalus glaber* | Naked mole-rat | -52.79 |
| 13 | *Bison bison bison* | Plains bison | -52.78 |
| 14 | *Mustela erminea* | Ermine | -52.74 |
| 15 | *Phoca vitulina* | Harbor seal | -52.73 |
| 16 | *Bos indicus x Bos taurus* | Hybrid cattle | -52.71 |
| 17 | *Lontra canadensis* | Northern american river otter | -52.66 |
| 18 | *Odobenus rosmarus divergens* | Walrus | -52.62 |
| 19 | *Bos indicus* | Zebu cattle | -52.47 |
| 20 | *Peromyscus maniculatus bairdii* | Deer mouse | -52.38 |
| 21 | *Odocoileus virginianus texanus* | White-tailed deer | -52.24 |
| 22 | *Fukomys damarensis* | Damaraland mole-rat | -52.22 |
| 23 | *Ursus arctos horribilis* | Grizzly bear | -52.13 |
| 24 | *Monodon monoceros* | Narwhal | -51.98 |
| 25 | *Phocoena sinus* | Vaquita | -51.95 |
| 26 | *Microtus ochrogaster* | Prairie vole | -51.76 |
| 27 | *Balaenoptera acutorostrata scammoni* | North pacific minke whale | -51.65 |
| 28 | *Marmota marmota* | Alpine marmot | -51.62 |
| 29 | *Ictidomys tridecemlineatus* | Thirteen-lined ground squirrel | -51.54 |
| 30 | *Marmota flaviventris* | Yellow-bellied marmot | -51.50 |
| 31 | *Canis lupus dingo* | Dingo | -51.11 |
| 32 | *Ochotona princeps* | American pika | -51.01 |
| 33 | *Rousettus aegyptiacus* | Egyptian rousette | -50.91 |
| 34 | *Lagenorhynchus obliquidens* | Pacific white-sided dolphin | -50.37 |
| 35 | *Nyctereutes procyonoides* | Raccoon dog | -50.20 |
| 36 | *Equus asinus* | Donkey | -50.02 |
| 37 | *Mirounga leonina* | Southern elephant seal | -49.75 |
| 38 | *Camelus dromedarius* | Arabian camel | -49.74 |
| 39 | *Phyllostomus discolor* | Pale spear-nosed bat | -49.73 |
| 40 | *Camelus ferus* | Wild bactrian camel | -49.63 |
| 41 | *Pteropus vampyrus* | Large flying fox | -49.29 |
| 42 | *Pteropus alecto* | Black flying fox | -49.29 |
| 43 | *Dipodomys ordii* | Ords kangaroo rat | -49.01 |
| 44 | *Loxodonta africana* | African savanna elephant | -48.80 |
| 45 | *Enhydra lutris kenyon* | Sea otter | -48.53 |
| 46 | *Trichechus manatus latirostris* | Florida manatee | -48.13 |
| 47 | *Octodon degus* | Common degu | -47.70 |
| 48 | *Vicugna pacos* | Alpaca | -47.11 |

### Case studies

We then analyzed several ACE2 proteins to show molecular details about why they may or may not be effectively used by SARS-CoV-2 as an entry receptor. The first case is a New-World monkey, marmoset (*Callithrix jacchus*), which is of extremely low susceptibility to SARS-CoV-2 both *in vivo* and *in vitro* [28, 35]. The marmoset achieved a high binding score of -46.81 EEU. Compared with hACE2, there were four residue substitutions in the marmoset ACE2, i.e. Y41H, Q42E, M82T, and G354Q (Table 3). In hACE2, Y41 could form hydrogen bonds with T500 in the RBD; Q42 could form a hydrogen bond with the carbonyl group of G446 and another hydrogen bond with Y449 where the NE2 atom of Q42 acts as the donor and the OH atom of Y446 as the acceptor (Fig. 4a, left). The substitution of Y41 into histidine not only results in a reduced van der Waals packing energy but also disrupts the favorable hydrogen bond with T500; mutation of Q42 into glutamic acid destroys the two hydrogen bonds with G446 and Y449; moreover, the M82T substitution could lead to a reduced packing interaction with F486 due to the smaller side-chain (Fig. 4a, right). The loss of three hydrogen bonds and the weakened van der Waals forces result in the poor binding energy. As reported, the double mutant H41Y/E42Q made the variant marmoset receptor more permissive to SARS-CoV-2 infection [35]. Besides the New-World monkeys, we found that the ACE2 proteins of four bats (i.e. *Eptesicus fuscus, Myotis brandtii, Myotis davidii*, and *Myotis lucifugus*) also have the Y41H/Q42E substitution (Supporting Information Table S7); interestingly, they were also predicted to be less effective with a binding score of >-47 EEU.

**Table 3.** Comparison of ACE2 interface residues and binding energy for humans, marmosets, pangolins, and turtles. Five key residues are underlined. Amino acid mutations relative to hACE2 are shown in bold.

| Species | Human<br>(*H. sapiens*) | Marmoset<br>(*C. jacchus*) | Pangolin<br>(*M. javanica*) | Turtle<br>(*C. picta*) | Turtle<br>(*C. mydas*) | Turtle<br>(*P. sinensis*) |
| --- | --- | --- | --- | --- | --- | --- |
| ACE2<br>interface<br>residues | Q24 | Q24 | **E24** | **E24** | **E24** | **E24** |
|  | T27 | T27 | T27 | **N27** | **N27** | **N27** |
|  | F28 | F28 | F28 | F28 | F28 | F28 |
|  | D30 | D30 | **E30** | **S30** | **S30** | **S30** |
|  | <u>K31</u> | <u>K31</u> | <u>K31</u> | <u>**Q31**</u> | <u>**Q31**</u> | <u>**E31**</u> |
|  | H34 | H34 | **S34** | **V34** | **V34** | **V34** |
|  | <u>E35</u> | <u>E35</u> | <u>E35</u> | <u>**R35**</u> | <u>**R35**</u> | <u>**Q35**</u> |
|  | E37 | E37 | E37 | E37 | E37 | E37 |
|  | <u>D38</u> | <u>D38</u> | <u>**E38**</u> | <u>D38</u> | <u>D38</u> | <u>D38</u> |
|  | Y41 | **H41** | Y41 | Y41 | Y41 | Y41 |
|  | Q42 | **E42** | Q42 | **A42** | **A42** | **A42** |
|  | L79 | L79 | **I79** | **N79** | **N79** | **N79** |
|  | <u>M82</u> | <u>**T82**</u> | <u>**N82**</u> | <u>**K82**</u> | <u>**K82**</u> | <u>**K82**</u> |
|  | Y83 | Y83 | Y83 | Y83 | Y83 | Y83 |
|  | N330 | N330 | N330 | N330 | N330 | N330 |
|  | <u>K353</u> | <u>K353</u> | <u>K353</u> | <u>K353</u> | <u>K353</u> | <u>K353</u> |
|  | G354 | **Q354** | **H354** | **K354** | **K354** | **K354** |
|  | D355 | D355 | D355 | D355 | D355 | D355 |
|  | R357 | R357 | R357 | R357 | R357 | R357 |
|  | R359 | R359 | R359 | R359 | R359 | R359 |
| Binding energy<br>(EEU) | -55.16 | -46.81 | -51.73 | -43.61 | -42.23 | -40.68 |

**Fig. 4.**
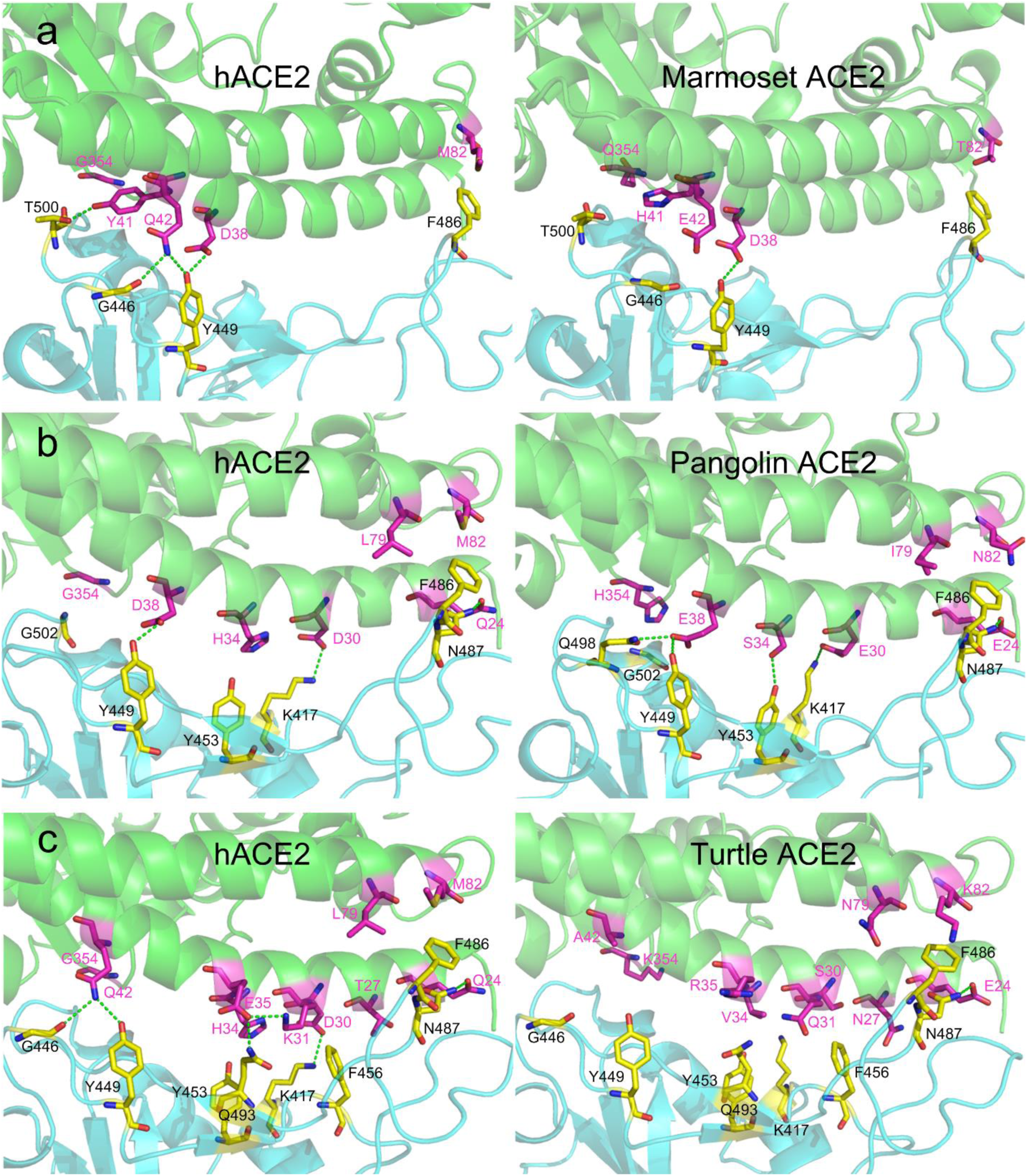
Comparison of the mutated interface between hACE2/S-RBD and animal-ACE2/S-RBD. (a) hACE2/S-RBD versus marmoset-ACE2/S-RBD; (b) hACE2/S-RBD versus pangolin-ACE2/S-RBD; and (c) hACE2/S-RBD versus turtle-ACE2/S-RBD. Residues in ACE2 and S-RBD are shown in magenta and yellow, respectively. Hydrogen bonds are shown in green dashed-lines.

The second case is Malayan pangolin (*Manis javanica*), which has been suggested as a potential intermediate host in a few studies [9, 10]. Pangolin ACE2 shared an identity of 84.8%, 65%, and 60% with hACE2 for all, interface, and the key residues, respectively (Supporting Information Table S4). Although pangolin ACE2 had seven residues mutated compared with hACE2, i.e. Q24E, D30E, H34S, D38E, L79I, M82N, and G354H, it still achieved a relatively low binding energy of -51.73 EEU (Table 3). In the hACE2/S-RBD complex, Q24 forms a hydrogen bond with N487, and D38 forms a hydrogen bond with Y449; D30 forms a salt bridge with K417; L79 and M82 form favorable van der Waals contacts with F486 (Fig. 4b, left). In the pangolin-ACE2/S-RBD complex, favorable interactions are also extensively formed. E38 could form two hydrogen bonds with Q498 and Y449; E30 and E24 could form a hydrogen bond with K417 and N487, respectively; S34 could form a hydrogen bond with Y453 though it has a reduced van der Waals interaction due to the small size compared with H34; I79 and N82 could also form favorable packing interactions with F486 (Fig. 4b, right). Therefore, although pangolin ACE2 achieved a higher binding score than hACE2, probably due to worse contacting geometries, the extensive favorable interactions demonstrate that pangolin ACE2 can still be an effective receptor to SARS-CoV-2. Thus, the binding analysis and molecular details supported Malayan pangolin as a possible intermediate host.

The third case is turtles (*Chrysemys picta, Chelonia mydas*, and *Pelodiscus sinensis*), which have been suggested as a potential intermediate host by Liu et al [43]. They argued that turtles have two important residues (Y41 and K353) in their ACE2 that are identical to those in hACE2 and that turtles in the markets were more common than pangolins [43]. Although it may be true that Y41 and K353 play an important role in binding S-RBD, it is, however, not a unique feature in the ACE2 of turtles and humans. As shown, many mammals have Y41 and K353 in their ACE2 proteins (Supporting Information Table S7). Besides, the first reported case of infection was suggested not to be associated with the market [17]. Therefore, their rules for screening intermediate hosts were not persuasive. The ACE2 protein of these turtles has ten amino acid substitutions compared with hACE2 (Table 3). *C. picta* and *C. mydas* have identical interface residues in their ACE2 proteins. *P. sinensis* has two different interface residues (E31 and Q35) compared with *C. picta* and *C. mydas* (Table 3). In *C. picta*, only E24 could form a hydrogen bond with N487 (Fig. 4c, right), while the other mutations resulted in a substantial loss of favorable hydrogen bonds and salt bridges compared with those in the hACE2/S-RBD (Fig. 4c, left). Expectedly, the three turtles (i.e. *C. picta, C. mydas*, and *P. sinensis*) achieved a poor binding score of -43.61, -42.23, and -40.68 EEU, respectively. Therefore, structure modeling did not support turtles as intermediate hosts.

## Discussion

As the COVID-19 pandemic continues, the direct zoonotic origin (intermediate host) of SARS-CoV-2 remains elusive. Many animals have been reported to be infected by SARS-CoV-2 in nature or the laboratory, suggesting a possibly wide host range for this novel coronavirus. Currently, the number of animals that have been experimentally tested is very small compared to the huge number of animal species. Previous studies suggested that receptor recognition is an important determinant of host range [5, 19, 29]. Therefore, we proposed a computational pipeline for identifying the intermediate hosts of SARS-CoV-2 by modeling the binding affinity between host ACE2 and the viral S-RBD.

### The reasonability of ignoring TMPRSS2

It has been shown that SARS-CoV-2 cell entry depends on ACE2 and the serine protease TMPRSS2 [47]. However, we did not consider the role of TMPRSS2 for host prediction, due to the following reasons. First, TMPRSS2’s role for priming spike may be replaced by some other proteases like cathepsin B and L [48]. Second, different from ACE2 which is used as a binding receptor only, TMPRSS2 cleaves Spike through chemical catalysis. Thus, to quantify the impact of TMPRSS2, its catalytic activity for cleavage needs to be predicted; this is, however, an impossible task to achieve at present, as almost all protease cleavage predictors were trained to predict cleavage sites for one known protease of one species [49]. Third, TMPRSS2 proteases from different species may be similarly efficient. This is supported in part by the fact that wild-type mice are insusceptible to SARS-CoV-2, while transgenic mice that express hACE2 can be infected [50], suggesting that mouse TMPRSS2 may be sufficiently efficient at cleaving Spike. Besides, a recent study showed that computational modeling failed to distinguish the binding capability of TMPRSS2 from different animals [41]. As a result, we believe that it may be reasonable to ignore TMPRSS2 for host prediction.

### ACE2 Sequence analysis alone is not accurate enough for host identification

Built on the fact that hACE2 is highly susceptible to SARS-CoV-2, many previous studies only performed sequence analyses and used the sequence identity between animal ACE2 proteins and hACE2 to predict intermediate hosts [38, 43, 51], as it is believed that the ACE2 proteins that are similar to hACE2 may also be susceptible [6, 29]. We calculated the MCCs for distinguishing experimentally determined effective ACE2 receptors from the less effective ones listed in Table 1 using sequence identity. The maximum MCCs were 0.51,0.73, and 0.53 with the optimum sequence identity cutoff of 66%-78%, 61%∼65%, and ≤60% in terms of all, interface, and key residues, respectively (Supporting Information Fig. S3), which were much lower than that achieved by the classification via binding energy assessment (Supporting Information Fig. S2). Four New-World monkeys (*Sapajus apella, Aotus nancymaae, Saimiri boliviensis*, and *Callithrix jacchus*) share a relatively high sequence identity of >92%, 80%, and 80% with hACE2 in terms of all, interface, and key residues, respectively (Supporting Information Tables S4 and S7). Following the optimum sequence identity cutoffs, the ACE2 proteins of these New-World monkeys were predicted to be very effective receptors. However, *in vivo* infection studies showed *C. jacchus* was not susceptible to SARS-CoV-2 [28]; *in vitro* experiments also suggested that the ACE2 proteins of *S. apella, S. boliviensis*, and *C. jacchus* cannot be used by SARS-CoV-2 [35].

In contrast, dogs, cats, and ferrets, which have a much lower sequence identity to hACE2 than the new-world monkeys (Supporting Information Table S4), can be infected by SARS-CoV-2 in nature and/or *in vivo* [20-24]. These results suggest that an ACE2 protein with a higher sequence identity to hACE2 is not necessarily an effective receptor, whereas those with a lower identity is not necessarily a poor one. Therefore, sequence identity between hACE2 and animal ACE2 may not be a good descriptor for host identification.

### Binding energy is a better descriptor for host prediction

As indicated by the high MCC achieved (Supporting Information Fig. S2), structure-based binding energy assessment was more accurate than sequence identity for distinguishing experimentally confirmed species, provided that high-quality structure models were used. Critically, the structure models are very likely to be very reliable given the high sequence similarity between hACE2 and the ACE2 orthologs and the application of advanced structure modeling tools [30, 32].

Moreover, we argued that it is critical to model binding energy using structure ensembles rather than a single model. We found that the binding scores that were calculated for different models of the same ACE2/S-RBD complex fluctuated considerably (Supporting Information Fig. S4). The maximum MCC of the classification by the binding energy derived from the first model was only about 0.63 (Supporting Information Fig. S5), suggesting a single model was not sufficiently accurate for the classification even if a perfect scoring function is available. To circumvent the randomness of binding energy from a single model, we evaluated a large ensemble of structure models (e.g. 500 models in this work) for each complex and took the lowest binding score as the binding energy for ACE2 usage analysis. With a proper threshold (i.e. -47 EEU), the binding energy calculated in this way correlated well with experimental data, perfectly distinguishing the experimentally determined effective ACE2 receptors from the less effective ones with the maximum MCC of one (Supporting Information Fig. S2).

### Identification and screening of potential zoonotic origins

The most definitive strategy to identify the direct zoonotic origins of SARS-CoV-2 is to isolate related viruses from animal sources [6]. Unlike SARS-CoV and MERS-CoV, whose direct zoonotic origins were identified to be civets [15] and camels [16], respectively, soon after their outbreak, the clue for SARS-CoV-2 remain elusive as the first reported case of infection was suggested not to be associated with the Huanan Seafood and Wildlife Market [17, 18]. As a result, a large number of animals have to be sampled to isolate viral strains that are highly similar to SARS-CoV-2 (e.g. >99% genome identity); this is a formidable task that would require extensive effort. In this regard, our work presents a fast, yet reliable approach for screening potential animals for further analysis.

Our result suggests that many mammals are likely to be potential intermediate hosts of SARS-CoV-2, which is consistent with a few recent studies [35, 37, 52]. Here, the ACE2 proteins of 285 species were assessed because their sequences were of good quality. In reality, there are more animals whose ACE2 proteins have not been sequenced yet. Thus, although 96 mammals in this study were predicted to have an effective ACE2 receptor capable of binding SARS-CoV-2 Spike, it does not necessarily mean that the real intermediate host must be one of them. The list may be further screened by considering the living environment of animals. For instance, some mammals like whales and dolphins live in the water, and therefore the chance for them to transmit bat viruses to humans may be extremely low, considering that bats are terrestrial animals.

## Conclusion

The direct zoonotic origin (intermediate host) of SARS-CoV-2 that caused the COVID-19 pandemic remains elusive. In this work, we developed a computational pipeline to facilitate the identification of potential intermediate hosts of SARS-CoV-2 by modeling the binding affinity between the SARS-CoV-2 Spike receptor-binding domain and the ACE2 protein of host animals. The effectiveness of this method was verified by its performance on perfectly distinguishing the experimentally determined effective ACE2 receptors from the less effective ones with a maximum Matthews correlation coefficient (MCC) of one. Although the sequence identity-based descriptors have been widely used for predicting intermediate hosts, our results showed that their performance for discriminating between effective and less effective receptors was much worse than the binding-affinity based approach proposed here by achieving a maximum MCC of 0.73. Our results reveal that SARS-CoV-2 may have a broad host range and a few mammals, especially some primates, rodents, and carnivores, rather than the non-mammal animals could be potential hosts of SARS-CoV-2. Besides, as a supplementarity of our previous pangolin coronavirus genome assembly studies, the detailed structural modeling here also supported pangolins as a possible intermediate host with molecular-level insights. Since these animals are likely to be susceptible to SARS-CoV-2, continuous monitoring of viral circulation in these animals is very important for disease control and wildlife protection efforts.

## Materials and methods

### Collection and examination of ACE2 orthologs

A list of ACE2 orthologs from 318 vertebrate species was downloaded from the NCBI website (https://www.ncbi.nlm.nih.gov/gene/59272/ortholog/?scope=7742). Besides these, we also considered the ACE2 orthologs from three mammals that are not included in this list, namely, palm civets, raccoon dogs, and Chinese rufous horseshoe bats, as civets and raccoon dogs were suggested to be intermediate hosts of SARS-CoV [15]. Additionally, it was shown that the ACE2 proteins of civets and horseshoe bats can also be utilized by SARS-CoV-2 for viral entry in cell-level experiments [3].

Among the 321 ACE2 orthologs, 30 sequences had one or more amino acids that were either nonstandard or incorrectly parsed, i.e. annotated as ‘X’, and thus these ACE2 orthologs were excluded from the detailed analysis (Supporting Information Table S1). Moreover, sequence alignment analysis (see below) showed that the ACE2 proteins of six species had five or more missing S-RBD binding residues (Supporting Information Table S2), i.e. *Acanthisitta chloris* (protein accession ID: XP_009082150.1), *Apteryx mantelli mantelli* (XP_013805736.1), *Salmo salar* (XP_014062928.1), *Rhinopithecus bieti* (XP_017744069.1), *Leptonychotes weddellii* (XP_030886750.1), and *Petromyzon marinus* (XP_032835032.1). Subsequent binding analysis (see below) showed that these ACE2 receptors had a much higher binding energy (and thus a lower binding capability) than the others (Supporting Information Table S2), partly because of the incomplete binding interface. Therefore, we cannot suggest whether these animals are susceptible to SARS-CoV-2 based on the defective information.

The remaining 285 ACE2 orthologs are summarized in Supporting Information Table S3, including 134 mammals (*Mammalia*), 57 birds (*Aves*), 69 fish (*Actinopterygii* (66), *Chondrichthyes* (2), *and Sarcopterygii* (1)), 20 reptiles (*Reptilia*), and five amphibians (*Amphibia*). The protein ID, scientific classification (*Class* and *Species*), and common name are provided for easy retrieval.

### Sequence analyses

291 ACE2 sequences, including the six ACE2 proteins with missing S-RBD binding residues in Supporting Information Table S2, were subjected to multiple sequence alignment (MSA) analysis using Clustal Omega [53] with default parameters. Pairwise sequence identities between the full-length sequence of hACE2 (accession ID: NP_001358344.1) and the other ACE2 sequences were calculated based on the MSAs. Besides the full-length sequence identities, the sequence identities for the 20 interface residues [31] and five critical S-RBD binding residues [29] were also calculated from the MSAs. The results for these three types of sequence identities are shown in Supporting Information Table S4.

### Structure modeling

It should be mentioned that in reality, the ACE2 receptors of some animals may not bind to S-RBD. However, to quantitatively compare the capability of different ACE2 receptors to bind to S-RBD, we first constructed initial ACE2/S-RBD complex models through homology modeling, assuming that all the ACE2 receptors could bind to S-RBD, and then computed the binding energies between the two partners.

Pairwise sequence alignments between hACE2 and the other ACE2 orthologs were extracted from the MSAs and trimmed accordingly, as hACE2 was not full-length in the template complex (PDB ID: 6m0j) [31]. The trimming should not affect binding analysis because it was shown that the protein-protein interface is unabridged in the experimentally determined hACE2/S-RBD complex structures [5, 31, 54]. We utilized Modeller v9.24 [30] to build the initial putative complex models. Each model was first optimized with the variable target function method (VTFM) with conjugate gradients (CG) using parameters *library_schedule = autosched*.*slow* and *max_var_iterations = 300*, and then refined using molecular dynamics (MD) with simulated annealing (SA) using parameter settings *md_level = refine*.*slow*. The whole cycle was repeated two times and was not stopped unless the objective function was >1e6 (parameter settings: *repeat_optimization = 2* and *max_molpdf = 1e6*). For each ACE2/S-RBD pair, 100 initial Modeller complex models were constructed.

### Binding energy calculation

Before binding energy calculation, each Modeller complex model was first repacked using FASPR [32] to eliminate rotamer outliers and then the interface residue side-chain conformations were thoroughly refined (both for ACE2 and S-RBD) using the EvoEF2 force field in conjunction with a simulated annealing Monte Carlo (SAMC) optimization procedure [33, 55], which was also utilized for anti-SARS-CoV-2 peptide design [56]. During the side-chain refinement process, both the ACE2 and S-RBD sequences were kept fixed, while the different rotameric side-chain conformations were sampled. Since a stochastic SAMC optimization procedure was used, obtaining the global energy minimum may not always be guaranteed. Therefore, the optimization of the interface residues was performed five times independently to generate five refined low-energy models. Hence, for each ACE2/S-RBD pair, 500 final models were generated and scored using EvoEF2 [33]. The minimum binding interaction score achieved among all 500 complex models was regarded as the binding energy.

## Author contributions

Y.Z. conceived and supervised the project; X.H. refined the structure models, performed sequence, structure, binding, and N-glycosylation analysis, and drafted the manuscript; C.Z. constructed the initial structure models; R.P. and G.S.O. participated in the discussion and edited the manuscript. All authors proofread and approved the final version of the manuscript.

## Acknowledgments

This work is supported in part by the National Institute of General Medical Sciences (GM136422, S10OD026825), the National Institute of Allergy and Infectious Diseases (AI134678), and the National Science Foundation (IIS1901191, DBI2030790). This work used the XSEDE clusters [57] which is supported by the National Science Foundation (ACI1548562) for providing computational resources.

