## Supporting Information for "Identifying zoonotic origin of SARS-CoV-2 by modeling the binding affinity between Spike receptor-binding domain and host ACE2"

#### Table of Contents

##### Supporting Tables

**Table S1**. 30 ACE2 proteins that were excluded from analysis due to inaccurate annotations (shown as 'X') in their sequences.
**Table S2**. Six ACE2 proteins with five or more missing interface residues.
**Table S3**. 285 ACE2 proteins used for detailed analysis in this work.
**Table S4**. Binding energy and the three kinds of sequence identity for the 285 selected species.
**Table S5**. Comparison of experimental and predicted N-glycosylation sites on hACE2.
**Table S6**. Potential N-glycosylation sites in the 285 selected ACE2 proteins.
**Table S7**. Comparison of the interface residues in the 285 ACE2 proteins.

##### Supporting Figures

**Fig. S1**. The distribution of protein length, and sequence identity for all, interface, and the five key residues for 285 ACE2 orthologs.
**Fig. S2**. Matthews correlation coefficient for classifying experimentally determined effective ACE2 receptors from the less effective ones by the binding energy calculated from 500 models.
**Fig. S3**. Matthews correlation coefficient for classifying experimentally determined effective ACE2 receptors from the less effective ones by sequence identity in terms of all (a), interface (b), and key (c) residues.
**Fig. S3**. Matthews correlation coefficient for classifying experimentally determined effective ACE2 receptors from the less effective ones.
**Fig. S4**. Binding score of 500 models for the hACE2/S-RBD complex.
**Fig. S5**. Matthews correlation coefficient for classifying experimentally determined effective ACE2 receptors from the less effective ones by the binding energy calculated from the first model.

#### Supporting Tables

**Table S1**. 30 ACE2 proteins that were excluded from analysis due to inaccurate annotations (shown as 'X') in their sequences.

| **Accession ID** | **Species** | **Class** | **Common name** | **Length (aa)** | **Remark*** |
|---|---|---|---|---|---|
| XP_005231984.2 | *Falco peregrinus* | Aves | Peregrine falcon | 824 | Low |
| XP_005426221.1 | *Geospiza fortis* | Aves | Medium ground-finch | 809 | Low |
| XP_009478920.1 | *Pelecanus crispus* | Aves | Dalmatian pelican | 809 | Low |
| XP_009509070.1 | *Phalacrocorax carbo* | Aves | Great cormorant | 812 | |
| XP_009563864.1 | *Cuculus canorus* | Aves | Common cuckoo | 808 | Low |
| XP_009638257.1 | *Egretta garzetta* | Aves | Little egret | 806 | Low |
| XP_009703695.1 | *Cariama cristata* | Aves | Red-legged seriema | 806 | Low |
| XP_009867056.1 | *Apaloderma vittatum* | Aves | Bar-tailed trogon | 791 | Low |
| XP_009909849.1 | *Picoides pubescens* | Aves | Downy woodpecker | 387 | Partial |
| XP_009954393.1 | *Leptosomus discolor* | Aves | Madagascar cuckoo roller | 809 | Low |
| XP_009978415.1 | *Tauraco erythrolophus* | Aves | Red-crested turaco | 703 | Low/Partial |
| XP_010012481.1 | *Nestor notabilis* | Aves | Kea | 618 | Low |
| XP_010084373.1 | *Pterocles gutturalis* | Aves | Yellow-throated sandgrouse | 631 | Partial |
| XP_010169238.1 | *Antrostomus carolinensis* | Aves | Chuck-will's-widow | 807 | |
| XP_010206054.1 | *Colius striatus* | Aves | Speckled mousebird | 656 | Low |
| XP_010217584.1 | *Tinamus guttatus* | Aves | White-throated tinamou | 805 | Low |
| XP_013039300.1 | *Anser cygnoides domesticus* | Aves | Domestic goose | 805 | Low |
| XP_013888928.1 | *Austrofundulus limnaeus* | Actinopterygii | | 819 | |
| XP_017367865.1 | *Cebus capucinus imitator* | Mammalia | Panamanian white-faced capuchin | 805 | |
| XP_020465646.1 | *Monopterus albus* | Actinopterygii | Asian swamp eel | 812 | |
| XP_020863153.1 | *Phascolarctos cinereus* | Mammalia | Koala | 807 | Low |
| XP_021154486.1 | *Columba livia* | Aves | Rock dove | 799 | Low |
| XP_023417808.1 | *Cavia porcellus* | Mammalia | Domestic guinea pig | 813 | Low |
| XP_023679669.1 | *Paramormyrops kingsleyae* | Actinopterygii | | 805 | Low |
| XP_023998967.1 | *Salvelinus alpinus* | Actinopterygii | Arctic char | 789 | Low/Partial |
| XP_025976569.1 | *Dromaius novaehollandiae* | Aves | Emu | 804 | |
| XP_026530754.1 | *Notechis scutatus* | Reptilia | Mainland tiger snake | 828 | Low |
| XP_030271236.1 | *Sparus aurata* | Actinopterygii | Gilthead seabream | 816 | Low |
| XP_031814825.1 | *Sarcophilus harrisii* | Mammalia | Tasmanian devil | 806 | Low |
| XP_033056809.1 | *Trachypithecus francoisi* | Mammalia | Francoiss langur | 805 | Low |

*: Low, low-quality sequence; Partial, incomplete ACE2 sequence; Low/Partial, both.

**Table S2**. Six ACE2 proteins with five or more missing interface residues.

| Accession ID | Species | Class | Common name | Length (aa) | Binding (EEU) | Remark* |
|---|---|---|---|---|---|---|
| XP_009082150.1 | *Acanthisitta chloris* | Aves | Rifleman | 609 | -18.02 | Partial |
| XP_013805736.1 | *Apteryx mantelli mantelli* | Aves | North island brown kiwi | 777 | -20.07 | |
| XP_014062928.1 | *Salmo salar* | Actinopterygii | Atlantic salmon | 686 | -13.70 | |
| XP_017744069.1 | *Rhinopithecus bieti* | Mammalia | Black snub-nosed monkey | 724 | -20.36 | |
| XP_030886750.1 | *Leptonychotes weddellii* | Mammalia | Weddell seal | 344 | -16.90 | |
| XP_032835032.1 | *Petromyzon marinus* | Hyperoartia | Sea lamprey | 849 | -20.19 | |

*: partial, incomplete ACE2 sequence.

**Table S3**. 285 ACE2 proteins used for detailed analysis in this work.

| Accession ID | Species | Common name | Class | Length (aa) |
|---|---|---|---|---|
| AAX63775.1 | *Paguma larvata* | Masked palm civet | Mammalia | 805 |
| ABW16956.1 | *Nyctereutes procyonoides* | Raccoon dog | Mammalia | 804 |
| AGZ48803.1 | *Rhinolophus sinicus* | Chinese rufous horseshoe bat | Mammalia | 805 |
| NP_001012006.1 | *Rattus norvegicus* | Brown rat | Mammalia | 805 |
| NP_001116542.1 | *Sus scrofa* | Pig | Mammalia | 805 |
| NP_001124604.1 | *Pongo abelii* | Sumatran orangutan | Mammalia | 805 |
| NP_001129168.1 | *Macaca mulatta* | Rhesus macaque | Mammalia | 805 |
| NP_001158732.1 | *Canis lupus familiaris* | Dog | Mammalia | 804 |
| NP_001277036.1 | *Capra hircus* | Goat | Mammalia | 804 |
| NP_001297119.1 | *Mustela putorius furo* | Domestic ferret | Mammalia | 805 |
| NP_001358344.1 | *Homo sapiens* | Human | Mammalia | 805 |
| NP_081562.2 | *Mus musculus* | House mouse | Mammalia | 805 |
| XP_001490241.1 | *Equus caballus* | Horse | Mammalia | 805 |
| XP_001515597.2 | *Ornithorhynchus anatinus* | Platypus | Mammalia | 806 |
| XP_002194303.4 | *Taeniopygia guttata* | Zebra finch | Aves | 811 |
| XP_002719891.1 | *Oryctolagus cuniculus* | Rabbit | Mammalia | 805 |
| XP_002938293.2 | *Xenopus tropicalis* | Tropical clawed frog | Amphibia | 862 |
| XP_003261132.2 | *Nomascus leucogenys* | Northern white-cheeked gibbon | Mammalia | 805 |
| XP_003445853.2 | *Oreochromis niloticus* | Nile tilapia | Actinopterygii | 821 |
| XP_003503283.1 | *Cricetulus griseus* | Chinese hamster | Mammalia | 805 |
| XP_003791912.1 | *Otolemur garnettii* | Small-eared galago | Mammalia | 805 |
| XP_004269705.1 | *Orcinus orca* | Killer whale | Mammalia | 804 |
| XP_004386381.1 | *Trichechus manatus latirostris* | Florida manatee | Mammalia | 800 |
| XP_004415448.1 | *Odobenus rosmarus divergens* | Walrus | Mammalia | 732 |
| XP_004435206.1 | *Ceratotherium simum simum* | Southern white rhinoceros | Mammalia | 805 |
| XP_004449124.1 | *Dasypus novemcinctus* | Nine-banded armadillo | Mammalia | 804 |
| XP_004543482.1 | *Maylandia zebra* | Zebra mbuna | Actinopterygii | 803 |
| XP_004597549.2 | *Ochotona princeps* | American pika | Mammalia | 808 |
| XP_004612266.1 | *Sorex araneus* | Eurasian common shrew | Mammalia | 803 |
| XP_004671523.1 | *Jaculus jaculus* | Lesser egyptian jerboa | Mammalia | 805 |
| XP_004710002.1 | *Echinops telfairi* | Small madagascar hedgehog | Mammalia | 798 |
| XP_004866157.1 | *Heterocephalus glaber* | Naked mole-rat | Mammalia | 805 |
| XP_005037422.1 | *Ficedula albicollis* | Collared flycatcher | Aves | 810 |
| XP_005074266.1 | *Mesocricetus auratus* | Golden hamster | Mammalia | 805 |
| XP_005151516.2 | *Melopsittacus undulatus* | Budgerigar | Aves | 809 |
| XP_005169416.1 | *Danio rerio* | Zebrafish | Actinopterygii | 818 |
| XP_005228485.1 | *Bos taurus* | Cattle | Mammalia | 811 |
| XP_005316051.3 | *Ictidomys tridecemlineatus* | Thirteen-lined ground squirrel | Mammalia | 817 |
| XP_005358818.1 | *Microtus ochrogaster* | Prairie vole | Mammalia | 804 |
| XP_005443093.2 | *Falco cherrug* | Saker falcon | Aves | 813 |
| XP_005491832.2 | *Zonotrichia albicollis* | White-throated sparrow | Aves | 811 |
| XP_005516712.1 | *Pseudopodoces humilis* | Tibetan ground jay | Aves | 809 |
| XP_005593094.1 | *Macaca fascicularis* | Crab-eating macaque | Mammalia | 805 |
| XP_005724169.1 | *Pundamilia nyererei* |  | Actinopterygii | 803 |
| XP_005799835.1 | *Xiphophorus maculatus* | Southern platyfish | Actinopterygii | 808 |
| XP_005903173.1 | *Bos mutus* | Wild yak | Mammalia | 804 |
| XP_005943362.1 | *Haplochromis burtoni* | Burtons mouthbrooder | Actinopterygii | 803 |
| XP_005997915.2 | *Latimeria chalumnae* | Coelacanth | Sarcopterygii | 859 |
| XP_006041602.1 | *Bubalus bubalis* | Water buffalo | Mammalia | 803 |
| XP_006122891.1 | *Pelodiscus sinensis* | Chinese soft-shelled turtle | Reptilia | 808 |
| XP_006164754.1 | *Tupaia chinensis* | Chinese tree shrew | Mammalia | 805 |
| XP_006194263.1 | *Camelus ferus* | Wild bactrian camel | Mammalia | 805 |
| XP_006212709.1 | *Vicugna pacos* | Alpaca | Mammalia | 805 |
| XP_006639185.1 | *Lepisosteus oculatus* | Spotted gar | Actinopterygii | 809 |
| XP_006775273.1 | *Myotis davidii* | David's myotis | Mammalia | 819 |
| XP_006780474.1 | *Neolamprologus brichardi* | Princess cichlid | Actinopterygii | 803 |
| XP_006835673.1 | *Chrysochloris asiatica* | Cape golden mole | Mammalia | 799 |
| XP_006892457.1 | *Elephantulus edwardii* | Cape elephant shrew | Mammalia | 798 |

| | | | | |
|---|---|---|---|---|
| XP_006911709.1 | *Pteropus alecto* | Black flying fox | Mammalia | 805 |
| XP_006973269.1 | *Peromyscus maniculatus bairdii* | Deer mouse | Mammalia | 805 |
| XP_007070561.1 | *Chelonia mydas* | Green sea turtle | Reptilia | 811 |
| XP_007090142.1 | *Panthera tigris altaica* | Amur tiger | Mammalia | 797 |
| XP_007431942.2 | *Python bivittatus* | Burmese python | Reptilia | 827 |
| XP_007466389.1 | *Lipotes vexillifer* | Yangtze river dolphin | Mammalia | 804 |
| XP_007500935.1 | *Monodelphis domestica* | Gray short-tailed opossum | Mammalia | 806 |
| XP_007538670.1 | *Erinaceus europaeus* | Western european hedgehog | Mammalia | 804 |
| XP_007560208.1 | *Poecilia formosa* | Amazon molly | Actinopterygii | 808 |
| XP_007889845.1 | *Callorhinchus milii* | Ghost shark | Chondrichthyes | 836 |
| XP_007951028.1 | *Orycteropus afer* | Aardvark | Mammalia | 799 |
| XP_007989304.1 | *Chlorocebus sabaeus* | Green monkey | Mammalia | 805 |
| XP_008062810.1 | *Carlito syrichta* | Philippine tarsier | Mammalia | 805 |
| XP_008105455.1 | *Anolis carolinensis* | Green anole | Reptilia | 814 |
| XP_008153150.1 | *Eptesicus fuscus* | Big brown bat | Mammalia | 811 |
| XP_008290762.1 | *Stegastes partitus* | Bicolor damselfish | Actinopterygii | 807 |
| XP_008402714.1 | *Poecilia reticulata* | Guppy | Actinopterygii | 809 |
| XP_008492997.2 | *Calypte anna* | Annas hummingbird | Aves | 805 |
| XP_008542995.1 | *Equus przewalskii* | Mongolian wild horse | Mammalia | 805 |
| XP_008694637.1 | *Ursus maritimus* | Polar bear | Mammalia | 790 |
| XP_008839098.1 | *Nannospalax galili* | Israeli mole rat | Mammalia | 804 |
| XP_008937519.1 | *Merops nubicus* | Northern carmine bee-eater | Aves | 799 |
| XP_008972428.2 | *Pan paniscus* | Bonobo | Mammalia | 805 |
| XP_008987241.1 | *Callithrix jacchus* | Common marmoset | Mammalia | 805 |
| XP_009087922.1 | *Serinus canaria* | Island canary | Aves | 809 |
| XP_009275140.1 | *Aptenodytes forsteri* | Emperor penguin | Aves | 809 |
| XP_009323767.1 | *Pygoscelis adeliae* | Adelie penguin | Aves | 798 |
| XP_009474590.1 | *Nipponia nippon* | Crested ibis | Aves | 809 |
| XP_009574896.1 | *Fulmarus glacialis* | Northern fulmar | Aves | 632 |
| XP_009667495.1 | *Struthio camelus australis* | South african ostrich | Aves | 808 |
| XP_009816127.1 | *Gavia stellata* | Red-throated loon | Aves | 809 |
| XP_009887331.1 | *Charadrius vociferus* | Killdeer | Aves | 809 |
| XP_009925641.1 | *Haliaeetus albicilla* | White-tailed eagle | Aves | 629 |
| XP_009938970.1 | *Opisthocomus hoazin* | Hoatzin | Aves | 824 |
| XP_009992128.1 | *Chaetura pelagica* | Chimney swift | Aves | 806 |
| XP_010120523.1 | *Chlamydotis macqueenii* | Macqueen's bustard | Aves | 808 |
| XP_010136813.1 | *Buceros rhinoceros silvestris* | Rhinoceros hornbill | Aves | 794 |
| XP_010156467.1 | *Eurypyga helias* | Sunbittern | Aves | 809 |
| XP_010178703.1 | *Mesitornis unicolor* | Brown roatelo | Aves | 809 |
| XP_010290019.1 | *Phaethon lepturus* | White-tailed tropicbird | Aves | 809 |
| XP_010334925.1 | *Saimiri boliviensis* | Black-capped squirrel monkey | Mammalia | 805 |
| XP_010364367.2 | *Rhinopithecus roxellana* | Golden snub-nosed monkey | Mammalia | 805 |
| XP_010392735.2 | *Corvus cornix cornix* | Hooded crow | Aves | 811 |
| XP_010579828.1 | *Haliaeetus leucocephalus* | Bald eagle | Aves | 745 |
| XP_010643477.1 | *Fukomys damarensis* | Damaraland mole-rat | Mammalia | 805 |
| XP_010730146.1 | *Larimichthys crocea* | Large yellow croaker | Actinopterygii | 810 |
| XP_010790455.1 | *Notothenia coriiceps* | Black rockcod | Actinopterygii | 806 |
| XP_010833001.1 | *Bison bison bison* | Plains bison | Mammalia | 431 |
| XP_010884777.2 | *Esox lucius* | Northern pike | Actinopterygii | 812 |
| XP_010966303.1 | *Camelus bactrianus* | Bactrian camel | Mammalia | 805 |
| XP_010991717.1 | *Camelus dromedarius* | Arabian camel | Mammalia | 805 |
| XP_011361275.1 | *Pteropus vampyrus* | Large flying fox | Mammalia | 804 |
| XP_011733505.1 | *Macaca nemestrina* | Pig-tailed macaque | Mammalia | 805 |
| XP_011850923.1 | *Mandrillus leucophaeus* | Drill | Mammalia | 805 |
| XP_011891198.1 | *Cercocebus atys* | Sooty mangabey | Mammalia | 805 |
| XP_011961657.1 | *Ovis aries* | Sheep | Mammalia | 804 |
| XP_012290105.1 | *Aotus nancymaae* | Ma's night monkey | Mammalia | 805 |
| XP_012494185.1 | *Propithecus coquereli* | Coquerel's sifaka | Mammalia | 826 |
| XP_012585871.1 | *Condylura cristata* | Star-nosed mole | Mammalia | 800 |
| XP_012887572.1 | *Dipodomys ordii* | Ords kangaroo rat | Mammalia | 805 |

| | | | | |
|---|---|---|---|---|
| XP_012949915.2 | *Anas platyrhynchos* | Duck | Aves | 805 |
| XP_013362428.1 | *Chinchilla lanigera* | Long-tailed chinchilla | Mammalia | 805 |
| XP_013926936.1 | *Thamnophis sirtalis* | Common garter snake | Reptilia | 461 |
| XP_014399780.1 | *Myotis brandtii* | Brandts bat | Mammalia | 819 |
| XP_014713133.1 | *Equus asinus* | Donkey | Mammalia | 783 |
| XP_014731370.1 | *Sturnus vulgaris* | Common starling | Aves | 810 |
| XP_014815705.1 | *Calidris pugnax* | Ruff | Aves | 809 |
| XP_014837025.1 | *Poecilia mexicana* | Cave molly | Actinopterygii | 808 |
| XP_014895313.1 | *Poecilia latipinna* | Sailfin molly | Actinopterygii | 808 |
| XP_015226730.1 | *Cyprinodon variegatus* | Sheepshead minnow | Actinopterygii | 819 |
| XP_015273067.1 | *Gekko japonicus* | Schlegel's japanese gecko | Reptilia | 816 |
| XP_015343540.1 | *Marmota marmota* | Alpine marmot | Mammalia | 817 |
| XP_015486815.1 | *Parus major* | Great tit | Aves | 814 |
| XP_015742063.1 | *Coturnix japonica* | Japanese quail | Aves | 807 |
| XP_015808977.1 | *Nothobranchius furzeri* | Turquoise killifish | Actinopterygii | 805 |
| XP_015974412.1 | *Rousettus aegyptiacus* | Egyptian rousette | Mammalia | 805 |
| XP_016058453.1 | *Miniopterus natalensis* | Natal long-fingered bat | Mammalia | 804 |
| XP_016345325.1 | *Sinocyclocheilus anshuiensis* | | Actinopterygii | 809 |
| XP_016422243.1 | *Sinocyclocheilus rhinocerous* | | Actinopterygii | 777 |
| XP_016798468.1 | *Pan troglodytes* | Chimpanzee | Mammalia | 805 |
| XP_016887914.1 | *Cynoglossus semilaevis* | Tongue sole | Actinopterygii | 802 |
| XP_017295385.1 | *Kryptolebias marmoratus* | Mangrove rivulus | Actinopterygii | 814 |
| XP_017313836.1 | *Ictalurus punctatus* | Channel catfish | Actinopterygii | 805 |
| XP_017505746.1 | *Manis javanica* | Malayan pangolin | Mammalia | 805 |
| XP_017550079.1 | *Pygocentrus nattereri* | Red-bellied piranha | Actinopterygii | 804 |
| XP_017583883.1 | *Corvus brachyrhynchos* | American crow | Aves | 758 |
| XP_017667729.1 | *Lepidothrix coronata* | Blue-crowned manakin | Aves | 811 |
| XP_017939494.2 | *Manacus vitellinus* | Golden-collared manakin | Aves | 811 |
| XP_018418558.1 | *Nanorana parkeri* | High himalaya frog | Amphibia | 773 |
| XP_018539189.1 | *Lates calcarifer* | Barramundi perch | Actinopterygii | 807 |
| XP_018584732.1 | *Scleropages formosus* | Asian bonytongue | Actinopterygii | 809 |
| XP_018874749.1 | *Gorilla gorilla* | Western gorilla | Mammalia | 805 |
| XP_019273508.1 | *Panthera pardus* | Leopard | Mammalia | 805 |
| XP_019350687.1 | *Alligator mississippiensis* | American alligator | Reptilia | 805 |
| XP_019381060.1 | *Gavialis gangeticus* | Gharial | Reptilia | 803 |
| XP_019384826.1 | *Crocodylus porosus* | Saltwater crocodile | Reptilia | 803 |
| XP_019467554.1 | *Meleagris gallopavo* | Wild turkey | Aves | 861 |
| XP_019522936.1 | *Hipposideros armiger* | Great roundleaf bat | Mammalia | 806 |
| XP_019742561.1 | *Hippocampus comes* | Tiger tail seahorse | Actinopterygii | 805 |
| XP_019781177.2 | *Tursiops truncatus* | Atlantic bottle-nosed dolphin | Mammalia | 804 |
| XP_019811719.1 | *Bos indicus* | Zebu cattle | Mammalia | 811 |
| XP_019935235.1 | *Paralichthys olivaceus* | Bastard halibut | Actinopterygii | 721 |
| XP_020140826.1 | *Microcebus murinus* | Gray mouse lemur | Mammalia | 851 |
| XP_020493627.1 | *Labrus bergylta* | Ballan wrasse | Actinopterygii | 806 |
| XP_020642422.1 | *Pogona vitticeps* | Central bearded dragon | Reptilia | 821 |
| XP_020768965.1 | *Odocoileus virginianus texanus* | White-tailed deer | Mammalia | 679 |
| XP_020781598.1 | *Boleophthalmus pectinirostris* | Great blue-spotted mudskipper | Actinopterygii | 807 |
| XP_021009138.1 | *Mus caroli* | Ryukyu mouse | Mammalia | 805 |
| XP_021043935.1 | *Mus pahari* | Shrew mouse | Mammalia | 805 |
| XP_021178197.1 | *Fundulus heteroclitus* | Mummichog | Actinopterygii | 809 |
| XP_021240731.1 | *Numida meleagris* | Helmeted guineafowl | Aves | 810 |
| XP_021388026.1 | *Lonchura striata domestica* | Bengalese finch | Aves | 809 |
| XP_021433278.1 | *Oncorhynchus mykiss* | Rainbow trout | Actinopterygii | 807 |
| XP_021536480.1 | *Neomonachus schauinslandi* | Hawaiian monk seal | Mammalia | 805 |
| XP_021788732.1 | *Papio anubis* | Olive baboon | Mammalia | 805 |
| XP_022063988.1 | *Acanthochromis polyacanthus* | Spiny chromis | Actinopterygii | 817 |
| XP_022374078.1 | *Enhydra lutris kenyoni* | Sea otter | Mammalia | 805 |
| XP_022418360.1 | *Delphinapterus leucas* | Beluga whale | Mammalia | 804 |
| XP_022523929.1 | *Astyanax mexicanus* | Mexican tetra | Actinopterygii | 805 |
| XP_022605054.1 | *Seriola dumerili* | Greater amberjack | Actinopterygii | 807 |

| | | | | |
|---|---|---|---|---|
| XP_023054821.1 | *Piliocolobus tephrosceles* | Ugandan red colobus | Mammalia | 805 |
| XP_023104564.1 | *Felis catus* | Domestic cat | Mammalia | 807 |
| XP_023124156.1 | *Amphiprion ocellaris* | Clown anemonefish | Actinopterygii | 807 |
| XP_023257445.1 | *Seriola lalandi dorsalis* | California yellowtail | Actinopterygii | 807 |
| XP_023410960.1 | *Loxodonta africana* | African savanna elephant | Mammalia | 800 |
| XP_023575315.1 | *Octodon degus* | Common degu | Mammalia | 831 |
| XP_023609437.1 | *Myotis lucifugus* | Little brown bat | Mammalia | 819 |
| XP_023774184.1 | *Cyanistes caeruleus* | Eurasian blue tit | Aves | 814 |
| XP_023964517.1 | *Chrysemys picta* | Painted turtle | Reptilia | 828 |
| XP_023971279.1 | *Physeter macrocephalus* | Sperm whale | Mammalia | 804 |
| XP_024150631.1 | *Oryzias melastigma* | Marine medaka | Actinopterygii | 819 |
| XP_024425698.1 | *Desmodus rotundus* | Vampire bat | Mammalia | 804 |
| XP_024599894.1 | *Neophocaena asiaeorientalis* | Yangtze finless porpoise | Mammalia | 804 |
| XP_025066628.1 | *Alligator sinensis* | Chinese alligator | Reptilia | 803 |
| XP_025227847.1 | *Theropithecus gelada* | Gelada | Mammalia | 805 |
| XP_025292925.1 | *Canis lupus dingo* | Dingo | Mammalia | 804 |
| XP_025713397.1 | *Callorhinus ursinus* | Northern fur seal | Mammalia | 806 |
| XP_025790417.1 | *Puma concolor* | Puma | Mammalia | 805 |
| XP_025842512.1 | *Vulpes vulpes* | Red fox | Mammalia | 804 |
| XP_025891105.1 | *Nothoprocta perdicaria* | Chilean tinamou | Aves | 807 |
| XP_025942946.1 | *Apteryx rowi* | Okarito brown kiwi | Aves | 808 |
| XP_026020155.1 | *Astatotilapia calliptera* | Eastern happy | Actinopterygii | 803 |
| XP_026131313.1 | *Carassius auratus* | Goldfish | Actinopterygii | 812 |
| XP_026175949.1 | *Mastacembelus armatus* | Zig-zag eel | Actinopterygii | 807 |
| XP_026233431.1 | *Anabas testudineus* | Climbing perch | Actinopterygii | 806 |
| XP_026252505.1 | *Urocitellus parryii* | Arctic ground squirrel | Mammalia | 817 |
| XP_026333865.1 | *Ursus arctos horribilis* | Grizzly bear | Mammalia | 805 |
| XP_026570054.1 | *Pseudonaja textilis* | Eastern brown snake | Reptilia | 828 |
| XP_026705725.1 | *Athene cunicularia* | Burrowing owl | Aves | 834 |
| XP_026803610.1 | *Pangasianodon hypophthalmus* | Striped catfish | Actinopterygii | 805 |
| XP_026867211.2 | *Electrophorus electricus* | Electric eel | Actinopterygii | 805 |
| XP_026910297.1 | *Acinonyx jubatus* | Cheetah | Mammalia | 805 |
| XP_026951598.1 | *Lagenorhynchus obliquidens* | Pacific white-sided dolphin | Mammalia | 804 |
| XP_027024524.1 | *Tachysurus fulvidraco* | Yellow catfish | Actinopterygii | 806 |
| XP_027389727.1 | *Bos indicus x Bos taurus* | Hybrid cattle | Mammalia | 811 |
| XP_027465353.1 | *Zalophus californianus* | California sea lion | Mammalia | 806 |
| XP_027494818.1 | *Corapipo altera* | White-ruffed manakin | Aves | 811 |
| XP_027544864.1 | *Neopelma chrysocephalum* | Saffron-crested tyrant-manakin | Aves | 811 |
| XP_027593974.1 | *Pipra filicauda* | Wire-tailed manakin | Aves | 811 |
| XP_027691156.1 | *Vombatus ursinus* | Common wombat | Mammalia | 809 |
| XP_027757151.1 | *Empidonax traillii* | Willow flycatcher | Aves | 809 |
| XP_027802308.1 | *Marmota flaviventris* | Yellow-bellied marmot | Mammalia | 817 |
| XP_027871671.1 | *Xiphophorus couchianus* | Monterrey platyfish | Actinopterygii | 808 |
| XP_027970822.1 | *Eumetopias jubatus* | Steller sea lion | Mammalia | 806 |
| XP_028020351.1 | *Balaenoptera acutorostrata scammoni* | North pacific minke whale | Mammalia | 804 |
| XP_028257887.1 | *Parambassis ranga* | Indian glassy fish | Actinopterygii | 808 |
| XP_028297875.1 | *Gouania willdenowi* | Blunt-snouted clingfish | Actinopterygii | 809 |
| XP_028378317.1 | *Phyllostomus discolor* | Pale spear-nosed bat | Mammalia | 804 |
| XP_028441363.1 | *Perca flavescens* | Yellow perch | Actinopterygii | 820 |
| XP_028617961.1 | *Grammomys surdaster* | African woodland thicket rat | Mammalia | 805 |
| XP_028655640.1 | *Erpetoichthys calabaricus* | Rope fish | Actinopterygii | 799 |
| XP_028743609.1 | *Peromyscus leucopus* | White-footed mouse | Mammalia | 805 |
| XP_028837781.1 | *Denticeps clupeoides* | Denticle herring | Actinopterygii | 810 |
| XP_028999570.1 | *Betta splendens* | Siamese fighting fish | Actinopterygii | 814 |
| XP_029095804.1 | *Monodon monoceros* | Narwhal | Mammalia | 804 |
| XP_029140508.1 | *Protobothrops mucrosquamatus* | Protobothrops mucrosquamatus | Reptilia | 861 |
| XP_029283581.1 | *Cottoperca gobio* | Frogmouth | Actinopterygii | 807 |
| XP_029354066.1 | *Echeneis naucrates* | Live sharksucker | Actinopterygii | 814 |
| XP_029459086.1 | *Rhinatrema bivittatum* | Two-lined caecilian | Amphibia | 815 |
| XP_029702274.1 | *Takifugu rubripes* | Torafugu | Actinopterygii | 806 |

| | | | | |
|---|---|---|---|---|
| XP_029786256.1 | *Suricata suricatta* | Meerkat | Mammalia | 805 |
| XP_029855025.1 | *Aquila chrysaetos chrysaetos* | Golden eagle | Aves | 809 |
| XP_029904152.1 | *Myripristis murdjan* | Pinecone soldierfish | Actinopterygii | 806 |
| XP_029949252.1 | *Salarias fasciatus* | Jewelled blenny | Actinopterygii | 816 |
| XP_030058174.1 | *Microcaecilia unicolor* | Tiny Cayenne Caecilian | Amphibia | 811 |
| XP_030160839.1 | *Lynx canadensis* | Canada lynx | Mammalia | 805 |
| XP_030232530.1 | *Gadus morhua* | Atlantic cod | Actinopterygii | 818 |
| XP_030332639.1 | *Strigops habroptila* | Kakapo | Aves | 811 |
| XP_030407881.1 | *Gopherus evgoodei* | Goodes thornscrub tortoise | Reptilia | 808 |
| XP_030582139.1 | *Archocentrus centrarchus* | Flier cichlid | Actinopterygii | 803 |
| XP_030627971.1 | *Chanos chanos* | Milkfish | Actinopterygii | 804 |
| XP_030703991.1 | *Globicephala melas* | Long-finned pilot whale | Mammalia | 804 |
| XP_030811385.1 | *Camarhynchus parvulus* | Small tree finch | Aves | 809 |
| XP_031162227.1 | *Sander lucioperca* | Pike-perch | Actinopterygii | 808 |
| XP_031226742.1 | *Mastomys coucha* | Southern multimammate mouse | Mammalia | 806 |
| XP_031414786.1 | *Clupea harengus* | Atlantic herring | Actinopterygii | 821 |
| XP_031451919.1 | *Phasianus colchicus* | Common pheasant | Aves | 808 |
| XP_031584810.1 | *Oreochromis aureus* | Israeli tilapia | Actinopterygii | 821 |
| XP_031702716.1 | *Anarrhichthys ocellatus* | Wolf eel | Actinopterygii | 815 |
| XP_031956594.1 | *Corvus moneduloides* | New caledonian crow | Aves | 811 |
| XP_032058386.1 | *Aythya fuligula* | Tufted duck | Aves | 805 |
| XP_032082934.1 | *Thamnophis elegans* | Western terrestrial garter snake | Reptilia | 828 |
| XP_032141854.1 | *Sapajus apella* | Tufted capuchin | Mammalia | 805 |
| XP_032187677.1 | *Mustela erminea* | Ermine | Mammalia | 805 |
| XP_032245506.1 | *Phoca vitulina* | Harbor seal | Mammalia | 805 |
| XP_032355526.1 | *Etheostoma spectabile* | Orangethroat darter | Actinopterygii | 808 |
| XP_032417398.1 | *Xiphophorus hellerii* | Green swordtail | Actinopterygii | 808 |
| XP_032476001.1 | *Phocoena sinus* | Vaquita | Mammalia | 804 |
| XP_032536093.1 | *Chiroxiphia lanceolata* | Lance-tailed manakin | Aves | 811 |
| XP_032612508.1 | *Hylobates moloch* | Silvery gibbon | Mammalia | 805 |
| XP_032631197.1 | *Chelonoidis abingdonii* | Pinta island tortoise | Reptilia | 808 |
| XP_032736028.1 | *Lontra canadensis* | Northern american river otter | Mammalia | 805 |
| XP_032746145.1 | *Rattus rattus* | Black rat | Mammalia | 793 |
| XP_032865981.1 | *Tyto alba* | Western barn owl | Aves | 809 |
| XP_032888812.1 | *Amblyraja radiata* | Thorny skate | Chondrichthyes | 773 |
| XP_032907631.1 | *Catharus ustulatus* | Swainsons thrush | Aves | 872 |
| XP_032963186.1 | *Rhinolophus ferrumequinum* | Greater horseshoe bat | Mammalia | 805 |
| XP_033806113.1 | *Geotrypetes seraphini* | Gaboon caecilian | Amphibia | 820 |
| XP_034290793.1 | *Pantherophis guttatus* | Corn snake | Reptilia | 828 |
| XP_034341939.1 | *Arvicanthis niloticus* | African grass rat | Mammalia | 805 |
| XP_034505781.1 | *Ailuropoda melanoleuca* | Giant panda | Mammalia | 705 |
| XP_034612285.1 | *Trachemys scripta elegans* | Red-eared slider | Reptilia | 828 |
| XP_034852450.1 | *Mirounga leonina* | Southern elephant seal | Mammalia | 805 |
| XP_034971718.1 | *Zootoca vivipara* | Viviparous lizard | Reptilia | 806 |
| XP_035182951.1 | *Oxyura jamaicensis* | Ruddy duck | Aves | 808 |
| XP_035398116.1 | *Cygnus atratus* | Black Swan | Aves | 808 |
| XP_416822.2 | *Gallus gallus* | Chicken | Aves | 808 |

**Table S4**. Binding energy and the three kinds of sequence identity for the 285 selected species.

| Species | Order | Common name | Binding (EEU) | Sequence Identity (%) | | |
|---|---|---|---|---|---|---|
| | | | | All | Interface | Key |
| *Pongo abelii* | Primates | Sumatran orangutan | -56.21 | 98.1 | 100 | 100 |
| *Pan paniscus* | Primates | Bonobo | -55.99 | 99.0 | 100 | 100 |
| *Nomascus leucogenys* | Primates | Northern white-cheeked gibbon | -55.84 | 97.8 | 100 | 100 |
| *Gorilla gorilla* | Primates | Western gorilla | -55.84 | 99.0 | 100 | 100 |
| *Papio anubis* | Primates | Olive baboon | -55.77 | 95.3 | 100 | 100 |
| *Hylobates moloch* | Primates | Silvery gibbon | -55.73 | 98.0 | 100 | 100 |
| *Chlorocebus sabaeus* | Primates | Green monkey | -55.67 | 94.8 | 100 | 100 |
| *Macaca nemestrina* | Primates | Pig-tailed macaque | -55.42 | 95.3 | 100 | 100 |
| *Macaca fascicularis* | Primates | Crab-eating macaque | -55.38 | 95.2 | 100 | 100 |
| *Theropithecus gelada* | Primates | Gelada | -55.29 | 95.3 | 100 | 100 |
| *Macaca mulatta* | Primates | Rhesus macaque | -55.24 | 94.9 | 100 | 100 |
| *Cercocebus atys* | Primates | Sooty mangabey | -55.19 | 95.2 | 100 | 100 |
| *Homo sapiens* | Primates | Human | -55.16 | 100 | 100 | 100 |
| *Rhinopithecus roxellana* | Primates | Golden snub-nosed monkey | -55.09 | 95.3 | 100 | 100 |
| *Pan troglodytes* | Primates | Chimpanzee | -54.97 | 99.0 | 100 | 100 |
| *Mandrillus leucophaeus* | Primates | Drill | -54.94 | 94.9 | 100 | 100 |
| *Piliocolobus tephrosceles* | Primates | Ugandan red colobus | -54.79 | 95.0 | 100 | 100 |
| *Mesocricetus auratus* | Rodentia | Golden hamster | -53.84 | 84.5 | 90 | 80 |
| *Cricetulus griseus* | Rodentia | Chinese hamster | -53.77 | 84.3 | 90 | 80 |
| *Nannospalax galili* | Rodentia | Northern israeli blind subterranean mole rat | -53.69 | 82.7 | 90 | 80 |
| *Eumetopias jubatus* | Carnivora | Steller sea lion | -53.47 | 83.3 | 65 | 60 |
| *Propithecus coquereli* | Primates | Coquerel's sifaka | -53.35 | 83.5 | 95 | 80 |
| *Callorhinus ursinus* | Carnivora | Northern fur seal | -52.98 | 83.3 | 65 | 60 |
| *Equus caballus* | Perissodactyla | Horse | -52.95 | 86.8 | 70 | 60 |
| *Equus przewalskii* | Perissodactyla | Mongolian wild horse | -52.95 | 87.0 | 70 | 60 |
| *Panthera tigris altaica* | Carnivora | Amur tiger | -52.93 | 85.7 | 80 | 60 |
| *Acinonyx jubatus* | Carnivora | Cheetah | -52.88 | 85.2 | 75 | 60 |
| *Capra hircus* | Artiodactyla | Goat | -52.86 | 81.8 | 85 | 80 |
| *Oryctolagus cuniculus* | Lagomorpha | Rabbit | -52.84 | 85.2 | 80 | 80 |
| *Bos mutus* | Artiodactyla | Wild yak | -52.83 | 81.5 | 85 | 80 |
| *Puma concolor* | Carnivora | Puma | -52.79 | 85.6 | 80 | 60 |
| *Heterocephalus glaber* | Rodentia | Naked mole-rat | -52.79 | 84.6 | 85 | 80 |
| *Bison bison bison* | Artiodactyla | Plains bison | -52.78 | 81.2 | 85 | 80 |
| *Panthera pardus* | Carnivora | Leopard | -52.74 | 85.5 | 80 | 60 |
| *Mustela erminea* | Carnivora | Ermine | -52.74 | 83.0 | 65 | 60 |
| *Phoca vitulina* | Carnivora | Harbor seal | -52.73 | 82.7 | 65 | 60 |
| *Bos taurus* | Artiodactyla | Cattle | -52.71 | 78.1 | 85 | 80 |
| *Bos indicus x Bos taurus* | Artiodactyla | Hybrid cattle | -52.71 | 78.1 | 85 | 80 |
| *Lontra canadensis* | Carnivora | Northern american river otter | -52.66 | 82.7 | 65 | 60 |
| *Odobenus rosmarus divergens* | Carnivora | Walrus | -52.62 | 83.1 | 65 | 60 |
| *Neomonachus schauinslandi* | Carnivora | Hawaiian monk seal | -52.56 | 82.5 | 65 | 60 |
| *Mustela putorius furo* | Carnivora | Domestic ferret | -52.55 | 82.6 | 65 | 60 |
| *Zalophus californianus* | Carnivora | California sea lion | -52.53 | 83.1 | 65 | 60 |
| *Bos indicus* | Artiodactyla | Zebu cattle | -52.47 | 78.1 | 85 | 80 |
| *Bubalus bubalis* | Artiodactyla | Water buffalo | -52.45 | 81.1 | 85 | 80 |
| *Jaculus jaculus* | Rodentia | Lesser egyptian jerboa | -52.44 | 84.8 | 75 | 80 |
| *Peromyscus maniculatus bairdii* | Rodentia | Deer mouse | -52.38 | 83.6 | 85 | 80 |
| *Felis catus* | Carnivora | Domestic cat | -52.33 | 84.6 | 80 | 60 |
| *Odocoileus virginianus texanus* | Artiodactyla | White-tailed deer | -52.24 | 79.1 | 85 | 80 |
| *Fukomys damarensis* | Rodentia | Damaraland mole-rat | -52.22 | 84.7 | 85 | 80 |
| *Lynx canadensis* | Carnivora | Canada lynx | -52.21 | 85.1 | 80 | 60 |
| *Ailuropoda melanoleuca* | Carnivora | Giant panda | -52.21 | 83.5 | 75 | 80 |
| *Peromyscus leucopus* | Rodentia | White-footed mouse | -52.15 | 83.1 | 85 | 80 |
| *Ursus arctos horribilis* | Carnivora | Grizzly bear | -52.13 | 83.7 | 75 | 80 |
| *Ovis aries* | Artiodactyla | Sheep | -52.09 | 81.8 | 85 | 80 |
| *Delphinapterus leucas* | Artiodactyla | Beluga whale | -51.98 | 81.6 | 85 | 80 |
| *Monodon monoceros* | Artiodactyla | Narwhal | -51.98 | 81.7 | 85 | 80 |
| *Phocoena sinus* | Artiodactyla | Vaquita | -51.95 | 81.3 | 85 | 80 |
| *Physeter macrocephalus* | Artiodactyla | Sperm whale | -51.94 | 82.8 | 85 | 80 |
| *Ursus maritimus* | Carnivora | Polar bear | -51.93 | 83.9 | 75 | 80 |
| *Neophocaena asiaeorientalis asiaeorientalis* | Artiodactyla | Yangtze finless porpoise | -51.85 | 81.1 | 85 | 80 |
| *Microtus ochrogaster* | Rodentia | Prairie vole | -51.76 | 83.8 | 75 | 80 |
| *Manis javanica* | Pholidota | Malayan pangolin | -51.73 | 84.8 | 65 | 60 |
| *Balaenoptera acutorostrata scammoni* | Artiodactyla | North pacific minke whale | -51.65 | 82.6 | 80 | 80 |
| *Marmota marmota* | Rodentia | Alpine marmot | -51.62 | 84.0 | 85 | 80 |
| *Vulpes vulpes* | Carnivora | Red fox | -51.58 | 83.7 | 75 | 60 |
| *Ictidomys tridecemlineatus* | Rodentia | Thirteen-lined ground squirrel | -51.54 | 84.5 | 85 | 80 |
| *Marmota flaviventris* | Rodentia | Yellow-bellied marmot | -51.50 | 83.8 | 85 | 80 |
| *Canis lupus familiaris* | Carnivora | Dog | -51.38 | 83.6 | 75 | 60 |
| *Canis lupus dingo* | Carnivora | Dingo | -51.11 | 84.1 | 75 | 60 |
| *Ceratotherium simum simum* | Perissodactyla | Southern white rhinoceros | -51.08 | 85.8 | 80 | 80 |
| *Ochotona princeps* | Lagomorpha | American pika | -51.01 | 80.8 | 80 | 80 |

| | | | | | | |
|---|---|---|---|---|---|---|
| *Rousettus aegyptiacus* | Chiroptera | Egyptian rousette | -50.91 | 78.9 | 75 | 80 |
| *Sus scrofa* | Artiodactyla | Pig | -50.74 | 81.4 | 75 | 80 |
| *Urocitellus parryii* | Rodentia | Arctic ground squirrel | -50.62 | 83.8 | 80 | 80 |
| *Lagenorhynchus obliquidens* | Artiodactyla | Pacific white-sided dolphin | -50.37 | 81.5 | 75 | 80 |
| *Nyctereutes procyonoides* | Carnivora | Raccoon dog | -50.20 | 84.1 | 70 | 40 |
| *Equus asinus* | Perissodactyla | Donkey | -50.02 | 86.0 | 70 | 60 |
| *Rhinolophus sinicus* | Chiroptera | Chinese rufous horseshoe bat | -49.91 | 80.7 | 70 | 60 |
| *Camelus bactrianus* | Artiodactyla | Bactrian camel | -49.88 | 83.2 | 75 | 60 |
| *Mirounga leonina* | Carnivora | Southern elephant seal | -49.75 | 82.6 | 60 | 60 |
| *Camelus dromedarius* | Artiodactyla | Arabian camel | -49.74 | 83.2 | 75 | 60 |
| *Phyllostomus discolor* | Chiroptera | Pale spear-nosed bat | -49.73 | 79.9 | 60 | 40 |
| *Camelus ferus* | Artiodactyla | Wild bactrian camel | -49.63 | 83.2 | 75 | 60 |
| *Orcinus orca* | Artiodactyla | Killer whale | -49.47 | 81.3 | 75 | 80 |
| *Pteropus vampyrus* | Chiroptera | Large flying fox | -49.29 | 80.7 | 70 | 80 |
| *Pteropus alecto* | Chiroptera | Black flying fox | -49.29 | 81.5 | 70 | 80 |
| *Globicephala melas* | Artiodactyla | Long-finned pilot whale | -49.19 | 81.3 | 75 | 80 |
| *Tursiops truncatus* | Artiodactyla | Atlantic bottle-nosed dolphin | -49.05 | 81.5 | 75 | 80 |
| *Lipotes vexillifer* | Artiodactyla | Yangtze river dolphin | -49.05 | 81.8 | 75 | 80 |
| *Dipodomys ordii* | Rodentia | Ords kangaroo rat | -49.01 | 82.2 | 80 | 60 |
| *Loxodonta africana* | Proboscidea | African savanna elephant | -48.80 | 81.3 | 75 | 60 |
| *Orycteropus afer* | Tubulidentata | Aardvark | -48.79 | 80.0 | 60 | 60 |
| *Enhydra lutris kenyoni* | Carnivora | Sea otter | -48.53 | 82.9 | 65 | 60 |
| *Trichechus manatus latirostris* | Sirenia | Florida manatee | -48.13 | 82.0 | 75 | 60 |
| *Paguma larvata* | Carnivora | Masked palm civet | -47.97 | 83.5 | 65 | 40 |
| *Octodon degus* | Rodentia | Common degu | -47.70 | 81.8 | 75 | 40 |
| *Vicugna pacos* | Artiodactyla | Alpaca | -47.11 | 83.4 | 75 | 60 |
| *Callithrix jacchus* | Primates | Common marmoset | -46.81 | 91.7 | 80 | 80 |
| *Saimiri boliviensis* | Primates | Black-capped squirrel monkey | -46.71 | 92.0 | 80 | 80 |
| *Otolemur garnettii* | Primates | Small-eared galago | -46.54 | 81.9 | 65 | 40 |
| *Eptesicus fuscus* | Chiroptera | Big brown bat | -46.46 | 80.5 | 55 | 60 |
| *Myotis brandtii* | Chiroptera | Brandts bat | -46.44 | 79.5 | 55 | 40 |
| *Aotus nancymaae* | Primates | Ma's night monkey | -46.28 | 92.2 | 80 | 80 |
| *Cyanistes caeruleus* | Passeriformes | Eurasian blue tit | -46.23 | 67.1 | 45 | 40 |
| *Sapajus apella* | Primates | Tufted capuchin | -46.13 | 92.5 | 80 | 80 |
| *Myotis davidii* | Chiroptera | David's myotis | -46.10 | 79.2 | 60 | 40 |
| *Chinchilla lanigera* | Rodentia | Long-tailed chinchilla | -46.05 | 84.7 | 75 | 40 |
| *Carlito syrichta* | Primates | Philippine tarsier | -45.68 | 84.1 | 70 | 60 |
| *Suricata suricatta* | Carnivora | Meerkat | -45.57 | 82.9 | 60 | 40 |
| *Condylura cristata* | Eulipotyphla | Star-nosed mole | -45.56 | 77.6 | 55 | 40 |
| *Microcebus murinus* | Primates | Gray mouse lemur | -45.25 | 77.9 | 70 | 60 |
| *Myotis lucifugus* | Chiroptera | Little brown bat | -45.12 | 79.4 | 55 | 40 |
| *Dasypus novemcinctus* | Cingulata | Nine-banded armadillo | -45.00 | 79.2 | 55 | 20 |
| *Echinops telfairi* | Afrosoricida | Small madagascar hedgehog | -44.99 | 75.3 | 50 | 20 |
| *Nothoprocta perdicaria* | Tinamiformes | Chilean tinamou | -44.98 | 66.9 | 50 | 40 |
| *Chrysochloris asiatica* | Afrosoricida | Cape golden mole | -44.75 | 79.0 | 50 | 40 |
| *Mus pahari* | Rodentia | Shrew mouse | -44.74 | 83.0 | 65 | 60 |
| *Rhinolophus ferrumequinum* | Chiroptera | Greater horseshoe bat | -44.47 | 81.5 | 60 | 40 |
| *Mastomys coucha* | Rodentia | Southern multimammate mouse | -44.42 | 81.8 | 65 | 60 |
| *Monodelphis domestica* | Didelphimorphia | Gray short-tailed opossum | -44.04 | 71.0 | 55 | 20 |
| *Serinus canaria* | Passeriformes | Island canary | -44.01 | 66.9 | 55 | 40 |
| *Grammomys surdaster* | Rodentia | African woodland thicket rat | -43.96 | 82.5 | 70 | 60 |
| *Hipposideros armiger* | Chiroptera | Great roundleaf bat | -43.96 | 80.5 | 65 | 80 |
| *Sturnus vulgaris* | Passeriformes | Common starling | -43.88 | 66.4 | 45 | 40 |
| *Tupaia chinensis* | Scandentia | Chinese tree shrew | -43.88 | 80.7 | 50 | 60 |
| *Calidris pugnax* | Charadriiformes | Ruff | -43.85 | 64.8 | 40 | 40 |
| *Gekko japonicus* | Squamata | Schlegel's japanese gecko | -43.77 | 63.5 | 45 | 20 |
| *Chrysemys picta* | Testudines | Painted turtle | -43.61 | 64.7 | 50 | 40 |
| *Desmodus rotundus* | Chiroptera | Vampire bat | -43.60 | 79.7 | 55 | 20 |
| *Rattus rattus* | Rodentia | Black rat | -43.34 | 80.7 | 60 | 60 |
| *Mus caroli* | Rodentia | Ryukyu mouse | -43.20 | 82.2 | 65 | 60 |
| *Geotrypetes seraphini* | Gymnophiona | Gaboon caecilian | -43.20 | 60.2 | 60 | 40 |
| *Struthio camelus australis* | Struthioniformes | South african ostrich | -43.17 | 65.2 | 50 | 40 |
| *Rattus norvegicus* | Rodentia | Brown rat | -43.14 | 82.5 | 60 | 60 |
| *Falco cherrug* | Falconiformes | Saker falcon | -43.02 | 64.7 | 40 | 20 |
| *Aquila chrysaetos chrysaetos* | Accipitriformes | Golden eagle | -42.97 | 65.9 | 45 | 20 |
| *Coturnix japonica* | Galliformes | Japanese quail | -42.92 | 67.0 | 45 | 40 |
| *Pantherophis guttatus* | Squamata | Corn snake | -42.90 | 58.8 | 50 | 60 |
| *Danio rerio* | Cypriniformes | Zebrafish | -42.82 | 57.7 | 55 | 60 |
| *Cygnus atratus* | Anseriformes | Black Swan | -42.81 | 65.6 | 50 | 40 |
| *Miniopterus natalensis* | Chiroptera | Natal long-fingered bat | -42.79 | 80.8 | 55 | 40 |
| *Rhinatrema bivittatum* | Gymnophiona | Two-lined caecilian | -42.72 | 61.6 | 50 | 40 |
| *Aptenodytes forsteri* | Sphenisciformes | Emperor penguin | -42.71 | 66.9 | 45 | 20 |
| *Mus musculus* | Rodentia | House mouse | -42.62 | 82.1 | 60 | 40 |
| *Larimichthys crocea* | Perciformes | Large yellow croaker | -42.60 | 58.6 | 50 | 40 |
| *Anas platyrhynchos* | Anseriformes | Duck | -42.54 | 65.3 | 50 | 40 |

| | | | | | | |
|---|---|---|---|---|---|---|
| *Phasianus colchicus* | Galliformes | Common pheasant | -42.44 | 66.2 | 50 | 40 |
| *Corvus moneduloides* | Passeriformes | New caledonian crow | -42.38 | 67.1 | 45 | 20 |
| *Opisthocomus hoazin* | Opisthocomiformes | Hoatzin | -42.37 | 63.0 | 50 | 40 |
| *Camarhynchus parvulus* | Passeriformes | Small tree finch | -42.34 | 67.2 | 55 | 40 |
| *Ictalurus punctatus* | Siluriformes | Channel catfish | -42.23 | 57.8 | 40 | 20 |
| *Chelonia mydas* | Testudines | Green sea turtle | -42.23 | 66.3 | 50 | 40 |
| *Oxyura jamaicensis* | Anseriformes | Ruddy duck | -42.20 | 65.7 | 45 | 20 |
| *Aythya fuligula* | Anseriformes | Tufted duck | -42.17 | 65.8 | 50 | 40 |
| *Fundulus heteroclitus* | Cyprinodontiformes | Mummichog | -42.17 | 58.3 | 40 | 0 |
| *Elephantulus edwardii* | Macroscelidea | Cape elephant shrew | -42.16 | 78.2 | 60 | 40 |
| *Ficedula albicollis* | Passeriformes | Collared flycatcher | -42.15 | 66.7 | 45 | 40 |
| *Pundamilia nyererei* | Cichliformes | | -42.11 | 58.3 | 55 | 40 |
| *Mesitornis unicolor* | Gruiformes | Brown roatelo | -42.09 | 65.4 | 50 | 20 |
| *Gallus gallus* | Galliformes | Chicken | -42.07 | 66.1 | 50 | 40 |
| *Lates calcarifer* | Perciformes | Barramundi perch | -42.06 | 59.5 | 50 | 20 |
| *Chelonoidis abingdonii* | Testudines | Pinta island tortoise | -42.06 | 66.7 | 45 | 40 |
| *Arvicanthis niloticus* | Rodentia | African grass rat | -42.06 | 81.1 | 65 | 60 |
| *Thamnophis elegans* | Squamata | Western terrestrial garter snake | -42.02 | 58.6 | 55 | 60 |
| *Oreochromis niloticus* | Cichliformes | Nile tilapia | -41.99 | 57.1 | 60 | 40 |
| *Melopsittacus undulatus* | Psittaciformes | Budgerigar | -41.98 | 66.4 | 50 | 40 |
| *Buceros rhinoceros silvestris* | Bucerotiformes | Rhinoceros hornbill | -41.97 | 65.7 | 45 | 40 |
| *Meleagris gallopavo* | Galliformes | Wild turkey | -41.94 | 56.3 | 50 | 40 |
| *Tyto alba* | Strigiformes | Western barn owl | -41.93 | 65.8 | 50 | 40 |
| *Betta splendens* | Anabantiformes | Siamese fighting fish | -41.87 | 58.1 | 45 | 40 |
| *Python bivittatus* | Squamata | Burmese python | -41.82 | 60.1 | 40 | 40 |
| *Ornithorhynchus anatinus* | Monotremata | Platypus | -41.82 | 68.6 | 50 | 40 |
| *Pangasianodon hypophthalmus* | Siluriformes | Striped catfish | -41.81 | 58.0 | 40 | 20 |
| *Pseudopodoces humilis* | Passeriformes | Tibetan ground jay | -41.78 | 67.7 | 40 | 20 |
| *Zootoca vivipara* | Squamata | Viviparous lizard | -41.77 | 65.0 | 50 | 40 |
| *Chaetura pelagica* | Apodiformes | Chimney swift | -41.76 | 66.1 | 50 | 40 |
| *Calypte anna* | Apodiformes | Annas hummingbird | -41.74 | 66.2 | 50 | 40 |
| *Pygocentrus nattereri* | Characiformes | Red-bellied piranha | -41.70 | 58.1 | 40 | 20 |
| *Poecilia reticulata* | Cyprinodontiformes | Guppy | -41.68 | 58.0 | 40 | 20 |
| *Nipponia nippon* | Pelecaniformes | Crested ibis | -41.66 | 66.1 | 45 | 20 |
| *Athene cunicularia* | Strigiformes | Burrowing owl | -41.64 | 63.7 | 50 | 40 |
| *Chlamydotis macqueenii* | Gruiformes | Macqueen's bustard | -41.63 | 65.6 | 45 | 20 |
| *Microcaecilia unicolor* | Gymnophiona | | -41.60 | 61.5 | 45 | 40 |
| *Pygoscelis adeliae* | Sphenisciformes | Adelie penguin | -41.58 | 66.0 | 45 | 20 |
| *Notothenia coriiceps* | Perciformes | Black rockcod | -41.48 | 58.3 | 50 | 40 |
| *Parus major* | Passeriformes | Great tit | -41.47 | 67.1 | 40 | 20 |
| *Mastacembelus armatus* | Synbranchiformes | Zig-zag eel | -41.38 | 57.9 | 40 | 20 |
| *Zonotrichia albicollis* | Passeriformes | White-throated sparrow | -41.31 | 66.0 | 50 | 40 |
| *Fulmarus glacialis* | Procellariiformes | Northern fulmar | -41.26 | 68.2 | 45 | 20 |
| *Gavia stellata* | Gaviiformes | Red-throated loon | -41.20 | 66.4 | 45 | 20 |
| *Astatotilapia calliptera* | Cichliformes | Eastern happy | -41.19 | 58.5 | 55 | 40 |
| *Numida meleagris* | Galliformes | Helmeted guineafowl | -41.04 | 65.3 | 45 | 40 |
| *Taeniopygia guttata* | Passeriformes | Zebra finch | -40.97 | 66.2 | 45 | 20 |
| *Sorex araneus* | Eulipotyphla | Eurasian common shrew | -40.97 | 73.8 | 45 | 40 |
| *Sinocyclocheilus rhinocerous* | Cypriniformes | | -40.96 | 57.7 | 50 | 40 |
| *Poecilia mexicana* | Cyprinodontiformes | Cave molly | -40.85 | 57.9 | 40 | 20 |
| *Pseudonaja textilis* | Squamata | Eastern brown snake | -40.81 | 58.2 | 40 | 40 |
| *Salarias fasciatus* | Blenniiformes | Jewelled blenny | -40.72 | 58.3 | 45 | 20 |
| *Pelodiscus sinensis* | Testudines | Chinese soft-shelled turtle | -40.68 | 67.0 | 50 | 40 |
| *Perca flavescens* | Perciformes | Yellow perch | -40.53 | 58.4 | 45 | 40 |
| *Merops nubicus* | Coraciiformes | Northern carmine bee-eater | -40.52 | 66.3 | 45 | 20 |
| *Seriola dumerili* | Carangiformes | Greater amberjack | -40.45 | 58.5 | 35 | 20 |
| *Thamnophis sirtalis* | Squamata | Common garter snake | -40.43 | 57.3 | 55 | 60 |
| *Phaethon lepturus* | Phaethontiformes | White-tailed tropicbird | -40.37 | 65.4 | 50 | 40 |
| *Parambassis ranga* | Ovalentaria incertae sedis | Indian glassy fish | -40.36 | 57.8 | 40 | 20 |
| *Neolamprologus brichardi* | Cichliformes | Princess cichlid | -40.35 | 58.2 | 55 | 40 |
| *Vombatus ursinus* | Diprotodontia | Common wombat | -40.33 | 71.9 | 55 | 40 |
| *Trachemys scripta elegans* | Testudines | Red-eared slider | -40.31 | 65.1 | 50 | 40 |
| *Apteryx rowi* | Apterygiformes | Okarito brown kiwi | -40.29 | 66.2 | 45 | 40 |
| *Poecilia formosa* | Cyprinodontiformes | Amazon molly | -40.29 | 57.8 | 40 | 20 |
| *Poecilia latipinna* | Cyprinodontiformes | Sailfin molly | -40.29 | 57.7 | 40 | 20 |
| *Carassius auratus* | Cypriniformes | Goldfish | -40.28 | 57.9 | 50 | 40 |
| *Stegastes partitus* | Ovalentaria incertae sedis | Bicolor damselfish | -40.24 | 58.5 | 45 | 20 |
| *Denticeps clupeoides* | Clupeiformes | Denticle herring | -40.20 | 59.8 | 45 | 40 |
| *Gopherus evgoodei* | Testudines | Goodes thornscrub tortoise | -40.20 | 66.2 | 45 | 40 |
| *Protobothrops mucrosquamatus* | Squamata | Protobothrops mucrosquamatus | -40.18 | 55.2 | 50 | 60 |
| *Seriola lalandi dorsalis* | Carangiformes | | -40.13 | 58.5 | 35 | 20 |
| *Xenopus tropicalis* | Anura | Tropical clawed frog | -39.96 | 55.8 | 45 | 20 |
| *Charadrius vociferus* | Charadriiformes | Killdeer | -39.90 | 66.6 | 50 | 20 |
| *Anolis carolinensis* | Squamata | Green anole | -39.89 | 63.0 | 40 | 20 |
| *Lonchura striata domestica* | Passeriformes | Bengalese finch | -39.84 | 65.6 | 45 | 20 |

| | | | | | | |
|---|---|---|---|---|---|---|
| *Corapipo altera* | Passeriformes | White-ruffed manakin | -39.83 | 65.2 | 40 | 0 |
| *Esox lucius* | Esociformes | Northern pike | -39.71 | 56.1 | 40 | 40 |
| *Maylandia zebra* | Cichliformes | Zebra mbuna | -39.66 | 58.5 | 55 | 40 |
| *Neopelma chrysocephalum* | Passeriformes | Saffron-crested tyrant-manakin | -39.51 | 65.5 | 50 | 40 |
| *Pogona vitticeps* | Squamata | Central bearded dragon | -39.51 | 63.3 | 45 | 20 |
| *Catharus ustulatus* | Passeriformes | Swainsons thrush | -39.50 | 61.9 | 35 | 0 |
| *Haliaeetus albicilla* | Accipitriformes | White-tailed eagle | -39.46 | 70.3 | 45 | 20 |
| *Corvus brachyrhynchos* | Passeriformes | American crow | -39.44 | 67.8 | 45 | 20 |
| *Haplochromis burtoni* | Cichliformes | Burtons mouthbrooder | -39.43 | 58.5 | 55 | 40 |
| *Manacus vitellinus* | Passeriformes | Golden-collared manakin | -39.37 | 65.5 | 40 | 0 |
| *Cynoglossus semilaevis* | Pleuronectiformes | Tongue sole | -39.37 | 57.9 | 45 | 40 |
| *Empidonax traillii* | Passeriformes | Willow flycatcher | -39.36 | 65.9 | 50 | 40 |
| *Archocentrus centrarchus* | Cichliformes | Flier cichlid | -39.35 | 58.8 | 55 | 40 |
| *Amphiprion ocellaris* | Ovalentaria incertae sedis | Clown anemonefish | -39.32 | 58.0 | 45 | 20 |
| *Hippocampus comes* | Syngnathiformes | Tiger tail seahorse | -39.32 | 57.3 | 45 | 40 |
| *Myripristis murdjan* | Holocentriformes | Pinecone soldierfish | -39.17 | 58.9 | 45 | 20 |
| *Corvus cornix cornix* | Passeriformes | Hooded crow | -39.16 | 66.8 | 40 | 0 |
| *Amblyraja radiata* | Rajiformes | Thorny skate | -38.93 | 59.2 | 35 | 0 |
| *Lepidothrix coronata* | Passeriformes | Blue-crowned manakin | -38.83 | 65.6 | 40 | 0 |
| *Alligator mississippiensis* | Crocodilia | American alligator | -38.81 | 66.7 | 45 | 0 |
| *Alligator sinensis* | Crocodilia | Chinese alligator | -38.79 | 66.4 | 45 | 0 |
| *Nothobranchius furzeri* | Cyprinodontiformes | Turquoise killifish | -38.73 | 59.5 | 45 | 0 |
| *Sander lucioperca* | Perciformes | Pike-perch | -38.73 | 59.3 | 45 | 40 |
| *Crocodylus porosus* | Crocodilia | Saltwater crocodile | -38.71 | 66.4 | 40 | 0 |
| *Erinaceus europaeus* | Eulipotyphla | Western european hedgehog | -38.67 | 79.1 | 50 | 0 |
| *Pipra filicauda* | Passeriformes | Wire-tailed manakin | -38.54 | 65.5 | 40 | 0 |
| *Sinocyclocheilus anshuiensis* | Cypriniformes | | -38.49 | 57.2 | 50 | 40 |
| *Gavialis gangeticus* | Crocodilia | Gharial | -38.48 | 66.5 | 45 | 0 |
| *Xiphophorus hellerii* | Cyprinodontiformes | Green swordtail | -38.46 | 58.4 | 35 | 0 |
| *Strigops habroptila* | Psittaciformes | Kakapo | -38.32 | 64.6 | 45 | 20 |
| *Acanthochromis polyacanthus* | Ovalentaria incertae sedis | Spiny chromis | -38.30 | 55.8 | 35 | 0 |
| *Nanorana parkeri* | Anura | High himalaya frog | -38.23 | 61.2 | 35 | 0 |
| *Erpetoichthys calabaricus* | Polypteriformes | Rope fish | -38.22 | 61.5 | 45 | 20 |
| *Oreochromis aureus* | Cichliformes | Israeli tilapia | -38.15 | 57.1 | 55 | 40 |
| *Kryptolebias marmoratus* | Cyprinodontiformes | Mangrove rivulus | -38.13 | 58.6 | 45 | 20 |
| *Etheostoma spectabile* | Perciformes | Orangethroat darter | -37.98 | 58.9 | 40 | 20 |
| *Anarrhichthys ocellatus* | Perciformes | Wolf eel | -37.83 | 58.8 | 45 | 20 |
| *Callorhinchus milii* | Chimaeriformes | Ghost shark | -37.70 | 58.1 | 35 | 20 |
| *Cyprinodon variegatus* | Cyprinodontiformes | Sheepshead minnow | -37.67 | 58.0 | 45 | 20 |
| *Electrophorus electricus* | Gymnotiformes | Electric eel | -37.36 | 58.5 | 40 | 20 |
| *Chanos chanos* | Gonorynchiformes | Milkfish | -37.27 | 57.7 | 40 | 20 |
| *Chiroxiphia lanceolata* | Passeriformes | Lance-tailed manakin | -37.18 | 65.4 | 40 | 0 |
| *Anabas testudineus* | Anabantiformes | Climbing perch | -36.81 | 59.4 | 40 | 20 |
| *Scleropages formosus* | Osteoglossiformes | Asian bonytongue | -36.79 | 59.1 | 35 | 0 |
| *Latimeria chalumnae* | Coelacanthiformes | Coelacanth | -36.68 | 58.0 | 40 | 0 |
| *Oncorhynchus mykiss* | Salmoniformes | Rainbow trout | -36.65 | 58.2 | 35 | 0 |
| *Oryzias melastigma* | Beloniformes | Marine medaka | -36.57 | 56.9 | 45 | 0 |
| *Eurypyga helias* | Gruiformes | Sunbittern | -36.54 | 65.1 | 35 | 0 |
| *Boleophthalmus pectinirostris* | Gobiiformes | Great blue-spotted mudskipper | -36.34 | 56.6 | 30 | 20 |
| *Gouania willdenowi* | Gobiesociformes | Blunt-snouted clingfish | -36.31 | 57.6 | 35 | 0 |
| *Haliaeetus leucocephalus* | Accipitriformes | Bald eagle | -36.19 | 68.6 | 40 | 20 |
| *Tachysurus fulvidraco* | Siluriformes | Yellow catfish | -36.07 | 57.4 | 40 | 20 |
| *Labrus bergylta* | Labriformes | Ballan wrasse | -35.94 | 57.6 | 40 | 20 |
| *Cottoperca gobio* | Perciformes | Frogmouth | -35.92 | 57.1 | 45 | 20 |
| *Takifugu rubripes* | Tetraodontiformes | Torafugu | -35.88 | 56.8 | 35 | 0 |
| *Xiphophorus couchianus* | Cyprinodontiformes | Monterrey platyfish | -35.79 | 58.5 | 35 | 0 |
| *Astyanax mexicanus* | Characiformes | Mexican tetra | -35.53 | 60.9 | 40 | 20 |
| *Lepisosteus oculatus* | Lepisosteiformes | Spotted gar | -35.26 | 59.5 | 35 | 0 |
| *Echeneis naucrates* | Carangiformes | Live sharksucker | -34.66 | 57.7 | 40 | 20 |
| *Paralichthys olivaceus* | Pleuronectiformes | Bastard halibut | -34.59 | 57.1 | 40 | 20 |
| *Xiphophorus maculatus* | Cyprinodontiformes | Southern platyfish | -34.09 | 58.2 | 35 | 0 |
| *Clupea harengus* | Clupeiformes | Atlantic herring | -33.62 | 56.0 | 30 | 0 |
| *Gadus morhua* | Gadiformes | Atlantic cod | -33.30 | 57.3 | 45 | 40 |

**Table S5**. Comparison of experimental and predicted N-glycosylation sites on hACE2.

| Position | Experimental | NGlycPred | N-GlyDE | NetNGlyc |
|---|---|---|---|---|
| 53 | Yes | No | Yes | Yes |
| 90 | Yes | Yes | Yes | Yes |
| 103 | Yes | Yes | Yes | Yes |
| 322 | Yes | Yes | Yes | Yes |
| 432 | Yes | No | Yes | Yes |
| 546 | Yes | Yes | No | No |
| 690 | Yes | Yes | Yes | Yes |

**Table S6**. Potential N-glycosylation sites in the 285 selected ACE2 proteins. Each site is shown in the format of digits-NX(S/T), where the number is the position of the glycosylated asparagine (indexed from 1), and X can be any canonical amino-acid type except proline. Interface glycosylation sites are highlighted in bold.

| Accession ID | Class | N-glycosylation sites [denoted as index-NX(S/T)] |
|---|---|---|
| AAX63775.1 | Mammalia | 53-NIT, 216-NYS, 322-NMT, 432-NET, 546-NST, 580-NVT, 660-NQT, 690-NVS |
| ABW16956.1 | Mammalia | 52-NIT, 215-NYS, 298-NQS, 321-NMT, 545-NSS, 659-NQT, 689-NVS |
| AGZ48803.1 | Mammalia | 82-NYS, 90-NVT, 322-NMT, 329-NNS, 432-NET, 546-NST, 690-NLS |
| NP_001012006.1 | Mammalia | 53-NIT, 82-NFS, 90-NAT, 299-NQS, 432-NET, 546-NST, 601-NST, 660-NQT, 690-NVS, 772-NET |
| NP_001116542.1 | Mammalia | 53-NIT, 299-NQS, 322-NMT, 329-NNS, 546-NST, 601-NSS, 660-NET, 690-NMS |
| NP_001124604.1 | Mammalia | 53-NIT, 90-NLT, 103-NGS, 322-NMT, 432-NET, 546-NST, 690-NVS |
| NP_001129168.1 | Mammalia | 53-NIT, 90-NLT, 103-NGS, 322-NMT, 432-NET, 546-NST, 690-NVS |
| NP_001158732.1 | Mammalia | 52-NIT, 215-NYS, 298-NQS, 321-NMT, 545-NSS, 659-NQT, 681-NFS, 689-NVS |
| NP_001277036.1 | Mammalia | 53-NIT, 90-NLT, 298-NQS, 328-NNS, 431-NET, 545-NST, 659-NET, 689-NVS, 790-NNS |
| NP_001297119.1 | Mammalia | 53-NIT, 299-NQS, 322-NMT, 546-NSS, 660-NQT, 690-NMS |
| NP_001358344.1 | Mammalia | 53-NIT, 90-NLT, 103-NGS, 322-NMT, 432-NET, 546-NST, 690-NVS |
| NP_081562.2 | Mammalia | 53-NIT, 536-NGS, 546-NST, 660-NQT, 690-NVS, 772-NET |
| XP_001490241.1 | Mammalia | 53-NIT, 90-NLT, 322-NMT, 546-NST, 609-NWS, 660-NQT, 690-NAS |
| XP_001515597.2 | Mammalia | 52-NIS, 78-NAS, 83-NLS, 321-NMT, 328-NNS, 678-NET, 790-NGS |
| XP_002194303.4 | Aves | **39-NIS**, 54-NIT, 80-NAS, 324-NMT, 536-NHT, 548-NST, 600-NNS, 692-NIS |
| XP_002719891.1 | Mammalia | 53-NIT, 90-NLT, 134-NQS, 432-NET, 546-NST, 636-NDS, 660-NQT, 791-NNS |
| XP_002938293.2 | Amphibia | 54-NIT, 76-NAS, 331-NNS, 859-NTT |
| XP_003261132.2 | Mammalia | 53-NIT, 90-NLT, 103-NGS, 322-NMT, 432-NET, 546-NST, 690-NVS |
| XP_003445853.2 | Actinopterygii | 390-NLS, 470-NIT, 681-NKT, 710-NAT |
| XP_003503283.1 | Mammalia | 53-NIT, 82-NYS, 432-NET, 546-NST, 658-NKT, 690-NVS |
| XP_003791912.1 | Mammalia | 53-NIT, 90-NRT, 218-NRS, 432-NET, 609-NWS, 690-NVS |
| XP_004269705.1 | Mammalia | 53-NIT, 90-NLT, 298-NQS, 321-NMT, 545-NST, 614-NQS, 659-NKT, 689-NMS |
| XP_004386381.1 | Mammalia | 53-NIT, 154-NST, 252-NQT, 298-NAT, 541-NST, 575-NVT, 596-NSS, 655-NQT, 685-NVS |
| XP_004415448.1 | Mammalia | 53-NIT, 216-NYS, 322-NMT, 432-NET, 616-NTS, 711-NLS |
| XP_004435206.1 | Mammalia | 53-NIT, 90-NVT, 216-NYS, 299-NQT, 322-NMT, 546-NST, 580-NVT, 660-NQT, 690-NVS |
| XP_004449124.1 | Mammalia | 52-NIT, 81-NFS, 89-NLT, 217-NRS, 321-NMT, 328-NNS, 545-NST, 600-NSS, 608-NWS, 659-NET |
| XP_004543482.1 | Actinopterygii | 54-NIT, 390-NLS, 470-NIT, 662-NKT |
| XP_004597549.2 | Mammalia | 3-NMS, 56-NIT, 93-NLT, 435-NET, 549-NST, 639-NDS, 693-NVS |
| XP_004612266.1 | Mammalia | **23-NAT**, **41-NSS**, 52-NIT, 121-NMS, 135-NNT, 320-NMT, 532-NHS, 599-NSS, 634-NDS, 658-NET, 688-NAS |
| XP_004671523.1 | Mammalia | 53-NIT, 432-NET, 546-NST, 690-NVS, 772-NQT |
| XP_004710002.1 | Mammalia | **38-NVS**, 53-NIT, 489-NES, 541-NST, 655-NQT |
| XP_004866157.1 | Mammalia | 53-NIT, 90-NLT, 546-NST, 690-NIT, 772-NRT |
| XP_005037422.1 | Aves | **21-NVT**, 54-NIT, 80-NAS, 324-NMT, 331-NNS, 536-NHT, 548-NST, 582-NAT, 600-NNS, 638-NDS, 692-NVT |
| XP_005074266.1 | Mammalia | 53-NIT, 82-NYS, 90-NLT, 432-NET, 546-NST, 658-NKT, 690-NVS, 772-NET |
| XP_005151516.2 | Aves | 52-NIT, 78-NAS, 153-NST, 329-NNS, 534-NHT, 580-NAS, 690-NIS |
| XP_005169416.1 | Actinopterygii | 328-NNS, 545-NST, 677-NFT, 687-NET |
| XP_005228485.1 | Mammalia | 53-NIT, 90-NLT, 298-NQS, 431-NET, 545-NST, 659-NET, 689-NVS |
| XP_005316051.3 | Mammalia | 63-NIT, 100-NFT, 164-NST, 332-NMT, 556-NST, 619-NWS, 670-NQT, 700-NVS, 782-NQT |
| XP_005358818.1 | Mammalia | 53-NIT, 546-NST, 659-NQT, 689-NVS, 771-NET |
| XP_005443093.2 | Aves | **41-NIS**, 56-NIT, 82-NAS, 333-NNS, 538-NHT, 550-NSS, 576-NVT, 602-NNS, 694-NIS |
| XP_005491832.2 | Aves | 54-NIT, 80-NAS, 331-NNS, 548-NST, 574-NIT, 600-NNS, 692-NIS |
| XP_005516712.1 | Aves | **19-NVT**, **37-NIS**, 52-NIT, 78-NAS, 546-NST, 598-NNS |
| XP_005593094.1 | Mammalia | 53-NIT, 90-NLT, 103-NGS, 322-NMT, 432-NET, 546-NST, 690-NVS |
| XP_005724169.1 | Actinopterygii | 54-NIT, 390-NLS, 470-NIT, 662-NKT |
| XP_005799835.1 | Actinopterygii | **37-NAS**, 391-NLS, 471-NIS |
| XP_005903173.1 | Mammalia | 53-NIT, 90-NLT, 298-NQS, 431-NET, 545-NST, 659-NET, 689-NVS, 790-NNS |
| XP_005943362.1 | Actinopterygii | 54-NIT, 390-NLS, 470-NIT, 662-NKT |
| XP_005997915.2 | Sarcopterygii | 97-NIT, 123-NAS, 375-NYS, 580-NHT, 592-NST, 618-NVT, 739-NIS, 742-NSS |
| XP_006041602.1 | Mammalia | 52-NIT, 89-NLT, 297-NQS, 430-NET, 544-NST, 658-NET, 688-NVS, 789-NNS |
| XP_006122891.1 | Reptilia | 52-NIT, 78-NAS, 89-NHT, 303-NAT, 387-NLS, 432-NET, 580-NAT, 598-NNS, 660-NQT, 690-NAT |
| XP_006164754.1 | Mammalia | 53-NIT, 78-NQS, 218-NRT, 322-NMT, 546-NST, 690-NVS |
| XP_006194263.1 | Mammalia | 53-NIT, 90-NVT, 299-NQS, 322-NMT, 546-NST, 660-NQT, 690-NVS, 791-NNS |
| XP_006212709.1 | Mammalia | 53-NIT, 90-NVT, 299-NQS, 322-NMT, 546-NST, 690-NVS, 791-NNS |
| XP_006639185.1 | Actinopterygii | **34-NAT**, 52-NIT, 328-NES, 387-NLS, 432-NET, 546-NST, 691-NNS, 692-NST, 790-NQT, 805-NHT |
| XP_006775273.1 | Mammalia | 53-NIT, 103-NGS, 431-NET, 493-NET, 674-NQT, 704-NVS, 799-NLS |
| XP_006780474.1 | Actinopterygii | 54-NIT, 390-NLS, 470-NIT, 662-NKT |
| XP_006835673.1 | Mammalia | 52-NIT, 89-NST, 153-NST, 316-NMT, 540-NST, 654-NQT, 684-NQS |
| XP_006892457.1 | Mammalia | 53-NIT, 154-NST, 298-NAT, 317-NMT, 427-NET, 541-NST, 575-NVT, 610-NQS, 654-NQT, 683-NGS |
| XP_006911709.1 | Mammalia | 53-NIT, 213-NGS, 322-NMT, 432-NET, 546-NST, 660-NLT, 690-NVS |
| XP_006973269.1 | Mammalia | 53-NIT, 82-NYS, 546-NST, 658-NKT, 690-NVS, 772-NET |
| XP_007070561.1 | Reptilia | 52-NIT, 78-NAS, 329-NNS, 387-NLS, 435-NET, 537-NHT, 549-NST, 583-NAT, 601-NNS, 663-NQT, 677-NVT, 693-NNT |
| XP_007090142.1 | Mammalia | 45-NIT, 82-NTT, 208-NYS, 291-NQS, 314-NMT, 538-NSS, 572-NVT, 652-NQT, 682-NVS |
| XP_007431942.2 | Reptilia | 77-NIT, 103-NAS, 344-NMT, 620-NNS, 631-NWT, 682-NQT, 712-NVS |
| XP_007466389.1 | Mammalia | 53-NIT, 90-NLT, 298-NQS, 321-NMT, 545-NST, 659-NET, 689-NVS, 790-NNS |
| XP_007500935.1 | Mammalia | 53-NIT, 75-NQS, 90-NAT, 301-NWS, 322-NMT, 432-NET, 546-NST, 689-NGT |
| XP_007538670.1 | Mammalia | **38-NVS**, 53-NIT, 103-NGS, 154-NST, 216-NYS, 329-NNS, 432-NET, 580-NVT, 660-NQT, 674-NLT |
| XP_007560208.1 | Actinopterygii | **37-NAS**, 391-NLS, 471-NIS |
| XP_007889845.1 | Chondrichthyes | 53-NIT, 548-NST, 702-NST, 719-NTT, 832-NNT |
| XP_007951028.1 | Mammalia | **38-NLS**, 53-NIT, 90-NST, 153-NST, 297-NAT, 488-NES, 540-NST, 601-NTS, 684-NES |
| XP_007989304.1 | Mammalia | 53-NIT, 90-NLT, 103-NGS, 322-NMT, 432-NET, 546-NST, 690-NVS |

| | | |
|---|---|---|
| XP_008062810.1 | Mammalia | 53-NIT, 90-NST, 103-NGS, 218-NST, 322-NMT, 432-NET |
| XP_008105455.1 | Reptilia | **46-NRS**, 61-NIT, 98-NDT, 555-NST, 589-NAS, 607-NNT, 683-NVT, 698-NDS, 809-NNS, 810-NST |
| XP_008153150.1 | Mammalia | **24-NAT**, 53-NIT, 90-NLT, 103-NGS, 215-NYS, 279-NLT, 328-NNS, 431-NET, 545-NST, 666-NQT, 696-NMS, 791-NLS |
| XP_008290762.1 | Actinopterygii | 54-NIT, 390-NLS, 470-NIS, 549-NST, 664-NQT |
| XP_008402714.1 | Actinopterygii | **37-NAS**, 391-NLS, 471-NIS |
| XP_008492997.2 | Aves | 52-NIT, 78-NAS, 329-NNS, 534-NHT, 546-NST, 598-NNS, 690-NIS |
| XP_008542995.1 | Mammalia | 53-NIT, 90-NLT, 322-NMT, 546-NST, 609-NWS, 660-NQT, 690-NAS |
| XP_008694637.1 | Mammalia | 38-NIT, 75-NST, 201-NYS, 307-NMT, 417-NET, 531-NSS, 675-NVS |
| XP_008839098.1 | Mammalia | 52-NIT, 431-NET, 545-NST, 689-NVS, 771-NQT |
| XP_008937519.1 | Aves | 52-NIT, 82-NAT, 146-NST, 524-NHT, 588-NNS, 682-NDT |
| XP_008972428.2 | Mammalia | 53-NIT, 90-NLT, 103-NGS, 322-NMT, 432-NET, 546-NST, 690-NVS |
| XP_008987241.1 | Mammalia | 53-NIT, 90-NLT, 103-NGS, 322-NMT, 609-NWS, 690-NVS |
| XP_009087922.1 | Aves | 52-NIT, 78-NAS, 322-NMT, 329-NNS, 534-NHT, 546-NST, 580-NAS, 598-NNS, 690-NIS |
| XP_009275140.1 | Aves | **37-NIS**, 52-NIT, 78-NAS, 257-NST, 329-NNS, 534-NHT, 572-NVT, 598-NNS, 636-NES, 690-NIS |
| XP_009323767.1 | Aves | **37-NIS**, 52-NIT, 78-NAS, 257-NST, 329-NNS, 523-NHT, 569-NAT, 587-NNS, 679-NIS |
| XP_009474590.1 | Aves | **37-NIS**, 52-NIT, 78-NAS, 153-NST, 329-NNS, 534-NHS, 598-NNS |
| XP_009574896.1 | Aves | 52-NIT, 78-NAS, 153-NST, 329-NNS, 534-NHT, 546-NST, 598-NNS |
| XP_009667495.1 | Aves | 52-NIT, 78-NAS, 322-NMT, 329-NNS, 467-NIT, 546-NST, 580-NAT, 598-NNS, 690-NIS, 782-NFS |
| XP_009816127.1 | Aves | **37-NIS**, 52-NIT, 78-NAS, 153-NST, 329-NNS, 534-NHT, 572-NVT, 598-NNS, 690-NIS, 775-NRS |
| XP_009887331.1 | Aves | **37-NIS**, 52-NIT, 78-NAS, 280-NLT, 329-NNS, 534-NHT, 546-NST, 598-NNS, 674-NQT, 690-NVS |
| XP_009925641.1 | Aves | **37-NIS**, 52-NIT, 150-NST, 191-NYS, 326-NNS, 543-NSS, 577-NAT, 595-NNS |
| XP_009938970.1 | Aves | 52-NIT, 78-NAS, 153-NST, 194-NYS, 329-NNS, 549-NHT, 561-NST, 613-NNS, 705-NNS |
| XP_009992128.1 | Aves | **19-NVT**, 52-NIT, 78-NAS, 194-NYS, 534-NHT, 598-NNS, 690-NIS |
| XP_010120523.1 | Aves | **37-NIS**, 52-NIT, 78-NAS, 153-NST, 194-NYS, 329-NNS, 533-NHT, 597-NNS, 635-NDS, 689-NIS |
| XP_010136813.1 | Aves | 63-NAS, 307-NMT, 314-NNS, 519-NHT, 557-NVT, 583-NNS, 675-NIS |
| XP_010156467.1 | Aves | **37-NIS**, 52-NIT, 78-NAS, 194-NYS, 534-NHT, 546-NST, 598-NNS |
| XP_010178703.1 | Aves | 52-NIT, 78-NAS, 329-NNS, 546-NST, 580-NAT, 598-NNS, 690-NMS |
| XP_010290019.1 | Aves | 52-NIT, 78-NAS, 153-NST, 329-NNS, 534-NHT, 598-NNS, 690-NVS |
| XP_010334925.1 | Mammalia | 53-NIT, 90-NLT, 103-NGS, 322-NMT, 609-NWS, 660-NQT, 690-NVS |
| XP_010364367.2 | Mammalia | 53-NIT, 90-NLT, 103-NGS, 136-NNS, 322-NMT, 432-NET, 546-NST, 690-NVS |
| XP_010392735.2 | Aves | **39-NIS**, 54-NIT, 80-NAS, 324-NMT, 536-NHT, 548-NST, 600-NNS, 692-NIS |
| XP_010579828.1 | Aves | **12-NIS**, 27-NIT, 53-NAS, 128-NST, 169-NYS, 304-NNS, 521-NSS, 555-NAT, 573-NNS, 665-NIS |
| XP_010643477.1 | Mammalia | 53-NIT, 90-NLT, 432-NET, 546-NST, 660-NKT, 690-NIT, 772-NKT, 791-NNT |
| XP_010730146.1 | Actinopterygii | 54-NIT, 76-NMS, 218-NYT, 331-NNS, 389-NLS, 434-NET, 577-NDT, 605-NRT, 678-NDT |
| XP_010790455.1 | Actinopterygii | 76-NMS, 331-NKS, 390-NLS, 430-NFT, 435-NET, 470-NIT, 664-NQT |
| XP_010833001.1 | Mammalia | 53-NIT, 90-NLT, 298-NQS |
| XP_010884777.2 | Actinopterygii | 69-NIT, 405-NQS, 450-NET, 564-NST, 679-NKT, 682-NVS, 806-NAT |
| XP_010966303.1 | Mammalia | 53-NIT, 90-NVT, 299-NQS, 322-NMT, 546-NST, 660-NQT, 690-NVS, 791-NNS |
| XP_010991717.1 | Mammalia | 53-NIT, 90-NVT, 299-NQS, 322-NMT, 546-NST, 660-NQT, 690-NVS, 791-NNS |
| XP_011361275.1 | Mammalia | 53-NIT, 213-NGS, 321-NMT, 431-NET, 545-NST, 659-NLT, 689-NVS |
| XP_011733505.1 | Mammalia | 53-NIT, 90-NLT, 103-NGS, 322-NMT, 432-NET, 546-NST, 690-NVS |
| XP_011850923.1 | Mammalia | 53-NIT, 90-NLT, 103-NGS, 322-NMT, 432-NET, 546-NST, 690-NVS |
| XP_011891198.1 | Mammalia | 53-NIT, 90-NLT, 103-NGS, 322-NMT, 432-NET, 546-NST, 690-NVS |
| XP_011961657.1 | Mammalia | 53-NIT, 90-NLT, 298-NQS, 431-NET, 545-NST, 659-NET, 689-NVS, 790-NNS |
| XP_012290105.1 | Mammalia | 53-NIT, 90-NLT, 103-NGS, 322-NMT, 660-NQT, 690-NVS |
| XP_012494185.1 | Mammalia | 74-NIT, 111-NVT, 239-NRS, 343-NMT, 453-NET, 567-NST, 630-NWS, 681-NQT, 711-NVS |
| XP_012585871.1 | Mammalia | **23-NQT**, **42-NSS**, 53-NIT, 317-NMT, 655-NET |
| XP_012887572.1 | Mammalia | 53-NIT, 432-NET, 660-NQT, 674-NLT, 689-NNS, 690-NSS |
| XP_012949915.2 | Aves | 52-NIT, 78-NAS, 322-NMT, 534-NHT, 546-NST, 580-NAT, 598-NNS, 634-NKS, 687-NAS |
| XP_013362428.1 | Mammalia | 53-NIT, 90-NLT, 299-NQS, 546-NST, 601-NAS, 660-NQT, 690-NIS |
| XP_013926936.1 | Reptilia | 77-NLT, 103-NAS, 114-NET, 130-NSS, 176-NWS, 309-NKT, 351-NNS |
| XP_014399780.1 | Mammalia | 53-NIT, 90-NLT, 103-NGS, 215-NYS, 279-NLT, 328-NNS, 431-NET, 674-NQT, 704-NIS, 799-NLS |
| XP_014713133.1 | Mammalia | 53-NIT, 90-NLT, 300-NMT, 524-NST, 587-NWS, 638-NQT, 668-NAS |
| XP_014731370.1 | Aves | 54-NIT, 80-NAS, 324-NMT, 548-NST, 574-NVT, 600-NNS, 692-NIT |
| XP_014815705.1 | Aves | 52-NIT, 78-NAS, 153-NST, 329-NNS, 546-NST, 580-NAT, 598-NNS, 690-NIS |
| XP_014837025.1 | Actinopterygii | **37-NAS**, 391-NLS, 471-NIS |
| XP_014895313.1 | Actinopterygii | **37-NAS**, 391-NLS, 471-NIS |
| XP_015226730.1 | Actinopterygii | **38-NAT**, 56-NIT, 284-NMS, 333-NNS, 392-NLS, 472-NFS, 800-NRS |
| XP_015273067.1 | Reptilia | **37-NLS**, 52-NIT, 102-NGS, 194-NYS, 303-NVT, 322-NMT, 534-NHT, 546-NST, 572-NVT, 580-NAT, 615-NDT, 660-NET, 690-NDS |
| XP_015343540.1 | Mammalia | 63-NIT, 100-NFT, 164-NST, 332-NMT, 442-NET, 556-NST, 619-NWS, 670-NQT, 700-NVS, 782-NQT |
| XP_015486815.1 | Aves | **24-NVT**, **42-NIS**, 57-NIT, 83-NAS, 140-NNS, 141-NSS, 551-NST, 585-NAT, 603-NNS |
| XP_015742063.1 | Aves | 52-NIT, 78-NAS, 194-NYS, 280-NLT, 322-NMT, 534-NHT, 546-NST, 580-NAT, 598-NNS, 690-NVS |
| XP_015808977.1 | Actinopterygii | **36-NAT**, 158-NYS, 390-NLS, 435-NET, 664-NQT, 678-NET |
| XP_015974412.1 | Mammalia | 136-NNS, 213-NGS, 322-NMT, 432-NET, 546-NST, 660-NLT, 690-NVS |
| XP_016058453.1 | Mammalia | 53-NIT, 68-NWS, 90-NSS, 216-NYS, 280-NVT, 322-NMT, 329-NNS, 546-NST, 659-NQT, 689-NVS, 784-NLS |
| XP_016345325.1 | Actinopterygii | 318-NMS, 545-NST, 681-NET |
| XP_016422243.1 | Actinopterygii | 318-NMS, 545-NST, 681-NET |
| XP_016798468.1 | Mammalia | 53-NIT, 90-NLT, 103-NGS, 322-NMT, 432-NET, 546-NST, 690-NVS |
| XP_016887914.1 | Actinopterygii | **41-NYS**, 52-NIT, 329-NNS, 388-NLS, 535-NHT, 662-NQT, 676-NET |
| XP_017295385.1 | Actinopterygii | 4-NMS, **41-NAT**, 163-NYS, 336-NNS, 395-NLS, 671-NQT |
| XP_017313836.1 | Actinopterygii | 318-NMS, 546-NST, 692-NAT, 789-NGT |
| XP_017505746.1 | Mammalia | 53-NIT, 90-NDT, 216-NYS, 299-NQT, 432-NET, 636-NDS, 690-NVS |
| XP_017550079.1 | Actinopterygii | 52-NIT, 318-NMS, 328-NNS, 545-NST, 691-NAS |
| XP_017583883.1 | Aves | **39-NIS**, 54-NIT, 271-NMT, 483-NHT, 495-NST, 547-NNS, 639-NIS |

| | | |
|---|---|---|
| XP_017667729.1 | Aves | **39-NIS**, 54-NIT, 80-NAS, 196-NYS, 324-NMT, 536-NHT, 548-NST, 582-NAT, 600-NNS, 692-NIS |
| XP_017939494.2 | Aves | **39-NIS**, 54-NIT, 80-NAS, 196-NYS, 324-NMT, 536-NHT, 548-NST, 582-NAT, 600-NNS, 692-NIS |
| XP_018418558.1 | Amphibia | 53-NIS, 75-NAS, 136-NGT, 329-NKS, 546-NST, 677-NQT |
| XP_018539189.1 | Actinopterygii | 54-NIT, 158-NYS, 331-NNS, 390-NQS, 435-NET, 606-NRT |
| XP_018584732.1 | Actinopterygii | **37-NAT**, 55-NIT, 216-NYS, 390-NLS, 435-NET, 549-NST, 797-NKS |
| XP_018874749.1 | Mammalia | 53-NIT, 90-NLT, 103-NGS, 322-NMT, 432-NET, 546-NST, 690-NVS |
| XP_019273508.1 | Mammalia | 53-NIT, 90-NTT, 216-NYS, 299-NQS, 322-NMT, 546-NSS, 580-NVT, 660-NQT, 690-NVS |
| XP_019350687.1 | Reptilia | **19-NVT**, 48-NIT, 74-NAS, 301-NAS, 384-NLT, 543-NST, 577-NAT, 595-NNT, 604-NTT, 687-NTS |
| XP_019381060.1 | Reptilia | **19-NVT**, 48-NIT, 74-NAS, 81-NET, 325-NNS, 382-NLT, 427-NDT, 541-NST, 575-NAT, 655-NET |
| XP_019384826.1 | Reptilia | **19-NVT**, 48-NIT, 74-NAS, 325-NNS, 541-NST, 575-NAT, 593-NNT, 602-NTT, 685-NIS |
| XP_019467554.1 | Aves | 52-NIT, 78-NAS, 194-NYS, 280-NLT, 322-NMT, 546-NST, 721-NGS, 746-NIS, 765-NVT, 782-NTT |
| XP_019522936.1 | Mammalia | 53-NIT, 90-NAT, 322-NMT, 432-NET, 546-NST, 601-NSS, 661-NQT |
| XP_019742561.1 | Actinopterygii | 54-NIT, 158-NYS, 329-NNS, 388-NLS, 428-NFT, 662-NNT |
| XP_019781177.2 | Mammalia | 53-NIT, 90-NLT, 298-NQS, 321-NMT, 545-NST, 614-NQS, 659-NKT, 689-NMS |
| XP_019811719.1 | Mammalia | 53-NIT, 90-NLT, 298-NQS, 431-NET, 545-NST, 659-NET, 689-NVS |
| XP_019935235.1 | Actinopterygii | 55-NYS, 66-NIT, 343-NNS, 482-NIS, 618-NRT, 676-NQT, 706-NTS |
| XP_020140826.1 | Mammalia | 79-NIT, 116-NLT, 244-NRS, 348-NMT, 572-NST, 606-NVT, 635-NWS, 736-NVS |
| XP_020493627.1 | Actinopterygii | **43-NYS**, 331-NNS, 389-NLS, 469-NIT, 663-NKT |
| XP_020642422.1 | Reptilia | 51-NLS, 66-NIT, 103-NET, 167-NST, 317-NVT, 343-NNS, 446-NET, 560-NST, 594-NAS, 612-NNT, 672-NKT, 704-NTT, 800-NTS |
| XP_020768965.1 | Mammalia | 53-NIT, 90-NLT, 298-NQS, 431-NET, 545-NST |
| XP_020781598.1 | Actinopterygii | **44-NYS**, 391-NLS, 550-NST, 666-NTT |
| XP_021009138.1 | Mammalia | 53-NIT, 546-NST, 634-NWT, 690-NVS, 772-NET |
| XP_021043935.1 | Mammalia | 53-NIT, 82-NFS, 299-NQS, 546-NST, 660-NQT, 690-NVS, 772-NET |
| XP_021178197.1 | Actinopterygii | **38-NAS**, 307-NET, 392-NLS, 670-NFT |
| XP_021240731.1 | Aves | 54-NIT, 80-NAS, 196-NYS, 282-NLT, 324-NMT, 548-NST, 582-NAT, 600-NNS, 692-NVS |
| XP_021388026.1 | Aves | **37-NIS**, 52-NIT, 78-NAS, 534-NHT, 546-NST, 598-NNS, 690-NIT |
| XP_021433278.1 | Actinopterygii | **37-NAT**, 391-NLS, 550-NST, 665-NKT, 794-NKT |
| XP_021536480.1 | Mammalia | 53-NIT, 75-NQS, 216-NYS, 299-NQS, 322-NMT, 546-NSS, 690-NVS, 785-NLS |
| XP_021788732.1 | Mammalia | 53-NIT, 90-NLT, 103-NGS, 322-NMT, 432-NET, 546-NST, 690-NVS |
| XP_022063988.1 | Actinopterygii | **36-NAT**, 54-NIT, 289-NQT, 331-NNS, 390-NLS, 430-NFT, 435-NYT, 470-NIT, 537-NHT, 606-NRT, 664-NQS |
| XP_022374078.1 | Mammalia | 53-NIT, 216-NYS, 299-NQS, 322-NMT, 690-NMS |
| XP_022418360.1 | Mammalia | 53-NIT, 90-NLT, 298-NQS, 321-NMT, 545-NST, 659-NET, 689-NMS, 790-NNS |
| XP_022523929.1 | Actinopterygii | 52-NIT, 318-NMS, 546-NST |
| XP_022605054.1 | Actinopterygii | 54-NIT, 158-NYS, 331-NDS, 390-NLS, 435-NET, 470-NIS, 664-NES |
| XP_023054821.1 | Mammalia | 53-NIT, 90-NLT, 103-NGS, 322-NMT, 432-NET, 546-NST, 690-NVS |
| XP_023104564.1 | Mammalia | 53-NIT, 90-NTT, 218-NYS, 301-NQS, 324-NMT, 548-NSS, 582-NVT, 662-NQT, 692-NVS |
| XP_023124156.1 | Actinopterygii | 54-NIT, 390-NLS, 435-NDT, 470-NIT, 664-NQS |
| XP_023257445.1 | Actinopterygii | 54-NIT, 158-NYS, 331-NDS, 390-NLS, 435-NET, 470-NIS, 664-NES |
| XP_023410960.1 | Mammalia | 53-NIT, 154-NST, 252-NQT, 298-NAT, 541-NST, 596-NSS, 655-NQT, 685-NAS |
| XP_023575315.1 | Mammalia | 79-NIT, 116-NLT, 458-NET, 572-NST, 716-NIS, 817-NNT |
| XP_023609437.1 | Mammalia | 53-NIT, 90-NST, 103-NGS, 215-NYS, 279-NLT, 328-NNS, 431-NET, 674-NQT, 704-NMS, 799-NLS |
| XP_023774184.1 | Aves | **24-NVT**, 57-NIT, 83-NAS, 140-NNS, 141-NSS, 539-NHT, 551-NST, 603-NNS |
| XP_023964517.1 | Reptilia | 52-NIT, 78-NAS, 329-NNS, 387-NLS, 432-NET, 534-NHT, 546-NST, 580-NAT, 598-NNS, 633-NSS, 710-NVS |
| XP_023971279.1 | Mammalia | 53-NIT, 90-NLT, 298-NQS, 321-NMT, 545-NST, 659-NET, 689-NVS, 790-NNS |
| XP_024150631.1 | Actinopterygii | 58-NIT, 394-NLS, 474-NIS |
| XP_024425698.1 | Mammalia | 53-NIT, 279-NLT, 386-NQS, 431-NET, 545-NST, 659-NQT, 784-NLS |
| XP_024599894.1 | Mammalia | 53-NIT, 298-NQS, 321-NMT, 545-NST, 600-NSS, 659-NET, 689-NMS, 790-NNS |
| XP_025066628.1 | Reptilia | 48-NIT, 74-NAS, 299-NAS, 382-NLT, 575-NAT, 593-NNT, 602-NTT, 685-NTS |
| XP_025227847.1 | Mammalia | 53-NIT, 90-NLT, 103-NGS, 322-NMT, 432-NET, 546-NST, 690-NVS |
| XP_025292925.1 | Mammalia | 52-NIT, 215-NYS, 298-NQS, 321-NMT, 545-NSS, 659-NQT, 689-NVS |
| XP_025713397.1 | Mammalia | 53-NIT, 216-NYS, 299-NQS, 322-NMT, 432-NET, 546-NSS, 690-NMS, 785-NLS |
| XP_025790417.1 | Mammalia | 53-NIT, 90-NTT, 216-NYS, 299-NQS, 322-NMT, 580-NVT, 690-NVS |
| XP_025842512.1 | Mammalia | 52-NIS, 215-NYS, 298-NQS, 321-NMT, 545-NSS, 689-NVS |
| XP_025891105.1 | Aves | 52-NIT, 78-NAS, 322-NMT, 329-NNS, 467-NIT, 546-NST, 580-NAT, 598-NNS, 636-NSS, 690-NVS, 777-NST |
| XP_025942946.1 | Aves | 52-NIT, 78-NAS, 322-NMT, 329-NYS, 467-NIT, 546-NST, 572-NIT, 580-NAT, 598-NNS, 690-NVS |
| XP_026020155.1 | Actinopterygii | 54-NIT, 390-NLS, 470-NIT, 662-NKT |
| XP_026131313.1 | Actinopterygii | 320-NMS, 388-NHS, 547-NST, 785-NKT |
| XP_026175949.1 | Actinopterygii | 54-NIT, 331-NNS, 390-NLS, 430-NFT, 435-NET, 470-NIT, 606-NRT, 664-NES |
| XP_026233431.1 | Actinopterygii | 155-NST, 331-NNS, 390-NQS, 664-NET |
| XP_026252505.1 | Mammalia | 63-NIT, 100-NFT, 164-NST, 332-NMT, 442-NET, 556-NST, 670-NQT, 700-NVS, 782-NQT |
| XP_026333865.1 | Mammalia | 53-NIT, 90-NST, 216-NYS, 322-NMT, 432-NET, 546-NSS, 690-NVS |
| XP_026570054.1 | Reptilia | 66-NAS, 77-NLT, 114-NET, 178-NYS, 309-NKT, 344-NMT, 631-NWT, 682-NKT, 712-NIS, 815-NST |
| XP_026705725.1 | Aves | 77-NIT, 103-NAS, 178-NST, 354-NNS, 559-NHT, 623-NNS, 715-NIS |
| XP_026803610.1 | Actinopterygii | 52-NIT, 318-NMS, 546-NST, 692-NAS, 789-NGT |
| XP_026867211.2 | Actinopterygii | 52-NIT, 77-NES, 546-NAT, 692-NAS, 798-NDS |
| XP_026910297.1 | Mammalia | 53-NIT, 90-NTT, 216-NYS, 299-NQS, 322-NMT, 546-NSS, 580-NVT, 690-NVS |
| XP_026951598.1 | Mammalia | 53-NIT, 90-NLT, 298-NQS, 321-NMT, 545-NST, 614-NQS, 635-NDS, 659-NKT, 689-NMS |
| XP_027024524.1 | Actinopterygii | 53-NIT, 319-NMS, 329-NNS, 388-NLS, 547-NST, 693-NAT, 799-NES |
| XP_027389727.1 | Mammalia | 53-NIT, 90-NLT, 298-NQS, 431-NET, 545-NST, 659-NET, 689-NVS |
| XP_027465353.1 | Mammalia | 53-NIT, 216-NYS, 299-NQS, 322-NMT, 432-NET, 546-NSS, 690-NMS, 785-NLS |
| XP_027494818.1 | Aves | **39-NIS**, 54-NIT, 80-NAS, 196-NYS, 324-NMT, 536-NHT, 548-NST, 582-NAT, 600-NNS, 692-NIS |
| XP_027544864.1 | Aves | 80-NAS, 196-NYS, 305-NAT, 536-NHT, 548-NST, 582-NAT, 600-NNS, 692-NIS |
| XP_027593974.1 | Aves | **39-NIS**, 54-NIT, 80-NAS, 196-NYS, 324-NMT, 536-NHT, 548-NST, 582-NAT, 600-NNS, 692-NIS |
| XP_027691156.1 | Mammalia | 80-NIS, 324-NMT, 548-NST, 582-NAT, 692-NGS, 773-NKS |

| | | |
|---|---|---|
| XP_027757151.1 | Aves | 52-NIT, 78-NAS, 194-NYS, 303-NAT, 546-NST, 580-NAT, 598-NNS, 690-NIS |
| XP_027802308.1 | Mammalia | 63-NIT, 100-NFT, 164-NST, 332-NMT, 442-NET, 556-NST, 619-NWS, 670-NQT, 700-NAS, 782-NQT |
| XP_027871671.1 | Actinopterygii | **37-NAS**, 391-NLS, 471-NIS |
| XP_027970822.1 | Mammalia | 53-NIT, 216-NYS, 299-NQS, 322-NMT, 432-NET, 546-NSS, 690-NMS, 785-NLS |
| XP_028020351.1 | Mammalia | 53-NIT, 90-NLT, 298-NQS, 321-NMT, 431-NVT, 545-NST, 659-NET, 689-NVS, 790-NNS |
| XP_028257887.1 | Actinopterygii | 55-NIT, 159-NYS, 332-NNS, 391-NLS, 436-NET, 471-NIT, 550-NST, 679-NET |
| XP_028297875.1 | Actinopterygii | 48-NAS, 66-NIT, 170-NYS, 229-NYT, 342-NNS, 401-NLS, 446-NET, 560-NST, 675-NET, 689-NET |
| XP_028378317.1 | Mammalia | 53-NIT, 319-NMT, 429-NET, 491-NET, 543-NST, 688-NDT, 784-NLS |
| XP_028441363.1 | Actinopterygii | 155-NST, 331-NNS, 390-NLS, 470-NIT, 549-NST, 606-NRT, 630-NSS, 676-NKT |
| XP_028617961.1 | Mammalia | 53-NIT, 82-NFS, 546-NST, 572-NVT, 615-NQS, 660-NQT, 690-NAS |
| XP_028655640.1 | Actinopterygii | 52-NIT, 78-NAT, 89-NYT, 150-NST, 318-NMT, 325-NNS, 530-NHS, 542-NST, 576-NAT, 662-NFT, 685-NQS, 786-NGS |
| XP_028743609.1 | Mammalia | 53-NIT, 82-NYS, 546-NST, 658-NKT, 690-NVS, 772-NET |
| XP_028837781.1 | Actinopterygii | **35-NAT**, 53-NIT, 330-NNS, 390-NLT, 549-NST, 796-NNS |
| XP_028999570.1 | Actinopterygii | 61-NIS, 397-NLS, 671-NQS |
| XP_029095804.1 | Mammalia | 53-NIT, 90-NLT, 298-NQS, 321-NMT, 545-NST, 659-NET, 689-NMS, 790-NNS |
| XP_029140508.1 | Reptilia | **41-NAS**, 52-NIT, 78-NAS, 89-NET, 153-NYS, 284-NKT, 319-NMT, 543-NST, 595-NNS, 606-NWT, 657-NKT, 687-NIS |
| XP_029283581.1 | Actinopterygii | 54-NIT, 331-NKS, 390-NLS, 430-NFT, 470-NIT, 534-NAS, 664-NET, 694-NLS |
| XP_029354066.1 | Actinopterygii | **43-NYS**, 54-NIT, 158-NYS, 331-NNS, 390-NLS, 435-NET, 470-NIT, 673-NQT, 687-NLT |
| XP_029459086.1 | Amphibia | 52-NIT, 78-NSS, 257-NKT, 301-NWT, 387-NLS, 432-NET, 534-NHS, 546-NST, 598-NNT, 662-NQT, 692-NLS |
| XP_029702274.1 | Actinopterygii | 26-NQT, **45-NYS**, 56-NIT, 81-NES, 160-NYS, 307-NET, 432-NFT, 472-NIT, 539-NHT, 608-NRT, 666-NET |
| XP_029786256.1 | Mammalia | 53-NIT, 90-NTT, 216-NYS, 546-NST, 690-NVS |
| XP_029855025.1 | Aves | **37-NIS**, 52-NIT, 78-NAS, 153-NST, 329-NNS, 534-NHT, 546-NSS, 598-NNS, 690-NIS |
| XP_029904152.1 | Actinopterygii | 54-NIT, 331-NNS, 390-NLS, 470-NIT, 537-NHT, 664-NMT |
| XP_029949252.1 | Actinopterygii | 54-NIT, 62-NVT, 158-NYS, 389-NLS, 429-NFT, 533-NAS, 672-NRT |
| XP_030058174.1 | Amphibia | **19-NVT**, 52-NIT, 78-NSS, 153-NST, 387-NLS, 546-NST, 580-NST, 598-NNT, 617-NDS, 662-NQT, 676-NET, 792-NIT |
| XP_030160839.1 | Mammalia | 53-NIT, 90-NTT, 216-NYS, 299-NQS, 322-NMT, 546-NSS, 580-NVT, 660-NQT, 690-NVS |
| XP_030232530.1 | Actinopterygii | 228-NYT, 400-NLS, 559-NST, 675-NKT, 805-NLS |
| XP_030332639.1 | Aves | **39-NIS**, 54-NIT, 80-NAS, 155-NST, 196-NYS, 331-NNS, 536-NHT, 600-NNS, 692-NIS |
| XP_030407881.1 | Reptilia | 52-NIT, 78-NAS, 329-NNS, 387-NLS, 432-NET, 534-NHT, 546-NST, 580-NAT, 598-NNS, 660-NQT, 690-NTS |
| XP_030582139.1 | Actinopterygii | 54-NIT, 390-NLS, 470-NIT, 549-NST, 662-NET, 796-NDS |
| XP_030627971.1 | Actinopterygii | **34-NAT**, 52-NIT, 328-NNS, 432-NET, 546-NST, 575-NHT |
| XP_030703991.1 | Mammalia | 53-NIT, 90-NLT, 298-NQS, 321-NMT, 545-NST, 614-NQS, 659-NKT, 689-NMS |
| XP_030811385.1 | Aves | 52-NIT, 78-NAS, 329-NNS, 546-NST, 598-NNS, 690-NIS |
| XP_031162227.1 | Actinopterygii | 54-NIS, 331-NNS, 390-NLS, 470-NIS, 549-NST, 606-NRT, 664-NKT, 678-NET |
| XP_031226742.1 | Mammalia | 54-NIT, 83-NFS, 300-NQS, 547-NST, 691-NVS, 773-NET |
| XP_031414786.1 | Actinopterygii | 60-NIT, 336-NDS, 395-NLS, 440-NET, 554-NST |
| XP_031451919.1 | Aves | 52-NIT, 78-NAS, 194-NYS, 280-NLT, 322-NMT, 534-NHT, 546-NST, 580-NAT, 598-NNS, 690-NVS |
| XP_031584810.1 | Actinopterygii | 390-NLS, 470-NIT, 681-NKT, 710-NAT |
| XP_031702716.1 | Actinopterygii | 62-NIT, 84-NMS, 98-NDT, 398-NLS, 443-NDT, 557-NST, 614-NRT, 672-NES |
| XP_031956594.1 | Aves | **39-NIS**, 54-NIT, 80-NAS, 324-NMT, 536-NHT, 548-NST, 600-NNS, 692-NIS |
| XP_032058386.1 | Aves | 52-NIT, 78-NAS, 322-NMT, 534-NHT, 546-NST, 580-NAT, 598-NNS, 687-NVS |
| XP_032082934.1 | Reptilia | 77-NLT, 103-NAS, 114-NET, 130-NSS, 176-NWS, 309-NKT, 351-NNS, 631-NWT, 656-NWT, 682-NKT, 712-NIS, 815-NST |
| XP_032141854.1 | Mammalia | 53-NIT, 90-NLT, 103-NGS, 322-NMT, 609-NWS, 690-NVS |
| XP_032187677.1 | Mammalia | 53-NIT, 299-NQS, 322-NMT, 660-NQT, 690-NMS |
| XP_032245506.1 | Mammalia | 53-NIT, 216-NYS, 322-NMT, 432-NET, 546-NSS, 660-NQT, 690-NVS, 785-NLS |
| XP_032355526.1 | Actinopterygii | 54-NIT, 331-NNS, 390-NLS, 470-NIT, 549-NST, 606-NRT, 664-NET |
| XP_032417398.1 | Actinopterygii | **37-NAS**, 391-NLS, 471-NIS |
| XP_032476001.1 | Mammalia | 53-NIT, 298-NQS, 321-NMT, 545-NST, 600-NSS, 659-NET, 689-NMS, 790-NNS |
| XP_032536093.1 | Aves | 54-NIT, 80-NAS, 196-NYS, 324-NMT, 536-NHT, 548-NST, 582-NAT, 692-NIS |
| XP_032612508.1 | Mammalia | 53-NIT, 90-NLT, 103-NGS, 322-NMT, 432-NET, 546-NST, 690-NVS, 772-NKS |
| XP_032631197.1 | Reptilia | 52-NIT, 78-NAS, 153-NST, 329-NNS, 387-NLS, 432-NET, 534-NHT, 546-NST, 580-NAT, 598-NNS, 660-NQT, 690-NTS |
| XP_032736028.1 | Mammalia | 53-NIT, 216-NYS, 322-NMT, 690-NMS |
| XP_032746145.1 | Mammalia | 53-NIT, 82-NFS, 299-NQS, 546-NST, 601-NST, 648-NQT, 678-NVS, 760-NET |
| XP_032865981.1 | Aves | 52-NIT, 78-NAS, 329-NNS, 534-NHT, 598-NNS, 690-NIS |
| XP_032888812.1 | Chondrichthyes | 78-NAS, 280-NFS, 329-NYS, 388-NQT, 535-NQT, 547-NST, 581-NST, 693-NIT |
| XP_032907631.1 | Aves | 7-NRS, 101-NIS, 116-NIT, 142-NAS, 393-NNS, 598-NHT, 610-NST, 644-NAT, 662-NNS, 754-NVT |
| XP_032963186.1 | Mammalia | **38-NLS**, 53-NIS, 82-NFS, 90-NDT, 322-NMT, 329-NNS, 432-NET, 546-NST, 660-NQT, 690-NLS |
| XP_033806113.1 | Amphibia | 52-NIT, 78-NSS, 153-NST, 299-NKS, 329-NNS, 432-NET, 546-NST, 598-NNT, 617-NDS, 676-NET |
| XP_034290793.1 | Reptilia | **66-NAS**, 77-NLT, 103-NAS, 114-NET, 126-NGS, 178-NYS, 309-NKT, 344-NMT, 568-NST, 620-NNS, 631-NWT, 682-NKT, 711-NNT, 712-NTS, 815-NST |
| XP_034341939.1 | Mammalia | 53-NIT, 82-NFS, 299-NQS, 546-NST, 572-NVT, 660-NQT, 690-NAS, 772-NET |
| XP_034505781.1 | Mammalia | 53-NIT, 90-NST, 216-NYS, 322-NMT, 432-NET, 546-NSS, 690-NVS |
| XP_034612285.1 | Reptilia | 52-NIT, 78-NAS, 329-NNS, 387-NLS, 432-NET, 534-NHT, 546-NST, 580-NAT, 598-NNS, 633-NSS, 680-NQT, 710-NVS |
| XP_034852450.1 | Mammalia | 53-NIT, 216-NYS, 299-NQS, 322-NMT, 432-NET, 690-NVS, 785-NLS |
| XP_034971718.1 | Reptilia | 52-NIT, 78-NAS, 303-NVT, 329-NNS, 387-NLS, 546-NST, 690-NGT |
| XP_035182951.1 | Aves | **19-NVT**, 52-NIT, 78-NAS, 322-NMT, 546-NST, 572-NLT, 598-NNS, 690-NVS |
| XP_035398116.1 | Aves | 52-NIT, 78-NAS, 322-NMT, 546-NST, 572-NLT, 580-NAT, 598-NNS, 690-NVS |
| XP_416822.2 | Aves | 52-NIT, 78-NAS, 194-NYS, 280-NLT, 322-NMT, 534-NHT, 546-NST, 580-NAT, 598-NNS, 690-NVS |

**Table S7**. Comparison of the interface residues in the 285 ACE2 proteins. Interface residue positions are indexed by hACE2. '-' stands for a gap, and '*' for the same amino acid type as that of hACE2.

| Species | 24 | 27 | 28 | 30 | 31 | 34 | 35 | 37 | 38 | 41 | 42 | 79 | 82 | 83 | 330 | 353 | 354 | 355 | 357 | 393 |
|---|---|---|---|---|---|---|---|---|---|---|---|---|---|---|---|---|---|---|---|---|
| *Homo sapiens* | Q | T | F | D | K | H | E | E | D | Y | Q | L | M | Y | N | K | G | D | R | R |
| *Pongo abelii* | * | * | * | * | * | * | * | * | * | * | * | * | * | * | * | * | * | * | * | * |
| *Pan paniscus* | * | * | * | * | * | * | * | * | * | * | * | * | * | * | * | * | * | * | * | * |
| *Nomascus leucogenys* | * | * | * | * | * | * | * | * | * | * | * | * | * | * | * | * | * | * | * | * |
| *Gorilla gorilla* | * | * | * | * | * | * | * | * | * | * | * | * | * | * | * | * | * | * | * | * |
| *Papio anubis* | * | * | * | * | * | * | * | * | * | * | * | * | * | * | * | * | * | * | * | * |
| *Hylobates moloch* | * | * | * | * | * | * | * | * | * | * | * | * | * | * | * | * | * | * | * | * |
| *Chlorocebus sabaeus* | * | * | * | * | * | * | * | * | * | * | * | * | * | * | * | * | * | * | * | * |
| *Macaca nemestrina* | * | * | * | * | * | * | * | * | * | * | * | * | * | * | * | * | * | * | * | * |
| *Macaca fascicularis* | * | * | * | * | * | * | * | * | * | * | * | * | * | * | * | * | * | * | * | * |
| *Theropithecus gelada* | * | * | * | * | * | * | * | * | * | * | * | * | * | * | * | * | * | * | * | * |
| *Macaca mulatta* | * | * | * | * | * | * | * | * | * | * | * | * | * | * | * | * | * | * | * | * |
| *Cercocebus atys* | * | * | * | * | * | * | * | * | * | * | * | * | * | * | * | * | * | * | * | * |
| *Rhinopithecus roxellana* | * | * | * | * | * | * | * | * | * | * | * | * | * | * | * | * | * | * | * | * |
| *Pan troglodytes* | * | * | * | * | * | * | * | * | * | * | * | * | * | * | * | * | * | * | * | * |
| *Mandrillus leucophaeus* | * | * | * | * | * | * | * | * | * | * | * | * | * | * | * | * | * | * | * | * |
| *Piliocolobus tephrosceles* | * | * | * | * | * | * | * | * | * | * | * | * | * | * | * | * | * | * | * | * |
| *Mesocricetus auratus* | * | * | * | * | * | Q | * | * | * | * | * | * | N | * | * | * | * | * | * | * |
| *Cricetulus griseus* | * | * | * | * | * | Q | * | * | * | * | * | * | N | * | * | * | * | * | * | * |
| *Nannospalax galili* | * | * | * | * | * | Q | * | * | * | * | * | * | K | * | * | * | * | * | * | * |
| *Eumetopias jubatus* | L | * | * | E | * | S | * | * | E | * | * | Q | T | * | * | * | H | * | * | * |
| *Propithecus coquereli* | * | * | * | * | * | * | * | * | * | * | * | * | T | * | * | * | * | * | * | * |
| *Callorhinus ursinus* | L | * | * | E | * | S | * | * | E | * | * | Q | T | * | * | * | H | * | * | * |
| *Equus caballus* | L | * | * | E | * | S | * | * | E | H | * | * | T | * | * | * | * | * | * | * |
| *Equus przewalskii* | L | * | * | E | * | S | * | * | E | H | * | * | T | * | * | * | * | * | * | * |
| *Panthera tigris altaica* | L | * | * | E | * | * | * | * | E | * | * | * | T | * | * | * | * | * | * | * |
| *Acinonyx jubatus* | L | * | * | E | * | * | * | * | E | * | * | * | T | * | K | * | * | * | * | * |
| *Capra hircus* | * | * | * | E | * | * | * | * | * | * | * | M | T | * | * | * | * | * | * | * |
| *Oryctolagus cuniculus* | L | * | * | E | * | Q | * | * | * | * | * | * | T | * | * | * | * | * | * | * |
| *Bos mutus* | * | * | * | E | * | * | * | * | * | * | * | M | T | * | * | * | * | * | * | * |
| *Puma concolor* | L | * | * | E | * | * | * | * | E | * | * | * | T | * | * | * | * | * | * | * |
| *Heterocephalus glaber* | * | * | * | * | * | Q | * | * | * | * | * | * | A | * | * | * | D | * | * | * |
| *Bison bison bison* | * | * | * | E | * | * | * | * | * | * | * | M | T | * | * | * | * | * | * | * |
| *Panthera pardus* | L | * | * | E | * | * | * | * | E | * | * | * | T | * | * | * | * | * | * | * |
| *Mustela erminea* | L | * | * | E | * | Y | * | * | E | * | * | H | T | * | * | * | R | * | * | * |
| *Phoca vitulina* | L | * | * | E | * | Y | * | * | E | * | * | Q | T | * | * | * | R | * | * | * |
| *Bos taurus* | * | * | * | E | * | * | * | * | * | * | * | M | T | * | * | * | * | * | * | * |
| *Bos indicus x Bos taurus* | * | * | * | E | * | * | * | * | * | * | * | M | T | * | * | * | * | * | * | * |
| *Lontra canadensis* | L | * | * | E | * | Y | * | * | E | * | * | N | T | * | * | * | R | * | * | * |
| *Odobenus rosmarus divergens* | L | * | * | E | * | Y | * | * | E | * | * | Q | T | * | * | * | H | * | * | * |
| *Neomonachus schauinslandi* | L | * | * | E | * | Y | * | * | E | * | * | Q | T | * | * | * | H | * | * | * |
| *Mustela putorius furo* | L | * | * | E | * | Y | * | * | E | * | * | H | T | * | * | * | R | * | * | * |
| *Zalophus californianus* | L | * | * | E | * | S | * | * | E | * | * | Q | T | * | * | * | H | * | * | * |
| *Bos indicus* | * | * | * | E | * | * | * | * | * | * | * | M | T | * | * | * | * | * | * | * |
| *Bubalus bubalis* | * | * | * | E | * | * | * | * | * | * | * | M | T | * | * | * | * | * | * | * |
| *Jaculus jaculus* | M | * | * | * | * | Q | * | * | * | * | * | V | T | * | * | * | N | * | * | * |
| *Peromyscus maniculatus bairdii* | * | I | * | * | * | Q | * | * | * | * | * | * | N | * | * | * | * | * | * | * |
| *Felis catus* | L | * | * | E | * | * | * | * | E | * | * | * | T | * | * | * | * | * | * | * |
| *Odocoileus virginianus texanus* | * | * | * | E | * | * | * | * | * | * | * | M | T | * | * | * | * | * | * | * |
| *Fukomys damarensis* | * | * | * | * | * | Q | * | * | * | * | * | * | A | * | * | * | N | * | * | * |
| *Lynx canadensis* | L | * | * | E | * | * | * | * | E | * | * | * | T | * | * | * | * | * | * | * |
| *Ailuropoda melanoleuca* | L | * | * | E | * | Y | * | * | * | * | * | H | T | * | * | * | * | * | * | * |
| *Peromyscus leucopus* | * | I | * | * | * | Q | * | * | * | * | * | * | N | * | * | * | * | * | * | * |
| *Ursus arctos horribilis* | L | * | * | E | * | Y | * | * | * | * | * | H | T | * | * | * | * | * | * | * |
| *Ovis aries* | * | * | * | E | * | * | * | * | * | * | * | M | T | * | * | * | * | * | * | * |
| *Delphinapterus leucas* | * | * | * | Q | * | * | * | * | * | * | * | I | T | * | * | * | * | * | * | * |
| *Monodon monoceros* | * | * | * | Q | * | * | * | * | * | * | * | I | T | * | * | * | * | * | * | * |
| *Phocoena sinus* | * | * | * | Q | * | * | * | * | * | * | * | I | T | * | * | * | * | * | * | * |
| *Physeter macrocephalus* | * | * | * | Q | * | * | * | * | * | * | * | T | T | * | * | * | * | * | * | * |
| *Ursus maritimus* | L | * | * | E | * | Y | * | * | * | * | * | H | T | * | * | * | * | * | * | * |
| *Neophocaena asiaeorientalis asiaeorientalis* | * | * | * | Q | * | * | * | * | * | * | * | I | T | * | * | * | * | * | * | * |
| *Microtus ochrogaster* | D | A | * | * | * | Q | * | * | * | * | * | * | S | * | * | * | D | * | * | * |
| *Manis javanica* | E | * | * | E | * | S | * | * | E | * | * | I | N | * | * | * | H | * | * | * |
| *Balaenoptera acutorostrata scammoni* | * | * | * | Q | * | * | * | * | * | * | R | I | T | * | * | * | * | * | * | * |
| *Marmota marmota* | L | * | * | * | * | Q | * | * | * | * | * | * | A | * | * | * | * | * | * | * |
| *Vulpes vulpes* | L | * | * | E | * | Y | * | * | E | * | * | * | T | * | * | * | * | * | * | * |
| *Ictidomys tridecemlineatus* | L | * | * | * | * | Q | * | * | * | * | * | * | A | * | * | * | * | * | * | * |
| *Marmota flaviventris* | L | * | * | * | * | Q | * | * | * | * | * | * | A | * | * | * | * | * | * | * |
| *Canis lupus familiaris* | L | * | * | E | * | Y | * | * | E | * | * | * | T | * | * | * | * | * | * | * |
| *Canis lupus dingo* | L | * | * | E | * | Y | * | * | E | * | * | * | T | * | * | * | * | * | * | * |
| *Ceratotherium simum simum* | L | * | * | E | * | P | * | * | * | * | * | * | T | * | * | * | * | * | * | * |
| *Ochotona princeps* | L | * | * | * | * | Q | * | * | * | * | * | * | T | * | * | * | D | * | * | * |
| *Rousettus aegyptiacus* | L | * | * | E | * | T | * | * | * | * | * | * | T | * | K | * | * | * | * | * |
| *Sus scrofa* | L | * | * | E | * | L | * | * | * | * | * | I | T | * | * | * | * | * | * | * |
| *Urocitellus parryii* | L | * | * | * | * | Q | * | * | * | H | * | * | D | * | * | * | * | * | * | * |
| *Lagenorhynchus obliquidens* | R | * | * | Q | * | R | * | * | * | * | * | I | T | * | * | * | * | * | * | * |

| | | | | | | | | | | | | | | | | | | | | |
|---|---|---|---|---|---|---|---|---|---|---|---|---|---|---|---|---|---|---|---|---|
| *Nyctereutes procyonoides* | L | * | * | E | * | Y | * | * | E | * | * | * | T | * | * | R | * | * | * | * |
| *Equus asinus* | L | * | * | E | * | S | * | * | E | H | * | * | T | * | * | * | * | * | * | * |
| *Rhinolophus sinicus* | E | M | * | * | * | T | K | * | * | H | * | * | N | * | * | * | * | * | * | * |
| *Camelus bactrianus* | L | * | * | E | E | * | * | * | * | * | * | T | T | * | * | * | * | * | * | * |
| *Mirounga leonina* | L | K | * | E | * | Y | * | * | E | * | * | Q | T | * | * | * | H | * | * | * |
| *Camelus dromedarius* | L | * | * | E | E | * | * | * | * | * | * | T | T | * | * | * | * | * | * | * |
| *Phyllostomus discolor* | D | K | * | E | N | N | * | * | E | * | * | * | N | * | * | * | K | * | * | * |
| *Camelus ferus* | L | * | * | E | E | * | * | * | * | * | * | T | T | * | * | * | * | * | * | * |
| *Orcinus orca* | R | * | * | Q | * | R | * | * | * | * | * | I | T | * | * | * | * | * | * | * |
| *Pteropus vampyrus* | L | * | * | E | * | T | * | * | * | * | * | * | A | * | K | * | * | * | * | K |
| *Pteropus alecto* | L | * | * | E | * | T | * | * | * | * | * | * | A | * | K | * | * | * | * | K |
| *Globicephala melas* | R | * | * | Q | * | R | * | * | * | * | * | I | T | * | * | * | * | * | * | * |
| *Tursiops truncatus* | R | * | * | Q | * | R | * | * | * | * | * | I | T | * | * | * | * | * | * | * |
| *Lipotes vexillifer* | R | * | * | Q | * | * | * | * | * | * | * | I | T | F | * | * | * | * | * | * |
| *Dipodomys ordii* | L | * | * | * | N | Q | * | * | * | * | * | * | I | * | * | * | * | * | * | * |
| *Loxodonta africana* | L | * | * | * | T | Q | * | * | * | * | * | * | D | F | * | * | * | * | * | * |
| *Orycteropus afer* | L | * | * | E | * | Q | * | * | N | * | * | I | S | F | K | * | * | * | * | * |
| *Enhydra lutris kenyoni* | P | * | * | E | * | Y | * | * | E | * | * | H | T | * | * | * | R | * | * | * |
| *Trichechus manatus latirostris* | L | * | * | * | T | Q | * | * | * | * | * | * | N | F | * | * | * | * | * | * |
| *Paguma larvata* | L | * | * | E | T | Y | * | Q | E | * | * | * | T | * | * | * | * | * | * | * |
| *Octodon degus* | * | * | * | * | N | Q | K | * | * | * | * | * | A | * | * | * | N | * | * | * |
| *Vicugna pacos* | L | * | * | K | E | * | * | * | * | * | * | A | I | * | * | * | * | * | * | * |
| *Callithrix jacchus* | * | * | * | * | * | * | * | * | * | H | E | * | T | * | * | * | Q | * | * | * |
| *Saimiri boliviensis* | * | * | * | * | * | * | * | * | * | H | E | * | T | * | * | * | Q | * | * | * |
| *Otolemur garnettii* | * | * | * | * | N | R | * | * | E | H | * | I | T | * | * | * | D | * | * | * |
| *Eptesicus fuscus* | N | I | * | E | N | S | * | * | * | H | E | * | T | * | * | * | N | * | * | * |
| *Myotis brandtii* | K | I | * | E | N | S | K | * | * | H | E | * | T | * | * | * | * | * | * | * |
| *Aotus nancymaae* | * | * | * | * | * | * | * | * | * | H | E | * | T | * | * | * | Q | * | * | * |
| *Cyanistes caeruleus* | E | E | * | E | E | R | R | * | * | * | E | N | N | F | * | * | N | * | * | * |
| *Sapajus apella* | * | * | * | * | * | * | * | * | * | H | E | * | T | * | * | * | Q | * | * | * |
| *Myotis davidii* | K | I | * | * | N | S | K | * | * | H | E | * | T | * | * | * | * | * | * | * |
| *Chinchilla lanigera* | * | * | * | * | N | E | K | * | * | * | * | * | A | * | * | * | D | * | * | * |
| *Carlito syrichta* | * | * | * | * | * | Q | * | * | * | H | * | I | S | * | * | N | S | * | * | * |
| *Suricata suricatta* | L | * | * | E | Q | * | * | Q | E | * | L | R | A | * | * | * | * | * | * | * |
| *Condylura cristata* | * | K | * | E | T | R | * | * | E | * | N | D | R | F | * | * | * | * | * | * |
| *Microcebus murinus* | * | * | * | E | N | N | * | * | * | H | * | * | T | * | K | * | * | * | * | * |
| *Myotis lucifugus* | K | I | * | E | N | S | K | * | * | H | E | * | T | * | * | * | * | * | * | * |
| *Dasypus novemcinctus* | * | * | * | E | T | Q | Q | * | E | H | * | M | N | F | * | * | * | * | * | * |
| *Echinops telfairi* | * | S | * | T | T | N | * | * | N | * | * | * | K | F | K | L | N | * | * | * |
| *Nothoprocta perdicaria* | E | V | * | * | E | I | K | * | * | * | E | N | K | F | * | * | K | * | * | * |
| *Chrysochloris asiatica* | L | A | * | N | N | Q | * | * | N | H | * | * | K | F | * | * | D | * | * | * |
| *Mus pahari* | N | * | * | N | * | Q | * | * | * | * | * | T | N | F | * | H | * | * | * | * |
| *Rhinolophus ferrumequinum* | L | K | * | * | D | S | * | * | N | H | * | * | N | F | * | * | * | * | * | * |
| *Mastomys coucha* | N | * | * | N | * | Q | * | * | * | * | * | I | N | F | * | H | * | * | * | * |
| *Monodelphis domestica* | D | * | * | * | D | A | K | * | E | H | * | I | T | * | * | * | N | * | * | * |
| *Serinus canaria* | * | K | * | * | E | R | R | * | * | * | E | N | K | F | * | * | N | * | * | * |
| *Grammomys surdaster* | E | * | * | * | * | Q | * | * | * | * | * | T | N | F | * | H | * | * | * | * |
| *Hipposideros armiger* | L | E | * | * | * | T | * | * | * | H | L | R | D | * | * | * | * | * | * | * |
| *Sturnus vulgaris* | E | I | * | E | E | R | R | * | * | * | E | N | S | F | * | * | N | * | * | * |
| *Tupaia chinensis* | E | V | * | N | * | I | * | * | E | H | * | Q | R | * | K | * | N | * | * | * |
| *Calidris pugnax* | * | M | * | E | E | R | M | V | * | * | E | N | C | F | * | * | N | * | * | K |
| *Gekko japonicus* | R | E | * | E | Q | P | R | * | N | * | E | M | S | F | * | * | * | * | * | * |
| *Chrysemys picta* | E | N | * | S | Q | V | R | * | * | * | A | N | K | * | * | * | K | * | * | * |
| *Desmodus rotundus* | E | * | * | E | N | T | * | * | E | * | * | I | T | * | * | N | K | * | * | * |
| *Rattus rattus* | K | S | * | N | * | Q | * | * | * | * | * | I | N | F | * | H | * | * | * | * |
| *Mus caroli* | N | * | * | N | * | Q | * | * | * | * | * | T | S | F | * | H | * | * | * | * |
| *Geotrypetes seraphini* | E | V | * | * | Q | P | K | * | * | * | * | N | R | * | * | * | N | * | * | * |
| *Struthio camelus australis* | * | M | * | T | E | V | K | * | * | * | E | N | N | F | * | * | K | * | * | * |
| *Rattus norvegicus* | K | S | * | N | * | Q | * | * | * | * | * | I | N | F | * | H | * | * | * | * |
| *Falco cherrug* | E | M | * | E | E | R | R | * | N | * | E | N | S | F | * | * | N | * | * | * |
| *Aquila chrysaetos chrysaetos* | * | M | * | E | E | R | R | * | N | * | E | N | S | F | * | * | N | * | * | * |
| *Coturnix japonica* | E | K | * | A | E | V | R | * | * | * | E | N | R | F | * | * | N | * | * | * |
| *Pantherophis guttatus* | * | E | * | K | Q | V | R | I | * | * | N | N | * | F | * | * | N | * | * | * |
| *Danio rerio* | R | E | * | N | * | E | * | S | * | * | * | E | A | * | * | N | K | * | * | * |
| *Cygnus atratus* | * | M | * | A | E | V | R | * | * | * | E | N | S | F | * | * | N | * | * | * |
| *Miniopterus natalensis* | K | K | * | E | G | S | Q | * | * | F | E | * | I | * | * | * | * | * | * | * |
| *Rhinatrema bivittatum* | * | E | * | R | Q | Q | Q | * | * | H | E | N | R | * | * | * | N | * | * | * |
| *Aptenodytes forsteri* | * | M | * | E | E | K | R | * | N | * | E | N | S | F | * | * | N | * | * | * |
| *Mus musculus* | N | * | * | N | N | Q | * | * | * | * | * | T | S | F | * | H | * | * | * | * |
| *Larimichthys crocea* | E | V | * | E | * | K | * | T | Q | * | * | * | Q | F | * | N | E | * | * | * |
| *Anas platyrhynchos* | * | M | * | A | E | V | R | * | * | * | E | N | N | F | * | * | N | * | * | * |
| *Phasianus colchicus* | E | * | * | A | E | A | R | * | * | * | E | N | R | F | * | * | N | * | * | * |
| *Corvus moneduloides* | * | M | * | E | E | R | R | * | N | * | E | N | S | F | * | * | N | * | * | * |
| *Opisthocomus hoazin* | * | L | * | E | E | R | R | * | * | * | E | N | S | F | * | * | N | * | * | * |
| *Camarhynchus parvulus* | * | * | * | E | E | M | R | * | * | * | E | N | K | F | * | * | N | * | * | * |
| *Ictalurus punctatus* | R | E | * | Q | * | E | D | S | R | * | * | Q | A | F | * | N | E | * | * | * |
| *Chelonia mydas* | E | N | * | S | Q | V | R | * | * | * | A | N | K | * | * | * | K | * | * | * |
| *Oxyura jamaicensis* | * | M | * | A | E | V | R | * | N | * | E | N | S | F | * | * | N | * | * | * |
| *Aythya fuligula* | * | M | * | A | E | V | R | * | * | * | E | N | N | F | * | * | N | * | * | * |
| *Fundulus heteroclitus* | E | A | * | E | R | Q | N | S | E | * | * | E | K | * | * | N | E | * | * | * |

| | | | | | | | | | | | | | | | | | | | | |
|---|---|---|---|---|---|---|---|---|---|---|---|---|---|---|---|---|---|---|---|---|
| *Elephantulus edwardii* | * | A | * | E | Q | Q | Q | * | * | * | * | V | N | F | * | * | * | * | * | * |
| *Ficedula albicollis* | E | I | * | E | E | R | R | * | * | * | E | N | R | F | * | * | N | * | * | * |
| *Pundamilia nyererei* | * | E | * | * | R | K | * | S | * | * | * | E | K | * | D | N | K | * | * | * |
| *Mesitornis unicolor* | * | M | * | E | E | M | R | * | V | * | E | N | S | * | * | * | N | * | * | * |
| *Gallus gallus* | E | * | * | A | E | V | R | * | * | * | E | N | R | F | * | * | N | * | * | * |
| *Lates calcarifer* | * | E | * | Q | R | K | * | T | E | * | * | E | K | * | * | N | E | * | * | * |
| *Chelonoidis abingdonii* | E | N | * | S | Q | V | R | * | * | * | A | N | K | F | * | * | K | * | * | * |
| *Arvicanthis niloticus* | K | * | * | * | * | R | * | * | * | H | * | T | N | F | * | L | * | * | * | * |
| *Thamnophis elegans* | * | E | * | K | Q | A | R | D | * | * | A | N | * | * | * | * | K | * | * | * |
| *Oreochromis niloticus* | * | E | * | * | R | K | * | S | * | * | * | E | K | * | * | N | K | * | * | * |
| *Melopsittacus undulatus* | * | M | * | E | E | R | K | * | * | * | D | N | S | F | * | * | N | * | * | * |
| *Buceros rhinoceros silvestris* | E | N | * | E | Q | R | R | * | * | * | E | N | N | F | * | * | K | * | * | * |
| *Meleagris gallopavo* | E | * | * | A | E | V | R | * | * | * | E | N | R | F | * | * | N | * | * | * |
| *Tyto alba* | * | M | * | E | E | R | R | * | * | * | E | N | R | F | * | * | N | * | * | * |
| *Betta splendens* | * | D | * | E | * | E | * | T | Q | * | * | E | K | F | E | N | E | * | * | * |
| *Python bivittatus* | E | E | * | M | Q | V | R | D | * | * | D | N | K | F | * | * | K | * | * | * |
| *Ornithorhynchus anatinus* | E | Q | * | T | Q | K | Q | * | * | * | * | N | K | F | * | * | N | * | * | * |
| *Pangasianodon hypophthalmus* | R | E | * | Q | * | E | D | S | R | * | * | Q | A | F | * | N | E | * | * | * |
| *Pseudopodoces humilis* | E | E | * | E | E | R | R | * | N | * | E | N | N | F | * | * | N | * | * | * |
| *Zootoca vivipara* | E | D | * | M | Q | A | R | * | * | * | E | N | R | F | * | * | * | * | * | * |
| *Chaetura pelagica* | * | M | * | E | E | R | K | * | * | * | E | N | N | F | * | * | K | * | * | * |
| *Calypte anna* | * | M | * | E | E | R | R | * | * | H | E | N | S | F | * | * | * | * | * | * |
| *Pygocentrus nattereri* | R | E | * | Q | * | E | D | S | R | * | * | E | A | F | * | N | E | * | * | * |
| *Poecilia reticulata* | E | A | * | E | R | E | N | S | * | * | K | E | N | * | * | N | E | * | * | * |
| *Nipponia nippon* | * | M | * | E | E | R | R | * | N | * | E | N | N | F | * | * | N | * | * | * |
| *Athene cunicularia* | * | M | * | E | E | R | R | * | * | * | E | N | S | F | * | * | N | * | * | * |
| *Chlamydotis macqueenii* | * | M | * | E | E | R | R | * | N | * | E | N | S | F | * | * | N | * | * | * |
| *Microcaecilia unicolor* | E | R | * | S | Q | R | L | * | * | N | E | N | R | * | * | * | N | * | * | * |
| *Pygoscelis adeliae* | * | M | * | E | E | K | R | * | N | * | E | N | S | F | * | * | N | * | * | * |
| *Notothenia coriiceps* | * | E | * | Q | * | E | * | T | Q | * | * | E | K | * | K | N | E | * | * | * |
| *Parus major* | E | E | * | E | E | M | R | * | N | * | E | N | N | F | * | * | N | * | * | * |
| *Mastacembelus armatus* | E | N | * | Q | S | E | * | T | E | * | * | E | K | F | * | N | K | * | * | * |
| *Zonotrichia albicollis* | * | M | * | V | E | R | R | * | * | * | E | N | K | F | * | * | N | * | * | * |
| *Fulmarus glacialis* | * | M | * | Q | E | S | S | * | S | * | E | N | S | F | * | * | N | * | * | * |
| *Gavia stellata* | * | M | * | E | E | K | R | * | N | * | E | N | S | F | * | * | N | * | * | * |
| *Astatotilapia calliptera* | * | E | * | * | R | K | * | S | * | * | * | E | K | * | D | N | K | * | * | * |
| *Numida meleagris* | E | I | * | A | E | V | R | * | * | * | E | N | R | F | * | * | N | * | * | * |
| *Taeniopygia guttata* | * | I | * | E | E | R | R | * | N | * | E | N | T | F | * | * | N | * | * | * |
| *Sorex araneus* | N | K | * | E | N | K | D | * | * | * | N | I | T | F | * | * | N | * | * | * |
| *Sinocyclocheilus rhinocerous* | S | E | * | K | * | E | * | T | N | * | * | E | A | * | * | N | E | * | * | * |
| *Poecilia mexicana* | E | A | * | E | R | E | N | S | * | * | K | E | N | * | * | N | E | * | * | * |
| *Pseudonaja textilis* | M | E | * | K | Q | I | R | V | * | * | N | N | I | F | * | * | E | * | * | * |
| *Salarias fasciatus* | R | E | * | * | Q | A | S | G | * | * | * | E | S | F | * | N | K | * | * | * |
| *Pelodiscus sinensis* | E | N | * | S | E | V | Q | * | * | * | A | N | K | * | * | * | K | * | * | * |
| *Perca flavescens* | K | E | * | Q | * | E | * | T | R | * | * | E | K | F | * | N | E | * | * | * |
| *Merops nubicus* | * | M | * | E | E | W | R | * | S | * | E | N | - | - | * | * | N | * | * | * |
| *Seriola dumerili* | K | E | * | Q | R | E | * | T | A | * | * | E | K | F | D | N | E | * | * | * |
| *Thamnophis sirtalis* | * | E | * | K | Q | A | R | D | * | * | A | N | * | * | * | * | K | * | * | * |
| *Phaethon lepturus* | * | M | * | E | E | R | R | * | * | * | E | N | S | F | * | * | N | * | * | * |
| *Parambassis ranga* | D | K | * | Q | N | K | * | T | E | * | * | E | K | F | * | N | E | * | * | * |
| *Neolamprologus brichardi* | * | E | * | * | R | K | * | S | * | * | * | E | K | * | H | N | K | * | * | * |
| *Vombatus ursinus* | R | E | * | E | T | K | * | * | E | * | * | I | T | F | * | * | * | * | * | * |
| *Trachemys scripta elegans* | E | N | * | S | Q | V | R | * | * | * | A | N | K | * | * | * | K | * | * | * |
| *Apteryx rowi* | * | M | * | T | E | I | K | * | * | * | E | N | N | F | Y | * | K | * | * | * |
| *Poecilia formosa* | E | A | * | E | R | E | N | S | * | * | K | E | N | * | * | N | E | * | * | * |
| *Poecilia latipinna* | E | A | * | E | R | E | N | S | * | * | K | E | N | * | * | N | E | * | * | * |
| *Carassius auratus* | S | E | * | K | * | E | * | T | N | * | * | E | A | * | * | N | E | * | * | * |
| *Stegastes partitus* | * | E | * | Q | * | I | D | T | A | * | * | E | R | F | * | N | N | * | * | * |
| *Denticeps clupeoides* | R | E | * | R | * | E | N | T | * | * | * | E | N | F | * | N | E | * | * | * |
| *Gopherus evgoodei* | E | N | * | S | Q | V | R | * | * | * | A | N | K | F | * | * | K | * | * | * |
| *Protobothrops mucrosquamatus* | * | E | * | K | Q | A | R | D | * | * | N | N | * | F | * | * | K | * | * | * |
| *Seriola lalandi dorsalis* | K | E | * | Q | R | E | * | T | A | * | * | E | K | F | D | N | E | * | * | * |
| *Xenopus tropicalis* | * | D | * | K | R | Q | * | * | V | H | * | N | A | F | * | M | N | * | * | * |
| *Charadrius vociferus* | * | M | * | Q | E | R | R | * | N | * | E | N | N | F | * | * | * | * | * | * |
| *Anolis carolinensis* | * | E | * | L | Q | I | N | * | N | * | E | R | T | F | * | * | N | * | * | K |
| *Lonchura striata domestica* | * | M | * | E | E | R | R | * | N | * | E | N | T | F | * | * | N | * | * | * |
| *Corapipo altera* | * | N | * | E | Q | R | M | * | N | * | E | N | S | F | * | L | N | * | * | * |
| *Esox lucius* | S | E | * | K | W | R | * | T | * | * | K | E | S | F | * | N | Q | * | * | * |
| *Maylandia zebra* | * | E | * | * | R | K | * | S | * | * | * | E | K | * | D | N | K | * | * | * |
| *Neopelma chrysocephalum* | * | N | * | E | Q | R | M | * | * | * | E | N | S | F | * | * | N | * | * | * |
| *Pogona vitticeps* | * | Q | * | M | E | A | Q | * | N | * | E | T | E | F | * | * | N | * | * | * |
| *Catharus ustulatus* | E | K | * | E | E | S | R | * | N | * | E | N | K | F | * | N | N | * | * | * |
| *Haliaeetus albicilla* | * | M | * | E | E | R | R | * | N | * | E | - | S | F | * | * | N | * | * | * |
| *Corvus brachyrhynchos* | * | M | * | E | E | R | R | * | N | * | E | - | - | - | * | * | N | * | * | * |
| *Haplochromis burtoni* | * | E | * | * | R | K | * | S | * | * | * | E | K | * | D | N | K | * | * | * |
| *Manacus vitellinus* | * | N | * | E | Q | R | M | * | N | * | E | N | S | F | * | N | N | * | * | * |
| *Cynoglossus semilaevis* | * | E | * | E | * | E | * | S | A | * | N | E | K | F | * | N | M | * | * | * |
| *Empidonax traillii* | * | Q | * | E | Q | R | M | * | * | * | E | N | G | F | * | * | N | * | * | * |
| *Archocentrus centrarchus* | * | Q | * | E | * | K | * | S | E | * | * | E | K | * | D | N | K | * | * | * |
| *Amphiprion ocellaris* | * | E | * | K | N | T | * | T | V | * | * | E | K | F | * | N | E | * | * | * |

| | | | | | | | | | | | | | | | | | | | | |
|---|---|---|---|---|---|---|---|---|---|---|---|---|---|---|---|---|---|---|---|---|
| *Hippocampus comes* | E | D | * | R | * | E | * | S | E | * | * | Q | K | F | * | N | K | * | * | * |
| *Myripristis murdjan* | R | V | * | Q | R | E | * | T | V | * | * | E | N | * | * | N | E | * | * | * |
| *Corvus cornix cornix* | * | M | * | E | E | R | R | * | N | * | E | N | S | F | * | R | N | * | * | * |
| *Amblyraja radiata* | E | E | * | A | T | E | D | * | Q | * | K | N | K | F | Y | N | L | * | * | * |
| *Lepidothrix coronata* | * | N | * | E | Q | R | M | * | N | * | E | N | S | F | * | N | N | * | * | * |
| *Alligator mississippiensis* | - | * | * | N | Q | Q | N | * | G | * | E | N | K | * | * | M | K | * | * | * |
| *Alligator sinensis* | - | * | * | N | Q | Q | N | * | G | * | E | N | K | * | * | M | K | * | * | * |
| *Nothobranchius furzeri* | * | D | * | Q | R | Q | N | T | A | * | * | E | K | * | * | N | E | * | * | * |
| *Sander lucioperca* | K | E | * | Q | * | E | * | T | R | * | * | E | K | F | * | N | E | * | * | * |
| *Crocodylus porosus* | - | V | * | N | Q | Q | D | * | G | * | E | N | R | * | * | N | K | * | * | * |
| *Erinaceus europaeus* | E | K | * | * | D | R | Q | * | N | * | E | T | N | * | * | N | * | * | * | * |
| *Pipra filicauda* | * | N | * | E | Q | R | M | * | N | * | E | N | S | F | * | N | N | * | * | * |
| *Sinocyclocheilus anshuiensis* | S | E | * | K | * | E | * | T | N | * | * | E | A | * | * | N | E | * | * | * |
| *Gavialis gangeticus* | - | * | * | N | Q | Q | D | * | G | * | E | N | R | * | * | N | K | * | * | * |
| *Xiphophorus hellerii* | E | A | * | E | R | Q | N | S | E | * | K | E | N | * | * | N | E | * | * | * |
| *Strigops habroptila* | * | M | * | E | E | R | R | * | N | * | E | N | T | F | * | * | N | * | * | * |
| *Acanthochromis polyacanthus* | R | E | * | K | D | T | N | T | E | * | * | E | Q | F | * | N | E | * | * | * |
| *Nanorana parkeri* | D | V | * | Q | E | N | V | * | Q | H | * | F | R | * | K | L | D | * | * | * |
| *Erpetoichthys calabaricus* | K | Q | * | N | D | I | * | P | I | * | * | N | D | F | * | N | K | * | * | * |
| *Oreochromis aureus* | * | E | * | * | R | K | * | S | * | * | * | E | K | * | D | N | K | * | * | * |
| *Kryptolebias marmoratus* | E | D | * | L | * | E | N | T | A | * | * | E | K | * | * | N | E | * | * | * |
| *Etheostoma spectabile* | K | E | * | Q | M | E | * | T | Q | * | * | E | K | F | * | N | E | * | * | * |
| *Anarrhichthys ocellatus* | K | V | * | K | M | K | * | T | Q | * | * | E | Q | F | * | N | E | * | * | * |
| *Callorhinchus milii* | E | A | * | K | E | T | K | Q | * | * | K | D | K | F | * | N | V | * | * | * |
| *Cyprinodon variegatus* | E | E | * | K | R | Q | N | T | * | * | * | E | K | * | * | N | E | * | * | * |
| *Electrophorus electricus* | R | E | * | K | * | E | D | T | R | * | * | E | A | F | * | N | E | * | * | * |
| *Chanos chanos* | K | E | * | R | * | E | N | T | H | * | * | Q | E | F | * | N | E | * | * | * |
| *Chiroxiphia lanceolata* | * | N | * | E | Q | R | M | * | S | * | E | N | S | F | * | L | N | * | * | * |
| *Anabas testudineus* | S | E | * | Q | R | E | * | T | Q | * | * | E | K | F | * | N | E | * | * | * |
| *Scleropages formosus* | K | E | * | Q | S | E | N | T | T | * | * | E | S | F | * | N | E | * | * | * |
| *Latimeria chalumnae* | * | E | * | K | D | Q | N | Q | E | * | * | N | Q | F | Y | N | K | * | * | * |
| *Oncorhynchus mykiss* | R | E | * | * | Q | G | N | T | H | * | * | E | K | F | D | N | E | * | * | * |
| *Oryzias melastigma* | * | E | * | K | G | E | Q | T | E | * | * | E | K | * | * | N | E | * | * | * |
| *Eurypyga helias* | * | M | * | N | E | S | R | * | N | * | E | N | K | F | K | N | K | * | * | * |
| *Boleophthalmus pectinirostris* | R | E | * | K | * | E | Q | S | A | * | N | E | N | F | E | N | K | * | * | * |
| *Gouania willdenowi* | R | E | * | Q | R | I | N | S | A | * | * | E | N | F | * | N | E | * | * | * |
| *Haliaeetus leucocephalus* | - | M | * | E | E | R | R | * | N | * | E | N | S | F | * | * | N | * | * | * |
| *Tachysurus fulvidraco* | R | E | * | K | * | E | D | S | R | * | * | E | A | F | * | N | E | * | * | * |
| *Labrus bergylta* | K | E | * | K | R | E | * | S | V | * | N | E | K | * | * | N | E | * | * | * |
| *Cottoperca gobio* | R | V | * | Q | * | E | D | S | Q | * | * | E | K | * | K | N | E | * | * | * |
| *Takifugu rubripes* | * | Q | * | K | E | E | K | T | E | * | N | E | K | F | * | N | E | * | * | * |
| *Xiphophorus couchianus* | E | A | * | E | R | Q | N | S | E | * | K | E | N | * | * | N | E | * | * | * |
| *Astyanax mexicanus* | R | D | * | L | * | E | D | S | R | * | * | E | E | F | * | N | E | * | * | * |
| *Lepisosteus oculatus* | D | V | * | * | E | K | N | T | H | * | L | E | K | F | E | N | K | * | * | * |
| *Echeneis naucrates* | K | E | * | K | R | E | * | T | K | * | N | E | N | * | * | N | E | * | * | * |
| *Paralichthys olivaceus* | K | E | * | L | R | Q | * | T | Q | * | N | K | E | F | * | N | E | * | * | * |
| *Xiphophorus maculatus* | E | A | * | E | R | Q | N | S | E | * | K | E | N | * | * | N | E | * | * | * |
| *Clupea harengus* | R | A | * | E | R | V | K | T | E | * | * | E | T | F | D | N | K | * | * | * |
| *Gadus morhua* | R | A | * | E | * | T | * | S | V | * | D | E | A | F | * | N | K | * | * | * |

### Supporting Figures

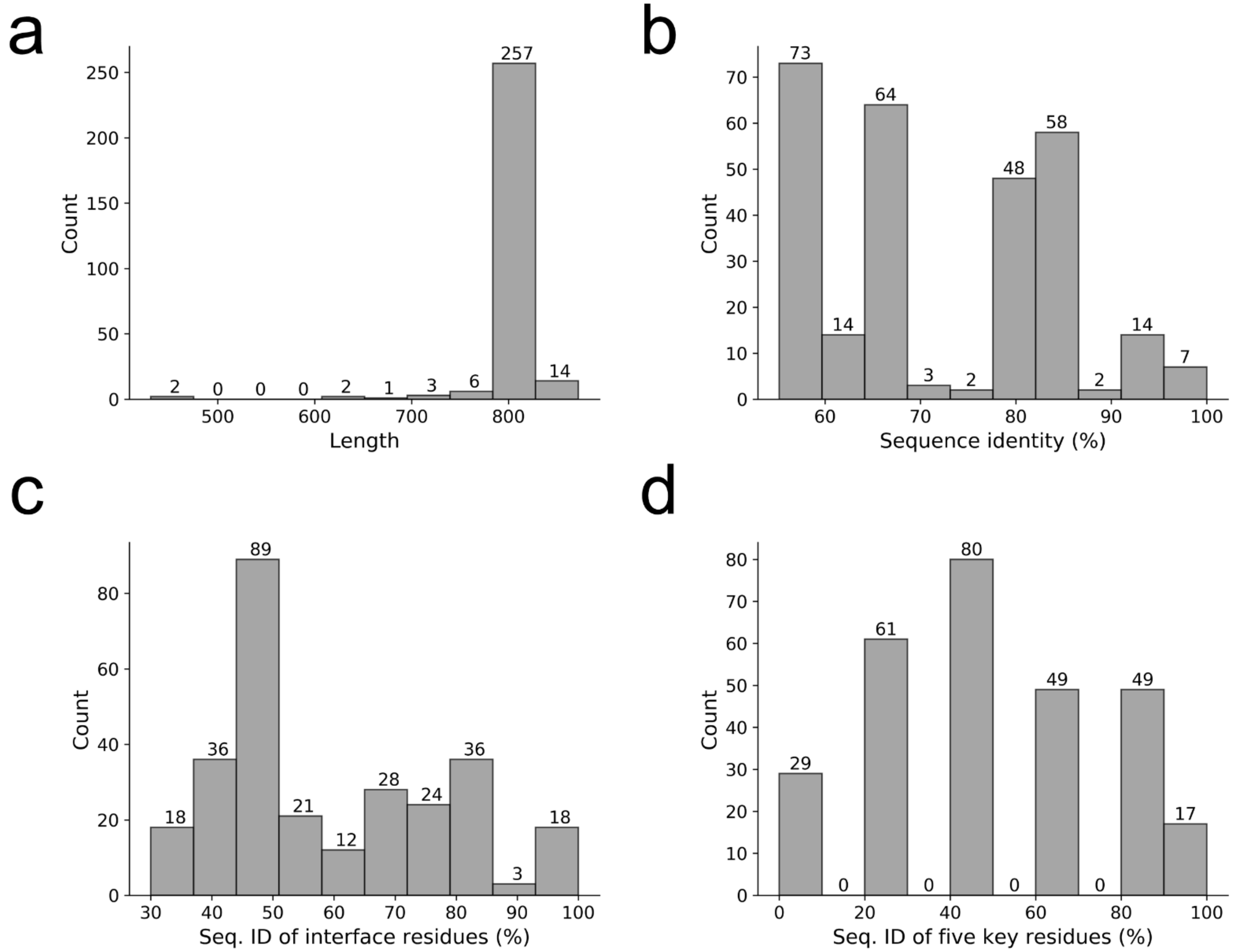


**Fig. S1**. The distribution of protein length, and sequence identity for all, interface, and the five key residues for 285 ACE2 orthologs.

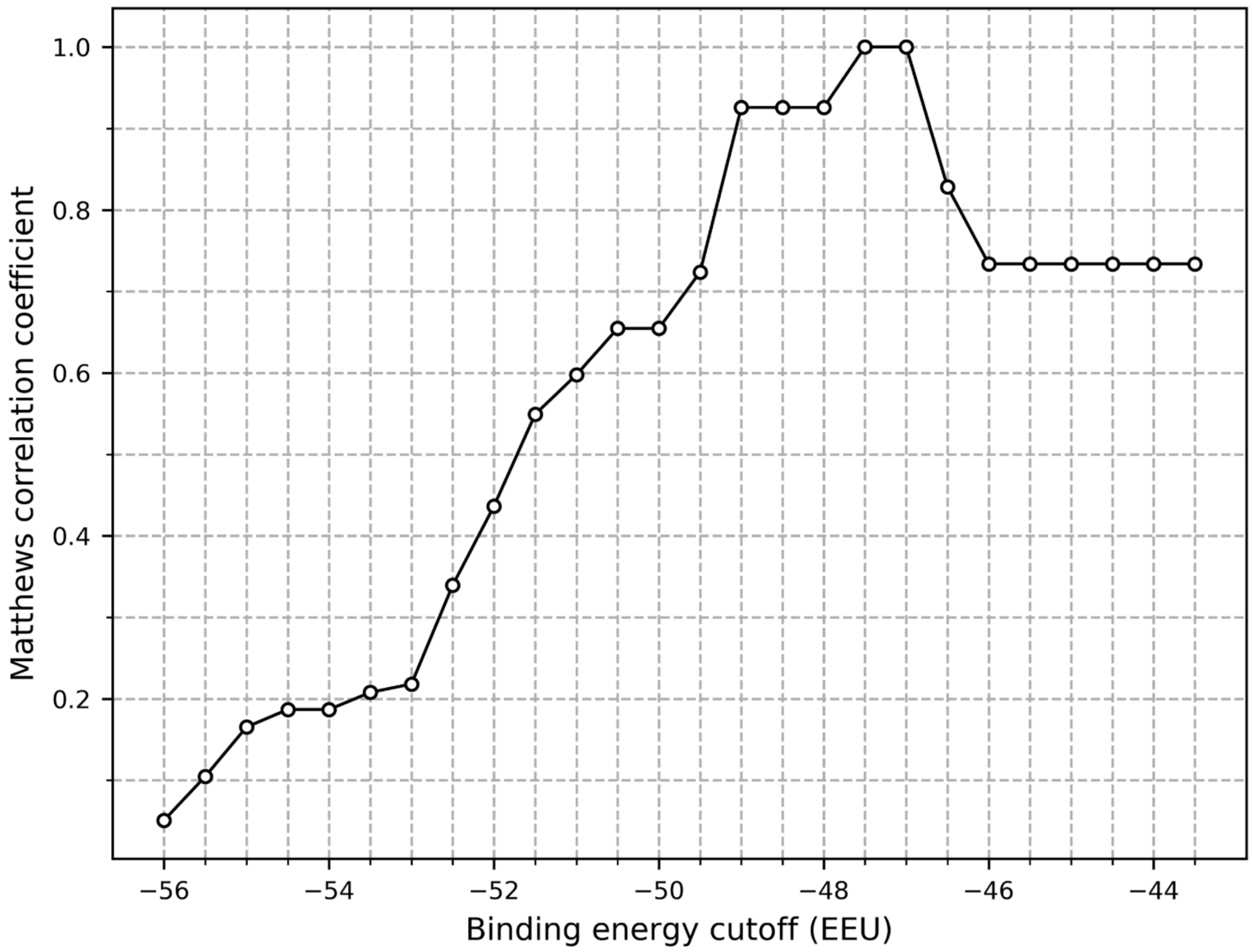


**Fig. S2**. Matthews correlation coefficient for classifying experimentally determined effective ACE2 receptors from the less effective ones by the binding energy calculated from 500 models.

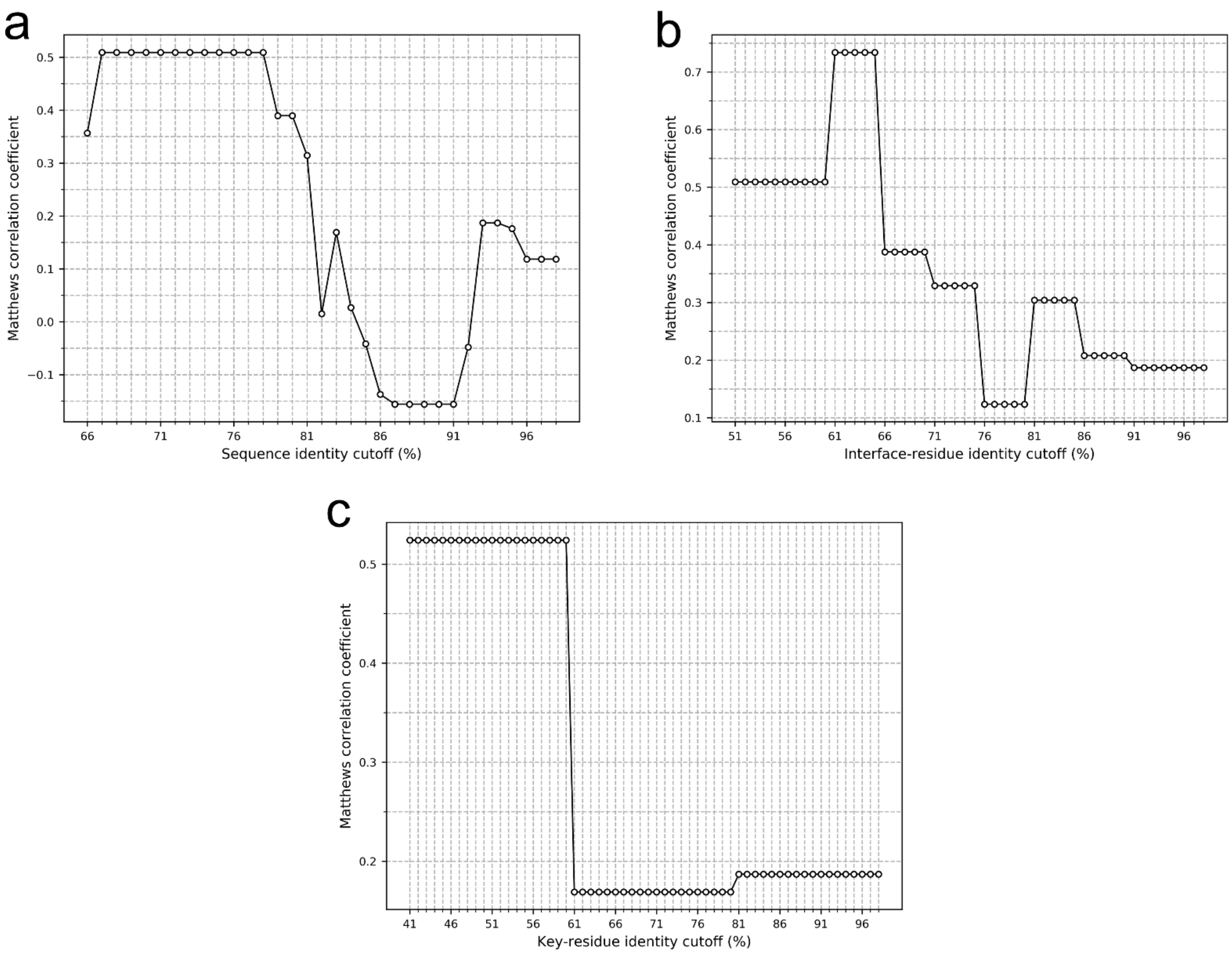


**Fig. S3**. Matthews correlation coefficient for classifying experimentally determined effective ACE2 receptors from the less effective ones by sequence identity in terms of all (a), interface (b), and key (c) residues.

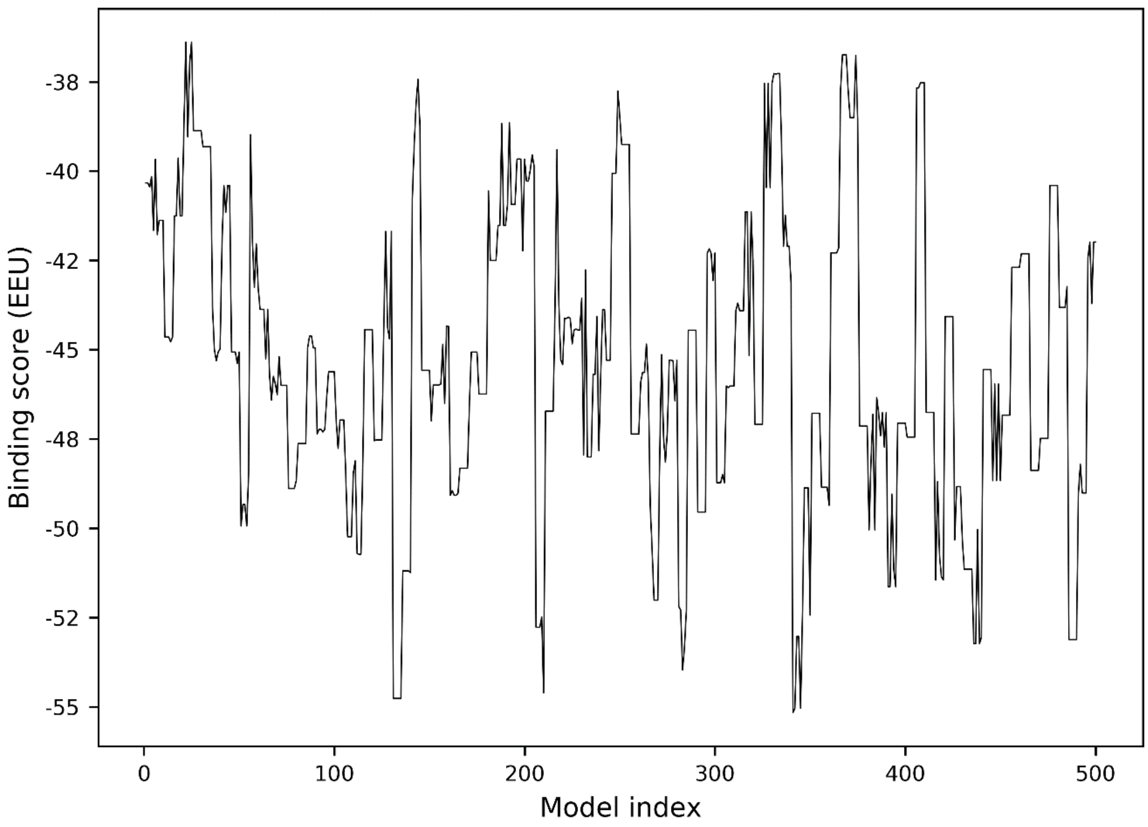


**Fig. S4**. Binding score of 500 models for the hACE2/S-RBD complex.

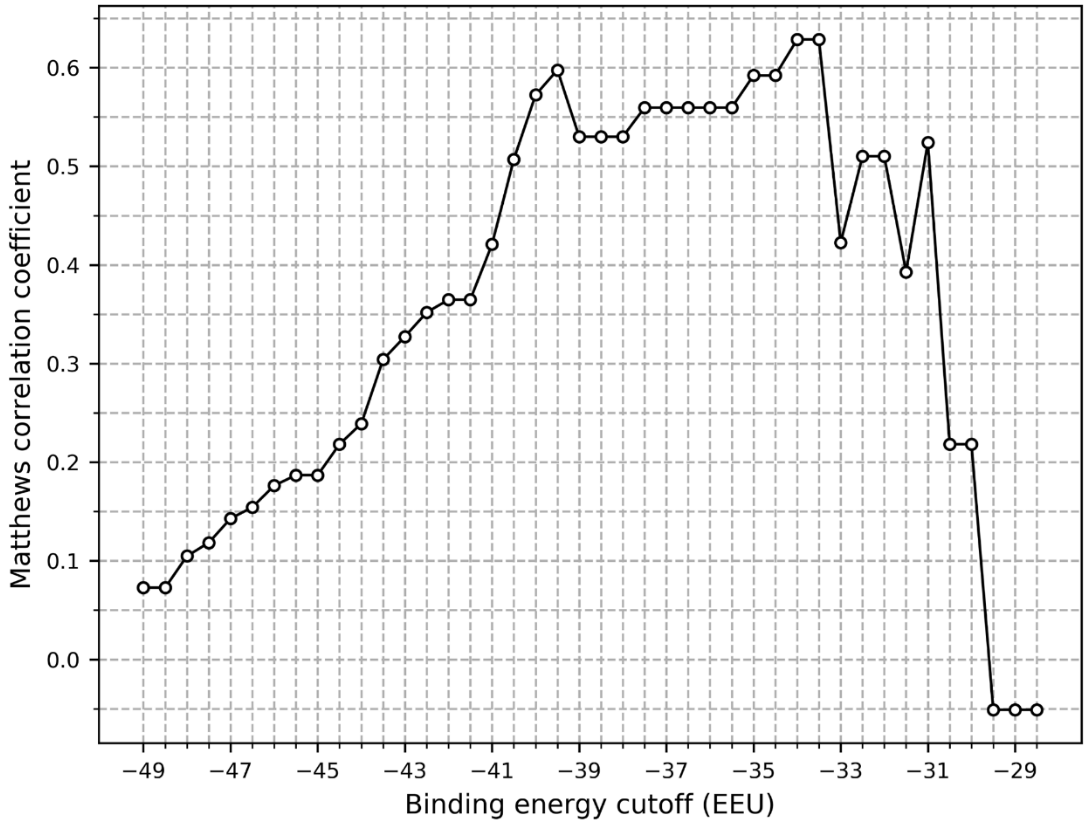


**Fig. S5**. Matthews correlation coefficient for classifying experimentally determined effective ACE2 receptors from the less effective ones by the binding energy calculated from the first model.
